# A chromosome-level, haplotype-resolved genome assembly and annotation for the Eurasian minnow (Leuciscidae: *Phoxinus phoxinus*) provide evidence of haplotype diversity

**DOI:** 10.1101/2023.11.30.569369

**Authors:** Temitope Opeyemi Oriowo, Ioannis Chrysostomakis, Sebastian Martin, Sandra Kukowka, Thomas Brown, Sylke Winkler, Eugene W Myers, Astrid Böhne, Madlen Stange

## Abstract

In this study we present an in-depth analysis of the Eurasian minnow (*Phoxinus phoxinus*) genome, highlighting its genetic diversity, structural variations, and evolutionary adaptations. We generated an annotated haplotype-phased, chromosome-level genome assembly (2n = 50) by integrating high-fidelity (HiFi) long reads and chromosome conformation capture data (Hi-C). We achieved a haploid size of 940 Megabase pairs (Mbp) for haplome one and 929 Mbp for haplome two with high scaffold N50 values of 36.4 Mb and 36.6 Mb and BUSCO scores of 96.9% and 97.2%, respectively, indicating a highly complete genome assembly.

We detected notable heterozygosity (1.43%) and a high repeat content (approximately 54%), primarily consisting of DNA transposons, which contribute to genome rearrangements and variations. We found substantial structural variations within the genome, including insertions, deletions, inversions, and translocations. These variations affect genes enriched in functions such as dephosphorylation, developmental pigmentation, phagocytosis, immunity, and stress response.

In the annotation of protein-coding genes, 30,980 mRNAs and 23,497 protein-coding genes were identified with a high completeness score, which further underpins the high contiguity of our genome assemblies. We performed a gene family evolution analysis by comparing our proteome to ten other teleost species, which identified immune system gene families that prioritise histone-based disease prevention over NLR-based immune responses.

Additionally, demographic analysis indicates historical fluctuations in the effective population size of *P. phoxinus*, likely correlating with past climatic changes.

This annotated, phased reference genome provides a crucial resource for resolving the taxonomic complexity within the genus *Phoxinus* and highlights the importance of haplotype-phased assemblies in understanding haplotype diversity in species characterised by high heterozygosity.

## Introduction

Species of the genus *Phoxinus* show considerable morphologic and genetic differences (Kottelat & Freyhof, 2007). For a long time, *Phoxinus phoxinus* (Linnaeus, 1758) — as its common name Eurasian minnow suggests — was thought to be a widespread species throughout Europe. However, the taxonomy of the genus was revealed to be more complex (Kottelat, 2007; Kottelat & Freyhof, 2007) including substantial cryptic species diversity (Palandačić et al., 2015, 2017, 2020). At present, 22 genetic *Phoxinus* lineages have been distinguished, comprising at least 13 recognised species (Palandačić et al., 2020). The neotype locality for *P. phoxinus* was placed on the Agger stream (Kottelat, 2007), a tributary of the river Sieg in Germany that feeds into the Rhine River. The designation of a neotype locality is used when the holotype material and sources used to describe a species are missing. Prior to 2012, minnows were a rare occurrence in that area (Landesamt für Natur, Umwelt und Verbraucherschutz Nordrhein-Westfalen, 2020). Taxonomically problematic is the fact that the connection between the species name *P. phoxinus* and its assigned lineage “10” (Knebelsberger et al., 2015) is uncertain. This is due to the lack of genetic analysis of the neotype (NRM-55108) designated by Kottelat, (2007), which comes from an area with multiple lineages present (5b, 10, 12; defined by a combination of morphological and single-gene markers) (Palandačić et al., 2020, 2022). Several studies and researchers are currently dealing with resolving the taxonomic and phylogenetic conundrum this genus poses (Bogutskaya et al., 2023; Corral-Lou et al., 2019; Denys et al., 2020; Palandačić et al., 2017, 2020; Reier et al., 2022; Vucić et al., 2018). Until now, these studies suffered from the lack of a high-quality reference genome, which would make it possible to go beyond the analysis of single-gene markers.

Due to the previously unrecognised species diversity, most of what is known about the natural history and behaviour of Eurasian minnows is attributed to *P. phoxinus* or its synonyms. Certain presumptions about minnows are assumed to be universally applicable for the entire genus. In the following, we will refer to Eurasian minnows including all cryptic species in the *Phoxinus* genus. Eurasian minnows are small, schooling freshwater fishes that occupy oxygen-rich, cold streams of flowing waters, lake-shores with wave-movements, or highland lakes (Frost, 1943; Mills, 1987, 1988). They assume a complex role in the river or lake ecosystem as they feed on algae, plant debris, molluscs, crustaceans, and insects (Frost, 1943). *Phoxinus* species interact with smaller animals such as water snails and insect larvae, but also larger fishes such as co-occurring salmonids (Museth et al., 2010). However, seemingly contradictory behaviours have been reported as well. According to Frost (1943), *Phoxinus* spp. spawn over clean gravel and shallow, flowing waters. Yet, populations from the Baltic Sea reportedly reside and spawn in brackish habitats (Svirgsden et al., 2018). While Eurasian minnows are sometimes reported to be sensitive to pollution (Bagge & Hakkari, 1992), they are also tolerant of polluted waters (King et al., 2011). They might even benefit from increased primary production in eutrophic waters (King et al., 2011; Moiseenko et al., 2009). In terms of dispersal and migration behaviour, Eurasian minnows from the Volga were less frequently found in migratory habitats (Pavlov & Mikheev, 2017), or demonstrated a tendency for downstream movement in a mesocosm experiment conducted with specimens from Wales (Jones et al., 2021). In contrast, Eurasian minnows were found to migrate upstream during the breeding season (Frost, 1943; Svirgsden et al., 2018) as well as through fishways (Jansen et al., 1999) in populations from England, Estonia, and Germany, respectively. The aforementioned reported differences or the seemingly broad occurrence pattern could be due to the fact that studies reporting on *P. phoxinus* or “the Eurasian minnow” were conducted on different *Phoxinus* species.

A crucial step towards informed and state-of-the-art biodiversity, phylogenetic, and comparative genomic research is the generation of haplotype-resolved, chromosome-level reference genomes. Phasing of haplomes, the haplotype-resolved elements in reference genome assemblies, is now facilitated with the latest development of chromosome conformation capture sequencing, namely Hi-C (Belton et al., 2012; Lin et al., 2018) and its integration into assembly pipelines (H. Cheng et al., 2021). Resultantly, the number of studies acknowledging genetic variation between haplomes is rapidly increasing, e.g. in Cavendish banana (Huang et al., 2023), ginger (S.-P. Cheng et al., 2021), carnation (L. Jiang et al., 2022), African cassava (Mansfeld et al., 2021), Australian lime (Nakandala et al., 2023), carnation (H. Jiang et al., 2023), cattle (Low et al., 2020), pearl oyster (Takeuchi et al., 2022), turkey (Barros et al., 2023), and ocellated puffer fish (Zeng et al., 2023).

We conducted comparisons between the two *Phoxinus* haplomes to identify both sequence and structural variations present in an individual *P. phoxinus* genome. When examining sequence variation, heterozygosity, polymorphism, and allelic diversity were considered (Carley et al., 2022; DeWoody et al., 2021). Structural variation can be examined within and among genomes by investigating the copy number, orientation and location of coding and non-coding sequences on the chromosome (Wellenreuther et al., 2019). Copy number variation comprises deletions, insertions, and duplications, while orientational variation encompasses inversions. Local variation can be attributed to translocations. Structural variation at the genome level has been associated with variation in both phenotypes and complex traits (Mérot et al., 2020; Weischenfeldt et al., 2013).

Ultimately, this reference genome has two major implications. Firstly, it should provide the *Phoxinus* research community with the necessary basis for the phylogenetic unravelling of this complex genus. Secondly, this haplotype-resolved, chromosome-level reference genome demonstrates once more that genetic diversity contains an individual layer, namely, the intra-individual diversity on the haplome level that needs to find attention in the scientific community.

## Methods

### Specimen Collection and Sampled Tissues

Two male *P. phoxinus* specimens from the Museum Koenig Bonn live exhibition, originally from the river Sieg close to the neotype locality, North-Rhine Westphalia, Germany, were sacrificed using Ethyl 3-aminobenzoate methanesulfonate salt (MS-222; Sigma-Aldrich) under permission per § 11 Abs. 1 Nr. 1 b, Sa and 8d Tierschutzgesetz (TierSchG) issued by the Amt für Umwelt, Verbraucherschutz und Lokale Agenda, Lebensmittelüberwachung und Veterinärdienste, under Zeichen 56.2. Tissues from fin, brain, liver, spleen, muscle, heart, skin, and testis were immediately sampled and flash-frozen in liquid nitrogen until use. A proxy specimen voucher from the neotype locality (River Agger, GPS: 50.841358, 7.201399) sampled on 11 May 2022 is available at the ichthyological collection, Museum Koenig Bonn, under ICH-132418 for the body and at the LIB Biobank, Museum Koenig Bonn, under accession number ZFMK-TIS-64936 for fin tissue, while additional fin tissue from the sampled individuals are deposited for future reference at LIB Biobank under accession numbers ZFMK-TIS-60719 and ZFMK-TIS-60720. Tissues (brain, liver, spleen, muscle, heart, gonad) from individual ZFMK-TIS-60720 were used for long read libraries to generate the genome assembly; tissues (brain, gill, gonad, liver, muscle, skin, spleen) from ZFMK-TIS-60719 were used for RNA-sequencing and subsequent genome annotation.

### Extraction of High Molecular Weight Genomic DNA

High molecular weight genomic DNA (HMW gDNA) of *P. phoxinus* was extracted with the circulomics Nanobind Tissue Big DNA kit (part number NB-900-701-01, protocol version Nanobind Tissue Big DNA Kit Handbook v1.0 (11/19)) according to the manufacturer’s instructions. In brief, spleen tissue was minced into small slices on a clean and cold surface. Tissues were finally homogenised with the TissueRuptor II device (Qiagen) making use of its maximal settings. After complete tissue lysis, the remaining cell debris were removed, and the gDNA was bound to circulomics Nanobind discs in the presence of isopropanol. HMW gDNA was eluted from the nanobind discs in EB. The integrity of the HMW gDNA was determined by pulse field gel electrophoresis using the Pippin Pulse^TM^ device (SAGE Science) and the Agilent Femtopulse. The majority of the gDNA was between 20 and more than 400 kilobases (kb) in length. All pipetting steps of long gDNA have been done very carefully with wide-bore pipette tips. HMW gDNA was further purified after extraction with 1x AMPure beads.

### PacBio HiFi Library Preparation and Sequencing

A long insert HiFi library was prepared as recommended by Pacific Biosciences according to the “Guidelines for preparing HiFi SMRTbell libraries using the SMRTbell Express Template Prep Kit 2.0” (PN 101-853-100, version 03). In summary, 10 µg of purified HMW gDNA was sheared twice, aiming for 25 and 20 kb fragments, respectively, with the MegaRuptor^TM^ device (Diagenode). The sheared gDNA was enriched with 1x AMPure beads according to the manufacturer’s instructions. 6 µg sheared gDNA have been used for Pacbio HiFi library preparation. The PacBio SMRTbell^TM^ library was size-selected for fragments larger than 6 kb with the BluePippin^TM^ device according to the manufacturer’s instructions. The final HiFi library had a fragment size of 12.8 kb on the Agilent Fragment Analyzer on the High sensitivity 50 kb large fragment kit.

60 pM of the size selected library were run on two Sequel II SMRT cells with the SEQUEL II sequencing kit 2.2 for 30 hours on the SEQUEL II; pre-extension time was 2 hours. PacBio CCS reads (read quality > 0.99) were created from the subreads.bam files using PacBio’s ccs v.6.3.0 (https://github.com/PacificBiosciences/ccs.git) command line tool and further refined by applying the tool DeepConsensus v0.2 (Baid et al., 2023), on PacBio reads within 98.0-99.5% read accuracy (Baid et al., 2023). Finally, reads containing the PacBio adapter sequence were filtered out by applying a Blastn search (v2.9) (Altschul et al., 1990) providing the PacBio adapter sequence and the following arguments: “reward 1 -penalty -5 -gapopen 3 -gapextend 3 -dust no -soft_masking false -evalue 700 -searchsp 1750000000000-outfmt 7”. This resulted in a total yield of 37,3 Gb data with a mean read length of 10.7 kb and a read N50 of 10.2 kb.

### Chromatin Conformation Capturing Library Preparation and Sequencing

Chromatin conformation capturing was done making use of the ARIMA HiC+ Kit (Material Nr. A410110) and followed the user guide for animal tissues (ARIMA-HiC 2.0 kit Document Nr: A160162 v00). In brief, 40 mg flash-frozen powdered muscle tissue was cross-linked chemically. The cross-linked chromatin was digested with a restriction enzyme cocktail consisting of four restriction enzymes. The 5’-overhangs were filled in and labelled with biotin. Spatially proximal digested DNA ends were ligated, and, finally, the ligated biotin-containing fragments were enriched and went for Illumina library preparation, which followed the ARIMA user guide for Library preparation using the Kapa Hyper Prep kit (ARIMA Document Part Number A160139 v00). The barcoded HiC libraries run on an S4 v1.5 XP flow cell of a NovaSeq6000 with 300 cycles.

### RNA Extraction and Sequencing

RNA was extracted from flash-frozen brain, gill, gonad, liver, muscle, skin, and spleen (specimen identifier ZFMK-TIS-60719) tissues. All tissues except for spleen and liver were extracted using the RNeasy Mini Kit (Qiagen, Cat. No. 74104) suitable for tissue amounts of 0.5 to 30 mg. Spleen and liver were extracted using QIAamp DNA Micro Kit (Qiagen) suitable for starting material of less than 5 mg. All tissues were homogenised using PowerBead Tubes Ceramic (Qiagen, diameter 2.8mm; Cat. No. 13114-50) on a PowerLyzer™ 24 Bench Top Bead-Based Homogenizer (MO BIO Laboratories, Inc). Before use, 180 µL 1 M dithiothreitol (DTT) was added to 4.5 ml lysis buffer RLT. 350 µL prepared RLT buffer was added to each tissue in a PowerBead Tube (Qiagen) and one cycle of 45 seconds at 3500 rpm and another for 45 seconds at 4200 rpm on the PowerLyzer™ Homogenizer with a 5-minute pause at -20°C were performed. After homogenisation, the solution was centrifuged for 1 minute at 14,800 rpm. The lysate including tissue debris was transferred into a fresh 1.5 mL tube (LoBind®) and again centrifuged at 14,800 rpm for 3 minutes at room temperature (RT). Finally, the lysate was transferred to a gDNA Eliminator column (provided in the QIAamp Kits) and centrifuged at 10,000 rpm for 30 sec at RT. Washing and elution steps followed the standard protocol of the Micro and Mini Kit, respectively. Samples were eluted into 14 µL (spleen, liver) and 30 µL (all other tissues) RNAse-free water during 1 minute at 10,000 rpm centrifugation. RNA library preparation and sequencing were performed at Biomarker Technologies (BMKGene) GmbH on an llumina Novaseq 6000 S4 chip.

### K-mer Based Genome Size and Heterozygosity Estimation

To estimate the genome size and heterozygosity of the genome, a k-mer-based approach was used. K-mers were counted in the HiFi reads using Jellyfish v2.3.0 (Marçais & Kingsford, 2011) with parameters count *-C -m*, with *-C* indicating that the input reads from both the forward and reverse strand are being used, and *-m* specifying a k-mer length; we specified k-mer lengths of 19, 23, 25 and 30. The resulting k-mer counts were exported into a k-mer count histogram using the *histo* command of Jellyfish for subsequent genome size and heterozygosity estimation in GenomeScope v2.0 (Ranallo-Benavidez et al., 2020).

### *De novo* Genome Assembly and Scaffolding

Initial contigs for two representative haplomes were generated using Hifiasm v0.16.1-r375 (H. Cheng et al., 2021) with parameters --h1 --h2 -l2. Next, purge-dups v1.2.3 (Guan et al., 2020) was run on each produced haplotype assembly. The purging parameter “-l2” (purge all haplotigs at a similarity threshold of 75%) was chosen over the default “-l3” (purge all haplotigs at a similarity threshold of 55%) based on BUSCO and contiguity scores (Table 1). Assembly completeness for each produced haplotype assembly was assessed using BUSCO v5.4.7 (Simão et al., 2015) run in genome mode with the actinopterygii_odb10 database.

**Table 1:** Assembly statistics for haplomes 1 and 2 (Hap1, Hap2) from a parameter sweep using differing purging parameters in Hifiasm, followed by the use of purge-dups. BUSCO scores shown are for Complete (C), Single Copy (S), Duplicated (D), Fragmented (F), and Missing (M) genes.

| Assembly | BUSCO<br>C | BUSCO<br>S | BUSCO<br>D | BUSCO<br>F | BUSCO<br>M | Assembly size<br>(bp) | No.<br>contigs | Contig<br>N50 (bp) |
| --- | --- | --- | --- | --- | --- | --- | --- | --- |
| **hifiasm -**<br>**l2 Hap1** | 3529 | 3461 | 68 | 31 | 80 | 971,720,047 | 1,130 | 4,025,542 |
| **+purge-**<br>**dups** | 3516 | 3476 | 40 | 32 | 92 | 939,827,819 | 556 | 4,091,346 |
| **hifiasm -**<br>**l2 Hap2** | 3551 | 3479 | 72 | 29 | 60 | 970,047,104 | 734 | 4,642,256 |
| **+purge-**<br>**dups** | 3534 | 3491 | 43 | 28 | 78 | 929,446,752 | 485 | 5,136,536 |
| **hifiasm -**<br>**l3 Hap1** | 3549 | 3480 | 69 | 28 | 63 | 975,430,134 | 1,153 | 4,449,420 |
| **+purge-**<br>**dups** | 3522 | 3481 | 41 | 27 | 91 | 928,353,632 | 507 | 4,580,342 |
| **hifiasm -**<br>**l3 Hap2** | 3532 | 3461 | 71 | 32 | 76 | 964,775,996 | 729 | 4,275,860 |
| **+purge-**<br>**dups** | 3522 | 3481 | 41 | 33 | 85 | 938,372,303 | 515 | 4,317,088 |

Initial scaffolding of the primary assembly was performed by mapping HiC reads to the primary contigs using bwa-mem v0.7.17-r1198-dirty (H. Li, 2013) and mappings were filtered following the VGP Arima mapping pipeline (https://github.com/VGP/vgp-assembly/tree/master/pipeline/salsa). The final bed file was given to yahs v1.1a (Zhou et al., 2022) for scaffolding. Scaffolds were then manually curated into chromosomes using higlass v2.1.11 (Kerpedjiev et al., 2018) with Cooler v0.9.1 (Abdennur & Mirny, 2020) to visualise the HiC data (Supplementary Figures S1 - S2).

Finally, the two haplotype assemblies were polished by mapping the HiFi reads to the assemblies using pbmm2 v1.3.0 (https://github.com/PacificBiosciences/pbmm2.git) with arguments --preset CCS -N 1 and variants called using deepvariant v0.2.0 (Poplin et al., 2018) with --model_type=PACBIO. Errors were corrected in the assembly by filtering the vcf file given by deepvariant with bcftools *view* v1.12 (Danecek et al., 2021) with arguments *-i ’FILTER=\“PASS\” && GT=\“1/1\”’* and a consensus called with bcftools *consensus*. To determine whether any contaminant sequences were present in the genome, we screened for adapter, vector, and foreign sequences using NCBI Foreign Contamination Screen (FCS) v0.2.1 and FCS-GX v0.3.0 (Astashyn et al., 2024). Resulting hits indicative of contaminants were removed from the assembled sequences.

Additionally, we estimated the correctness of the assembly by counting the k-mers of the HiFi reads using FastK (https://github.com/thegenemyers/FASTK.git) with options “-v -t1” and “kmer=31” and then running Merqury.FK v.1 (https://github.com/marbl/merqury.git) on both phased haplomes. The analysis yields Merqury’s consensus Quality Value (QV), which is estimated by comparing the read and assembly k-mer counts and then transformed to a log-scaled probability of base-call errors. A higher QV indicates a more accurate assembly.

To detect any possible biases in the input data that might have led to misassemblies, we aligned the raw PACBIO reads, the HiFi and HiC reads back to each phased assembly. For the PACBIO raw data we used pbmm2 v.1.13 (https://github.com/PacificBiosciences/pbmm2.git) with options “--preset “SUBREAD” --sort-memory 20G” for each SMRT cell. For HiFi reads, we used minimap2 v. 2.26 (Li, 2018) with options “minimap2 -acyL --secondary=no --MD --eqx -x map-hifi -k 20” and for the HiC we used bwa-mem2 v.2.2.1 (Vasimuddin et al., 2019) with option “-M”. All resulting mapping files were sorted with SAMtools v1.17 (Li et al., 2009) *sort* and the mapping quality was estimated with qualimap v.2.3 (Okonechnikov et al., 2016) with options “-c -outformat pdf --java-mem-size=10G”.

A mitochondrial assembly was created using MitoHiFi v2.0 (Uliano-Silva et al., 2023) using the DeepConsensus PacBio Reads and *Phoxinus phoxinus* reference mitochondrial genome NC_020358.1 as input. Any partial mitochondrial contigs remaining in the assembly were removed based on mapping synteny to the fully assembled mitogenome.

### Annotation of Repetitive Elements

Repeat sequences were identified and annotated in both haplomes of the *P. phoxinus* reference genome by combining homology-based and *de novo* approaches. We identified low-complexity repeats and tandem repeats using DustMasker from BLAST+ v. 2.9 of the NCBI C++ toolkit (Morgulis et al., 2006) and Tandem Repeat Finder v4.10.0 (Benson, 1999), respectively. We *de novo* identified transposable elements (TEs) with Repeatmodeler v2.0.1 (Smit and Hubley, 2008) to create a species-specific library, and we combined the identified species-specific transposable elements (TEs) with the *Danio rerio* (zebrafish, Cypriniformes) repeats library from Dfam 3.1 (Hubley et al., 2016) to generate a custom combined repeats library.

Using this custom combined library, repetitive elements were identified and annotated in our genome with RepeatMasker v.4.1.0 (https://repeatmasker.org/). Divergence of our repeat annotation from the consensus repeats sequence was estimated with the *calcDivergenceFromAlign.pl* script implemented in RepeatMasker using the *.align* output file of the previous masking step. The repeats landscape was plotted with the *createRepeatLandscape.pl* script, also in RepeatMasker. The output files of Dustmasker and Tandem Repeats Finder were converted to gff3 format with the custom perl script repeat_to_gff.pl v. 1.0 and the output files of RepeatMasker were converted to *gff3* format with RepeatMasker’s rmOutTogff3.pl script and edited with a custom perl command to add a unique ID to each entry. We merged the positions of repeats with Bedtools v2.29.2 (Quinlan, 2014) before our genome was softmasked with these identified repeats using Bedtools in preparation for the prediction of protein-coding genes.

### Annotation of Protein Coding Genes

The annotation of protein-coding genes in the *P. phoxinus* genome was carried out with a combination of *ab initio*, protein similarity, and transcriptome-based protein prediction models. The sequencing reads (NCBI accession numbers SRR26699630 to SRR26699636) from seven RNA samples from brain, gill, gonad, liver, muscle, skin, and spleen were mapped to both haplomes with STAR v2.7.10b (Dobin et al., 2013), the resulting mapping files (.bam) were merged with SAMtools v1.10 (H. Li et al., 2009). BRAKER3 (Brůna et al., 2021; Gabriel et al., 2023; Hoff et al., 2016), the latest genome annotation pipeline in the BRAKER (Hoff et al., 2019) suite, was then used for annotation of protein-coding genes. With the RNA mapping files as input, BRAKER3 uses StringTie2 (Kovaka et al., 2019) to create a draft transcriptome, which then serves as the basis to predict protein-coding genes with GeneMarkS-T (Tang et al., 2015). The genes with the best similarity scores and quality of *ab initio* predictions are then selected with GeneMark-ETP (Brůna et al., 2020). Then, GeneMark-ETP uses the protein, RNA and ab initio-predicted genes to produce three groups of hints, i) hints with transcript and protein similarity support, ii) hints with transcript and *ab initio* support and finally, iii) hints with only protein similarity support using ProtHint. All these hints were used to create a set of high-confidence genes. AUGUSTUS (Stanke et al., 2006) was trained with these gene sets and performed an *ab initio* prediction of another genome-wide gene set before TSEBRA (Gabriel et al., 2021) combined the results of AUGUSTUS and GeneMark-ETP to produce a final set of high-confidence genes. The final set was used as input to BUSCO v5.4.7, which was run in protein mode with the actinopterygii_odb10 database.

Functional annotation of the resulting predicted genes was done with the *blastp* algorithm implemented in DIAMOND v2.1.8 (Buchfink et al., 2021). Our predicted protein-coding genes were blasted against protein databases of Swiss-Prot, edition 2023-05-13 (The UniProt Consortium, 2023), the Protein Data Bank (PDB) database, edition 2023-05-13 (RCSB.org, Berman et al., 2000) and TrEMBL, edition 2023-05-14 (The UniProt Consortium, 2023) The blast was done with an e-value cut-off of < 1*10^-5^. Functional annotation of these protein domains was done with eggNOG-mapper v2 (Cantalapiedra et al., 2021).

Protein sequences of *Danio rerio* (zebrafish, Cypriniformes), GCF_000002035.6 (Howe et al., 2013); *Sinocyclocheilus rhinocerous* (rhinoceros golden-line barbel, Cypriniformes), GCF_001515625.1 (J. Yang et al., 2016); *Sinocyclocheilus grahami* (golden-line barbel, Cypriniformes), GCF_001515645.1 (J. Yang et al., 2016); *Cyprinus carpio* (common carp, Cypriniformes), GCF_000951615.1 (Xu et al., 2014); *Carassius auratus* (goldfish, Cypriniformes), GCF_003368295.1 (Z. Chen et al., 2019); *Pimephales promelas* (fathead minnow, Cypriniformes), GCF_016745375.1 (Martinson et al., 2022); *Megalobrama amblycephala* (Wuchang bream, Cypriniformes), GCF_018812025.1(H. Liu et al., 2021); *Ctenopharyngodon idella* (grass carp, Cypriniformes), GCF_019924925.1 (Wu et al., 2022); *Myxocyprinus asiaticus* (Chinese high-fin banded shark, Cypriniformes), GCF_019703515.2 (Krabbenhoft et al., 2021); and the outgroup, *Ictalurus punctatu*s (channel catfish, Siluriformes), GCF_001660625.3 (Waldbieser et al., 2023) were retrieved from NCBI RefSeq database (O’Leary et al., 2016) on 2023-07-12. We included *Pimephales promelas* (Fathead minnow) as the species most closely related to *P. phoxinus,* for which a well-documented whole genome assembly and annotation are available. To identify similar sequences, first AGAT v1.1.0 (Dainat, 2021) was used to extract the longest gene isoforms, against which the protein-coding gene set of *P. phoxinus* was then mapped with DIAMOND v2.1.8 (Buchfink et al., 2021).

### Genome-wide Comparison between Haplomes

#### Sequence Variation

Genome-wide average heterozygosity was estimated for both haplomes to assess sequence variation using the 25 largest scaffolds representing the 25 chromosomes. To do so, raw HiFi reads were mapped to both haplomes with minimap2 v2.1 (H. Li, 2018) with preset *-ax asm20*, which optimises alignments between sequences with less than 20% divergence. The resulting alignment files were filtered for mapping quality lower than 20 with the view *-q 20* command and sorted with the *sort* command using SAMtools v1.10 (H. Li et al., 2009). Duplicates were removed with the *markdup* -r command as implemented in Sambamba v0.7.1 (Tarasov et al., 2015). Furthermore, we generated windows of 100 kb with the *makewindows* command implemented in Bedtools v2.31.0 (Quinlan, 2014) with *-w 100000.* Then, we estimated depth in 100 kb windows for both haplomes using *coverage -mean* as implemented in Bedtools.

Site allele frequency likelihood was estimated using the GATK model for genotype likelihoods in combination with the -*doSAF 1* parameter in ANGSD v.0.940 (Korneliussen et al., 2014), the site frequency spectrum was estimated with the *realSFS* parameter also in ANGSD. Finally, heterozygosity was estimated from sample allele frequencies in non-overlapping windows of 1 Megabase (Mb). The number of heterozygous alleles was counted within each Mb and then converted to the number of heterozygous alleles per kb and plotted per chromosome across the genome using ggplot2 v3.1.3 (Wilkinson, 2011), to visually inspect patterns of heterozygosity across the entire genome. Patterns of heterozygosity often reflect standing genetic variation which may be linked to the demographic history of a species (Guevara et al., 2021; Miller et al., 2014; Nadachowska-Brzyska et al., 2016).

#### Demographic History of P. phoxinus

The demographic history of *P. phoxinus* was reconstructed using Pairwise Sequentially Markovian Coalescent (PSMC) v0.6.5 as implemented by Li & Durbin, (2011). Variants were called from both haplomes with a combination of SAMtools v1.17 *mpileup* (H. Li et al., 2009) and Bcftools v1.17 *call* (Danecek et al., 2021) commands. The resulting variant file (.vcf) was converted to a diploid consensus file with *vcfutils.pl vcf2fq* accessory implemented in the PSMC package. The resulting fastq files were converted into the input fasta format for PSMC using the *fqpsmcfa* accessory. The PSMC model was run with the following parameters: *-N25 -t15 -r5 -p “4+25*2+4+6”* and 100 bootstraps, a generation time of 3 years (Museth et al., 2010), which is the maturation age of the *P. phoxinus* and an assumed mutation rate of 3.51 x10^-9^ (Tian et al., 2022). The results of the run were plotted using ggplot2.

#### Structural Variation

We aligned both haplomes with minimap2 v2.26 (H. Li, 2018) using the *-asm5* preset setting. Alignments were processed with Synteny and Rearrangements Identifier (SyRI) v.1.6.3 (Goel et al., 2019) to detect structural variation between both haplomes and the results were visualised using the R package plotsr v.1.1.1 (Goel & Schneeberger, 2022). Genes overlapping insertions, deletions, and inversions were extracted using Bedtools v2.29.2 (Quinlan, 2014). Subsequently, functional enrichment of these genes was achieved with g:profiler (Raudvere et al., 2019) using the zebrafish database option. Finally, the results were visualised by go-figure v1.0.2 (Reijnders & Waterhouse, 2021).

### Identification of Orthologs and Gene Family Evolution

As input to Orthofinder v2.5.4 (Emms & Kelly, 2019) with default settings, we used the same set of proteomes from species that we downloaded and described in the section on annotation of protein-coding genes, after filtering out amino acids shorter than 30 base-pairs (bp). Orthofinder created orthogroup gene trees and then inferred a rooted species tree from gene duplication events with the STRIDE approach (Emms & Kelly, 2017) using just single-copy orthologs. The species tree underwent conversion into an ultrametric tree utilising Orthofinder’s *make_ultrametric.py* script, employing the parameter -r 142. This value corresponds to approximately 142 million years, which aligns with the estimated median divergence time between Siluriformes and Cypriniformes, as suggested by the *TimeTree* database (Kumar et al., 2017). This estimate is calculated by taking into account the median of published estimated divergence times using molecular sequence data analysis (Kumar et al., 2017; Kumar & Hedges, 2011). Orthologous gene counts from the Orthofinder run were filtered to remove orthologous groups that are overrepresented in a particular species using the *clade_and_size_filter.py* script in CAFE v5.0.0 (Mendes et al., 2021). Gene family evolution analysis was carried out in CAFE v5.0.0 using the ultrametric tree and the filtered orthologous gene counts as input. Genes that were confirmed to be species-specific, expanding, and contracting in *P. phoxinus,* were mapped against the TrEMBL database with diamond *blastp*. The GO terms derived from the TrEMBL database were then used as input into REViGO (Supek et al., 2011) to remove redundant terms and summarise GO results.

## Results and Discussion

### K-mer Based Genome Size and Heterozygosity Estimation

Genome size estimations using k-mer lengths 19, 23, 25, and 30, showed slight differences in the estimated genome sizes and heterozygosities (Figure S3). We chose the 19-mer length due to a lower error rate in comparison to other k-mer lengths. We estimated a haploid genome size of *P. phoxinus* of 805.8 Mbp with a unique content of 62.1% and a heterozygosity of 1.43%. This estimate deviates from the determined C-value of 1.15 pg (corresponding to 1.13 Gbp) (genomesize database; https://genomesize.com/). The estimated heterozygosity of 1.43% is high compared to other published k-mer-based heterozygosity estimates in Cypriniformes. For example, *Onychostoma macrolepis* has an estimated heterozygosity of 0.29% (L. Sun et al., 2020), *Myxocyprinus asiaticus* (Chinese high-fin banded shark) of 0.20% (Krabbenhoft et al., 2021), *Gymnocypris przewalskii* (Przewalskii’s naked carp) of 0.96% (Tian et al., 2022), and *Poropuntius huangchuchieni* of 0.68% (L. Chen et al., 2021). To the best of our knowledge, the highest k-mer-based heterozygosity estimate for a cypriniform is 1.82% estimated in *Gymnocypris eckloni* (F. Wang et al., 2022), while the lowest estimate of 0.1% was found in *Triplophysa tibetana* (X. Yang et al., 2019). The clear bimodal coverage distribution seen in the k-mer plot (Figure 1) is indicative of the diploidy of the *P. phoxinus* genome, with the first peak at approximately 18-fold coverage corresponding to heterozygous sites and the second peak at approximately 36-fold coverage corresponding to homozygous genome positions.

**Figure 1:**
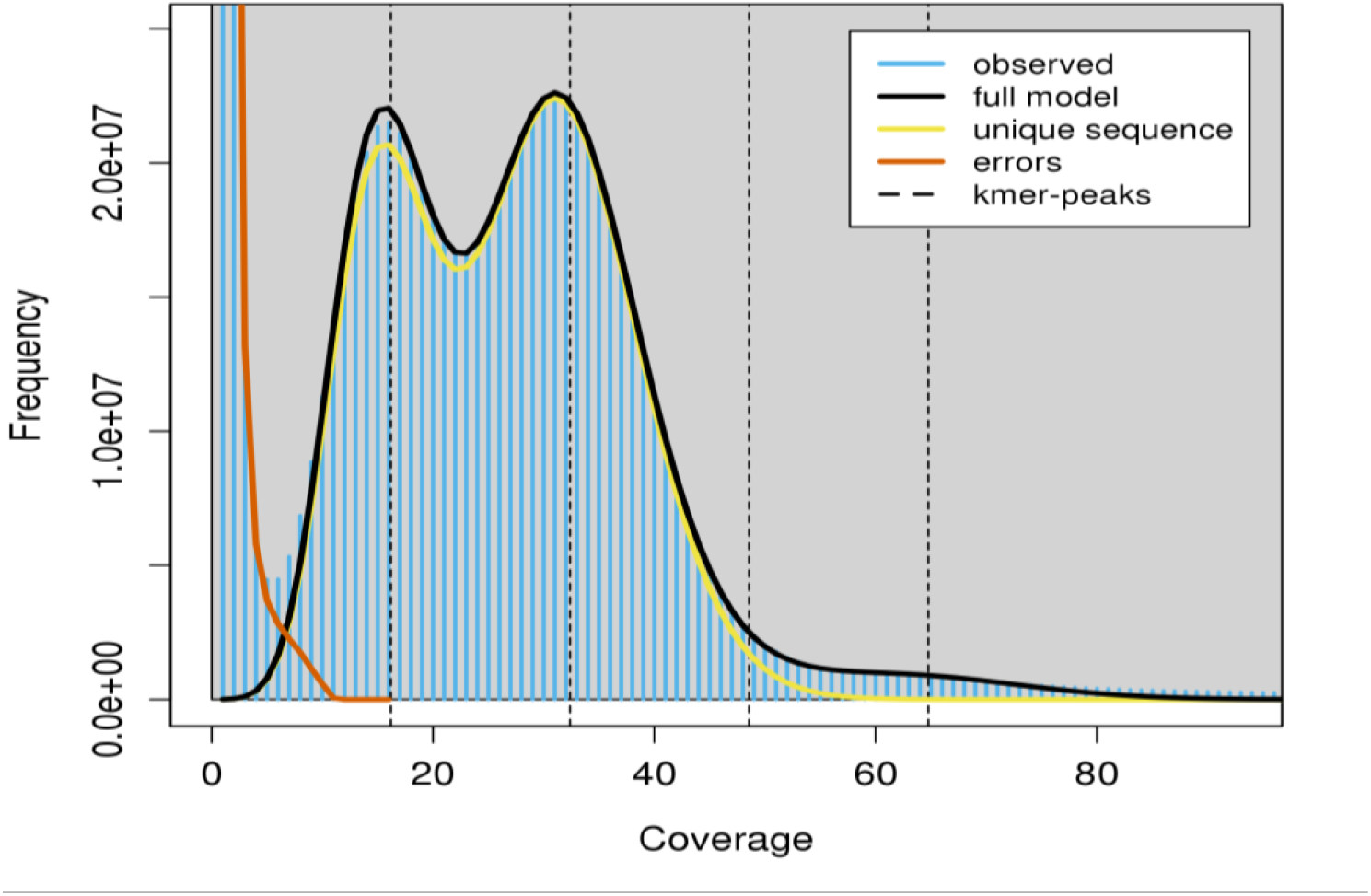
K-mer spectra profile of *Phoxinus phoxinus* generated from raw PacBio HiFi reads with GenomeScope2. The y-axis shows the k-mer counts and the x-axis shows sequencing depth. The clear bimodal pattern observed indicates a diploid genome with high heterozygosity of 1.43%. The haploid genome size was estimated to be around 805 Mb.

Utilising the proportion of the assembly sequence supported by HiFi reads, Merqury calculated a Quality Value (QV) of 58.9 (corresponding to 99.99871% accuracy) for haplome 1 (Hap1) and 58.8 (99.99868% accuracy) for haplome 2 (Hap2). Aligning the PacBio raw subreads, HiFi, and Illumina HiC reads to both haplomes revealed comparable coverage levels (600±378, 39.03±28, 83.6268±132-fold, respectively, for Hap1; 39±25, 84±82, 605±353-fold, respectively, for Hap2) and mapping quality scores (50, 52, 42, respectively, for Hap1; 50, 52, 43, respectively, for Hap2). These results further indicate that the assembly is well-phased with minimal assembly errors (Supplementary Table S1).

### Genome Assembly and Scaffolding

We generated a haplotype-resolved assembly of *P. phoxinus*, with a total assembly size of 940 Mbp and 929 Mbp for Hap1 and Hap2, respectively. The initial assembly of each haplome contained 394 primary contigs, which were further placed into 101 (N50: 36.4 Mb) and 81 (N50: 36.6 Mb) scaffolds, respectively, by mapping HiC reads to the primary contigs. Integrating Hi-C data, we ultimately scaffolded 99.6% of the Hap1 assembly and 99.5% of the Hap2 assembly each into 25 chromosomes (Supplementary Table S2). BUSCO analysis revealed the *P. phoxinus* haplomes had only 2.3% and 2.1% missing genes. The assembly statistics for both haplomes (Table 2) denote high contiguity and high completeness. The 11 Mb difference in size between both haplomes is a result of structural variations between haplomes.

**Table 2.** Summary of *P. phoxinus* haplome 1 (Hap1) and haplome 2 (Hap2) genome assembly statistics.

| Assembly Statistics | Hap1 | Hap2 |
| --- | --- | --- |
| Assembly Size | 940 Mb | 929 Mb |
| Contigs | 394 | 394 |
| Scaffolds | 101 | 81 |
| N50 Contigs | 3.85 Mb | 4.59 Mb |
| N90 Contigs | 869 kb | 1.05 Mb |
| N50 Scaffolds | 36.4 Mb | 36.6 Mb |
| N90 Scaffolds | 30.7 Mb | 30.3 Mb |
| % of Assembly in Scaffolds | 99.6% | 99.5% |
| Complete BUSCOs | 3528 (96.9%) | 3540 (97.2%) |
| Complete and Single-copy BUSCOs | 3488 (95.8%) | 3495 (96.0%) |
| Complete and duplicated BUSCOs | 40 (1.1%) | 45 (1.2%) |
| Fragmented BUSCOs | 30 (0.8%) | 25 (0.7%) |
| Missing BUSCOs | 82 (2.3%) | 75 (2.1%) |
| Total BUSCO Search | 3640 | 3640 |

### Annotation of Repetitive Elements

Teleost fish genomes are characterised by a wide range of repetitive elements (Gao et al., 2016). The genome of *P. phoxinus* follows this pattern, with approximately 506 Mbp (53.86%) of its assembly consisting of repetitive elements. This is higher than reported to date in other cyprinid species such as fathead minnow (43.27%) (Martinson et al., 2022), grass carp (43.26%) (Wu et al., 2022), common carp (31.3%) (Xu et al., 2014) or goldfish (39.6%) (Z. Chen et al., 2019). However, the repeat content is slightly lower than that of zebrafish, which contains around 54.47% repetitive elements (Howe et al., 2013).

From the identified TEs, DNA transposons were the most abundant repeat elements in the genome, making up 21.25% (approximately 199 Mb). DNA transposon content is similar to that of the fathead minnow genome, which contains an estimated 21.47% (Martinson et al., 2022). SINEs were relatively poorly represented at 1.03%, in line with other published fish genomes (Shao et al., 2019; Sotero-Caio et al., 2017). Retroelements made up 11%, LTRs 6.16% and LINEs 4.15%. Overall, we annotated 46.59% of the *P. phoxinus* genome to be interspersed repeats, the remaining repetitive elements were annotated as 0.47% small RNA, 1.50% satellites, 2.66% simple repeats, and 0.17% low complexity repeats (full details of all repetitive elements are described in Tables S3 and S4).

Repeat landscapes illustrate how TEs cluster in correlation to the Kimura substitution rate, i.e. how quickly TEs in a genome have diverged from the consensus TE sequence (Shao et al., 2019). The Kimura substitution level is correlated with the age of transposition activity: low substitution levels indicate recent transposition events; higher substitution levels suggest old transposition events. The repeat landscape of the annotated *P. phoxinus* genome (Figure 2) indicates a high level of active, i.e., recent, transposition bursts (Watanabe et al., 2014). In fish genomes, active transposons accumulate at a faster rate than they decline (Shao et al., 2019), leading to an abundance of active transposons with low Kimura substitution levels (Böhne et al., 2012; Gao et al., 2016; Sotero-Caio et al., 2017), an exact pattern observed in TEs in the *P. phoxinus* genome. For the repeat landscape of Hap2 we refer to Supplementary Figure S4.

**Figure 2:**
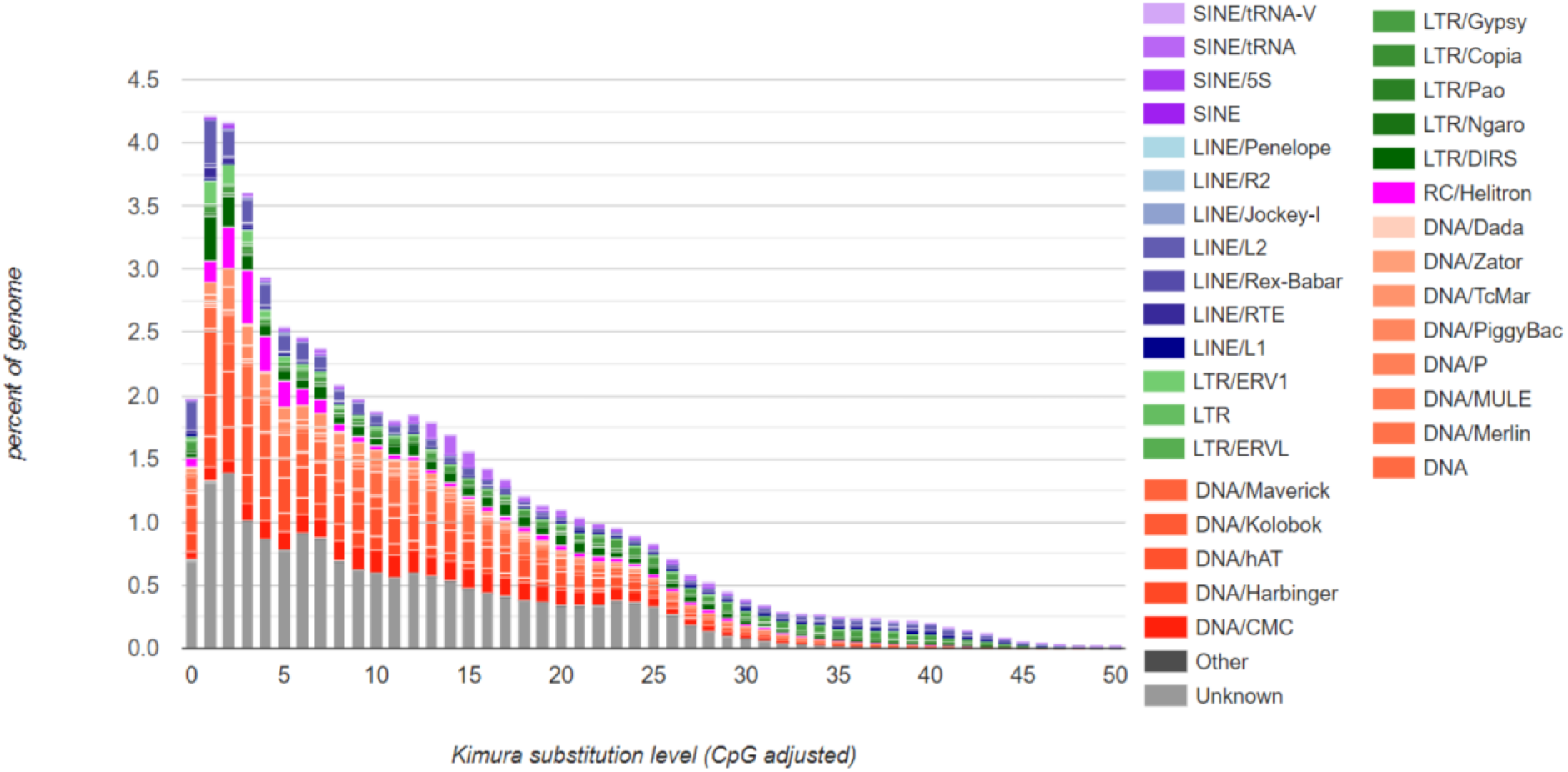
Repeat landscape of transposable elements (TEs) in the *Phoxinus phoxinus* genome across different Kimura substitution levels (in %). TEs on the left side of the histogram represent recently active TEs with low divergence from the consensus sequence of TEs, while TEs towards the right side of the histogram represent ancient TEs with higher degrees of divergence.

Active transposition and high repeat content in the *P. phoxinus* genome may result in rearrangements and variations in genome size (Kojima, 2019). Transposable elements can also function as recombination agents, generating structural variations such as deletions, insertions, translocations and inversions (Carvalho & Lupski, 2016; Cerbin & Jiang, 2018; Deininger et al., 2003), which can be sources of standing genetic variation present in an individual.

### Annotation of Protein Coding Genes

Prediction of protein-coding genes, based solely on transcripts derived from our RNA-seq data mapped to the haplomes (Supplementary Table S5), generated 40,364 gene models and a mRNA count of 42,408 in Hap1 and 38,754 gene models, and 40,716 mRNAs in Hap2. The *ab-initio* prediction resulted in 24,420 and 35,806 gene models in Hap1 and Hap2, respectively. After filtering and generating a final high-confidence gene set, the final annotation contained 30,980 mRNAs, of which 23,397 were marked as high-confidence genes in Hap 1; and 29,614 mRNAs, with 23,191 high-confidence genes in Hap 2 (full summary statistics of protein annotation for Hap1 and Hap2 in Supplementary Tables S6 and S7). We ran BUSCO with the actinopterygii_odb10 database and obtained a total of 3,484 (95.7%) complete BUSCOs, 3,435 (94.4%) complete and single-copy BUSCOS, 49 (1.3%) complete and duplicated BUSCOs, 20 (0.5%) fragmented BUSCOs and 136 (3.8%) missing BUSCOs. The predicted proteins were functionally annotated using a combination of similarity mapping to protein databases (Swiss-Prot, TrEMBL and PDB) and orthology assignment (eggNOG), resulting in 22,761 annotated genes (Table 3). These statistics hint at a well-assembled and annotated *P. phoxinus* genome. Similarity search of genes in *P. phoxinus* against zebrafish, fathead minnow, goldfish, common carp and grass carp resulted in 21,206 (90.6%) annotated genes across all 5 species (Supplementary Figure S5). Comparing the predicted gene content of Hap1 to its closest relative with available whole-genome resources, the fathead minnow contains more predicted genes (26,150) (Martinson et al., 2022). Still, for *P. phoxinus*, the complete BUSCO score of 95.7% (3,484) was higher compared to the fathead minnow’s 94.3% (Martinson et al., 2022).

**Table 3.** Functional annotation statistics for genes in *P. phoxinus* using TrEMBL, Swiss-Prot, PDB, and eggNOG indicate that 97.28% (22,761) of genes were successfully annotated, with 84.93% (19,871) present in all four databases.

| Annotation Database | Number of annotated genes identified in Hap1 | Percentage of annotated genes (%) |
| --- | --- | --- |
| Swiss-Prot | 19,978 | 85.39 |
| TrEMBL | 22,761 | 97.28 |
| egg-NOG | 21,296 | 91.02 |
| PDB | 19,978 | 85.39 |

### Genome-wide Comparison of Structural and Sequence Variation between Haplomes

#### Sequence variation follows universal pattern of eukaryotes

Previously, genomes with high heterozygosity presented assembly challenges due to multiple scaffold formation from the same sequence (Kajitani et al., 2014; Pryszcz & Gabaldón, 2016), which resulted in merging of sequence diversity into a single reference genome. However, the implementation of long-range information such as Hi-Fidelity long reads and Hi-C contact maps can aid in overcoming these difficulties by resolving the intra-individual sequence variation (Kronenberg et al., 2021). Our high-quality assembly, indicating increased levels of standing genetic variation, demonstrates the effectiveness of this approach. Previous studies have already indicated abundant levels of genetic variation in the *Phoxinus* genus (Bogutskaya et al., 2020; Reier et al., 2022).

Genome-wide estimates and patterns of mean heterozygosity were similar between both haplomes, with an estimate of 0.7866% in Hap1 and 0.7926% in Hap2. Assembly-based heterozygosity estimates were lower than k-mer-based estimates. A similar behaviour was reported in the Genomescope release paper (Vurture et al., 2017). In the present case, the difference between the two approaches used is likely the cause: the genome-wide approach generates heterozygosity estimates from direct observation of the confidently mapped loci as well as the thereof derived SNPs; the k-mer-based approach, however, additionally incorporates structural variants and is more sensitive to low coverage and error-prone regions and hence results in higher heterozygosity estimates. Heterozygosity estimates between the chromosomes varied, however, the pattern found on each chromosome was similar with regions of highest heterozygosity always found at the ends of chromosomes and the lowest heterozygosity in the central region (Figure 3).

**Figure 3:**
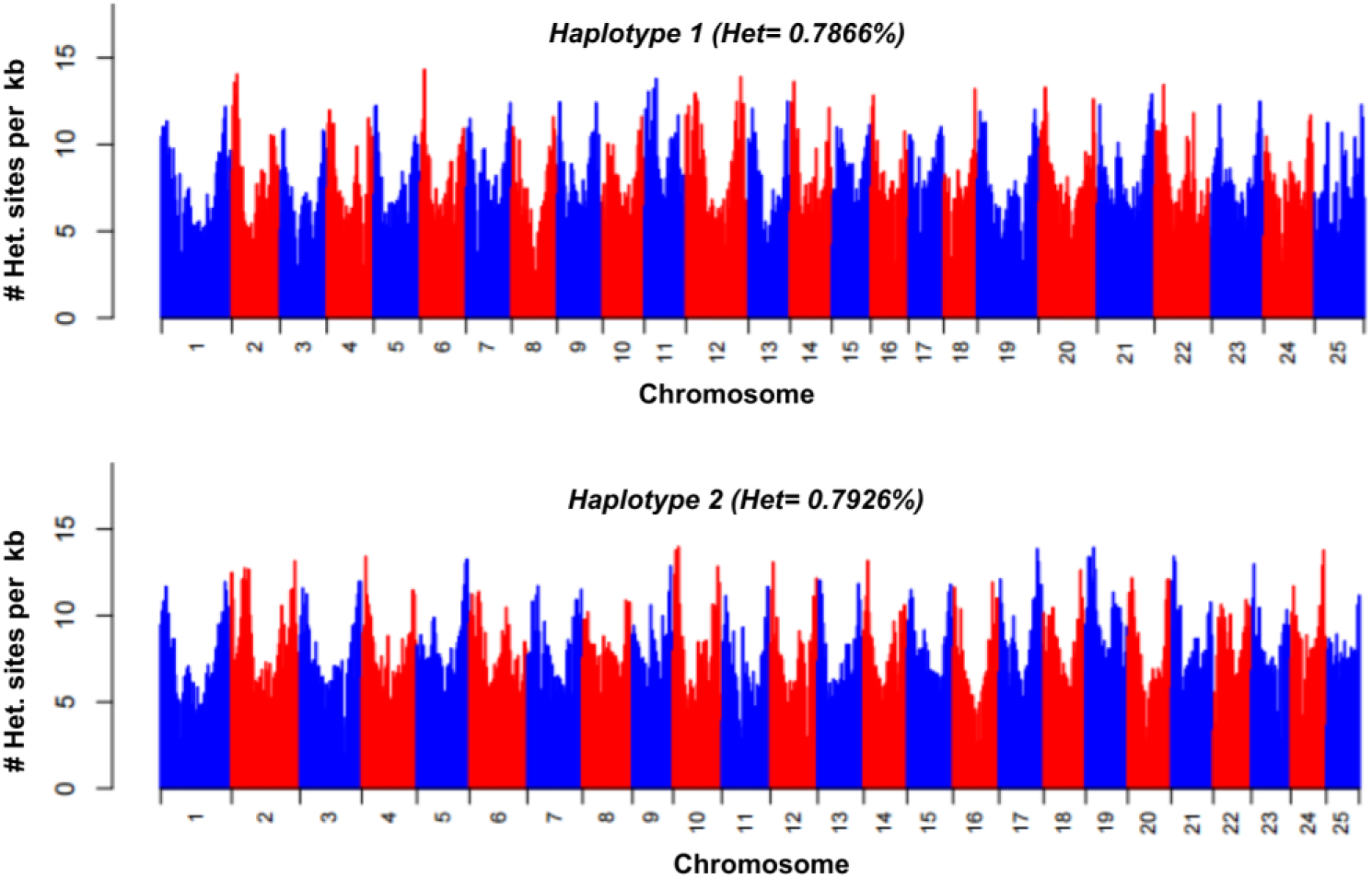
Distribution of heterozygosity along the 25 chromosomes in *P. phoxinus s*hown for Hap1 (upper plot) and Hap2 (lower plot). Mean heterozygosity estimates were calculated in 1Mb bins and converted to the number of heterozygous sites per kb across all 25 chromosomes with alternating colours to differentiate adjacent chromosomes.

Regions of high heterozygosity and high recombination rates may be linked to telomeric regions of the chromosome, which in turn can be associated with higher genetic diversity (Jaramillo-Correa et al., 2010). In fishes, this pattern was observed in the genomes of the Atlantic silverside (*Menidia menidia*) (Tigano et al., 2021), brown trout (*Salmo trutta)* (Leitwein et al., 2016) and sticklebacks (*Gasterosteus* spp.) (Sardell et al., 2018). This pattern is consistent with the pattern of heterozygosity observed in the *P. phoxinus* genome. Regions of reduced heterozygosity found at central locations may be associated with centromeres, which are known as “cold spots” for recombination in eukaryotes (Yan & Yu, 2022).

#### Demographic History of P. phoxinus

Climatic fluctuations have always affected the distribution and survival of species (Hewitt, 2000; J. Li et al., 2021). For aquatic species in the temperate zone, the effects of climate fluctuations due to glaciation cycles, which led to fluctuating sea water levels, were particularly pronounced (Munnecke et al., 2010). A PSMC analysis enabled us to track fluctuations in effective population size (*Ne*) of *P. phoxinus* from more than 10 million years ago (mya) until approximately 10 thousand years ago (kya).

We found a gradual and continuous rise in *Ne* with time, from roughly 20,000 individuals about 10 mya to around 25,000 individuals approximately 800 kya (Figure 4). This incremental increase coincides with the Neogene period, which is divided into the Miocene and the Pliocene epochs — both known for their global warmth and minimum ice volumes (Böhme et al., 2011; Haywood et al., 2000; Hui et al., 2018; Steinthorsdottir et al., 2021). The climate conditions prevailing during this timeframe suggest the existence of an environment favourable for the propagation and development of freshwater fishes. Following a small increase in *Ne*, the population declined to fewer than 20,000 individuals between 800 kya and 100 kya, during the mid-Pleistocene period, which was marked by severe quaternary glaciation and low global sea levels. A major glaciation in this period is the Saalian glaciation of Northern and North Central Europe from approximately 400 – 150 kya (Lang et al., 2018; Lauer & Weiss, 2018). During this epoch, temperatures were on average 11°C lower than current values (Hughes et al., 2007) and lowland glaciation periods were frequent (Rose, 2009). As a result, species either became extinct, migrated to warmer territories in southern Europe seeking refuge, or underwent adaptation (Feliner, 2011).

**Figure 4.**
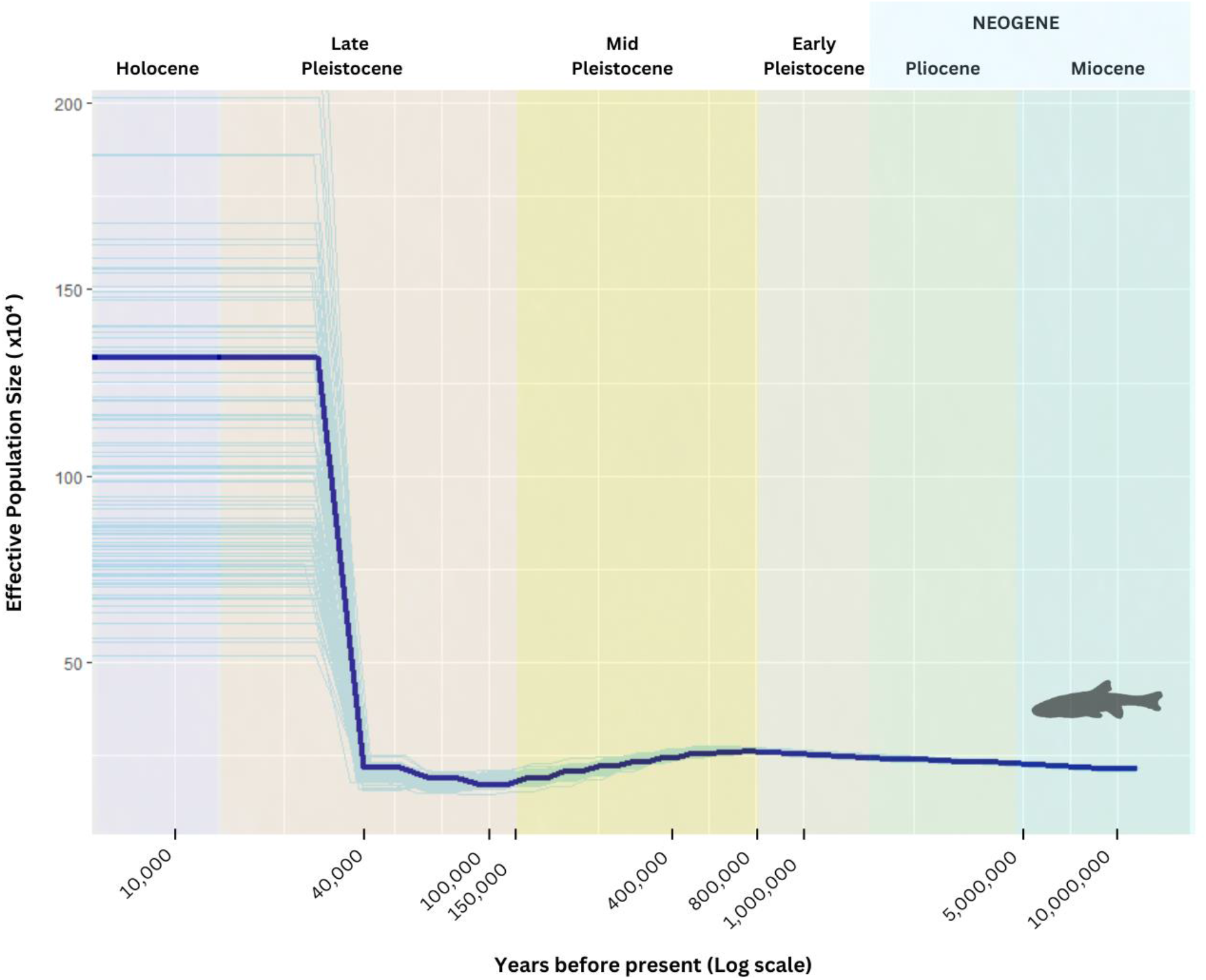
Inferred demographic history of *P. phoxinus* from PSMC analysis based on Hap1. The time ranges cover the Miocene, Pliocene, Early to Late Pleistocene, and Holocene, each shown in different colours. The x-axis shows years before present on a logarithmic scale and the y-axis shows the estimated effective population size. Bootstrap results are shown as transparent lines. Hapl2 showed a similar curve.

The reduction in *Ne* during mid-Pleistocene glaciations (Saalian glaciation) is generally observed in fishes (Brahimi et al., 2016; Gharrett et al., 2023), mammals (Fedorov et al., 2008), and birds (Kozma et al., 2016), especially in European species (Yuan et al., 2018). In *P. phoxinus*, the end of the mid-Pleistocene from around 40 kya was followed by a gradual and eventually sharp increase in *Ne*, peaking at nearly a million estimated individuals, indicating that *P. phoxinus* was able to repopulate previously covered freshwater habitats across Europe.

#### Large non-syntenic regions between P. phoxinus haplomes

A comparison of both haplomes of *P. phoxinus* revealed around 815 to 820 Mb of syntenic regions (Table 4, Figure 5). We also identified 1,572 translocations of 5.34 Mb and 5.36 Mb in both haplomes; 214 duplications totalling 571 kb in Hap1 and 667 duplications of around 1.8 Mb total length in Hap2; 215 inversions with a total length of 10.8 Mb in Hap1 and 11.1 Mb in Hap2 (Table 4, Figure 5).

**Figure 5:**
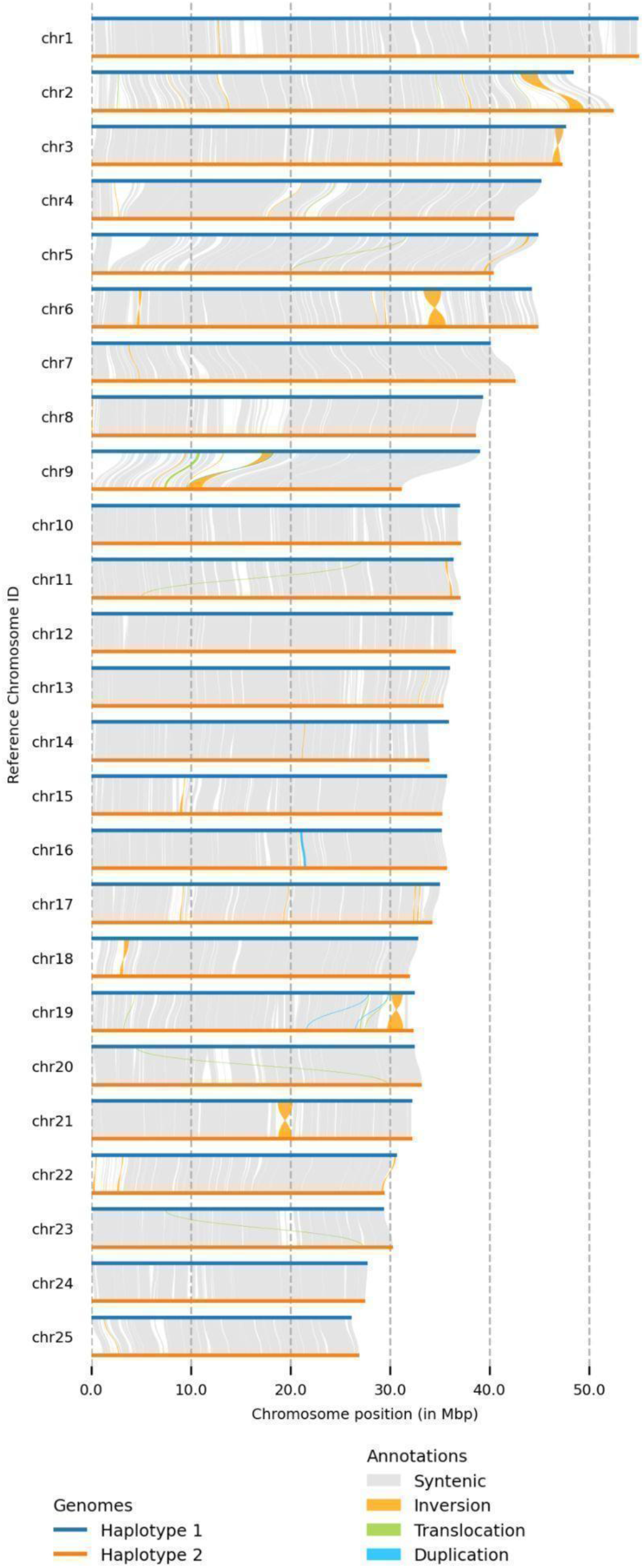
Synteny plot of all 25 chromosomes of the *P. phoxinus* Hap1 (top, blue line) and Hap2 (bottom, orange line) of assemblies. Aligned regions between both haplomes are shown in blocks of grey, unaligned regions are shown in gaps of white. Inversions are represented in orange, translocations in green, and duplications in light blue.

**Table 4.** Identity, number, and total lengths of identified structural variations between Hap1 and Hap2 in the *P. phoxinus* reference genome.

| Structural Variation | Count | Length Hap1<br>[bases] | Length Hap2<br>[bases] |
| --- | --- | --- | --- |
| Syntenic regions | 1457 | 813,594,094 | 820,681,234 |
| Inversions | 215 | 10,843,815 | 11,161,414 |
| Translocations | 1572 | 5,342,064 | 5,361,801 |
| Duplications (Hap1) | 214 | 57,1045 | - |
| Duplications (Hap2) | 667 | - | 1,776,709 |
| Insertions | 447,578 | - | 8,108,033 |
| Deletions | 447,699 | 8,113,615 | - |

Inversions represented the dominant structural variation within the *P. phoxinus* genome in terms of total size in either haplome (Table 4). The largest eight inversions made up 72% (8 Mb) of the estimated 11 Mb of inversions present in the *P. phoxinus* genome. Using Hap1 as a reference for inversion size, chromosome 6 contained the largest inversion with a size of 1.8 Mb; closely followed by a 1.7 Mb inversion on chromosome 2; a 1.5 Mb inversion on chromosome 21; a 1.1 Mb inversion on chromosomes 9 and 19, each; and finally, an 800 kb inversion on chromosome 3 (Figure 7).

**Figure 6:**
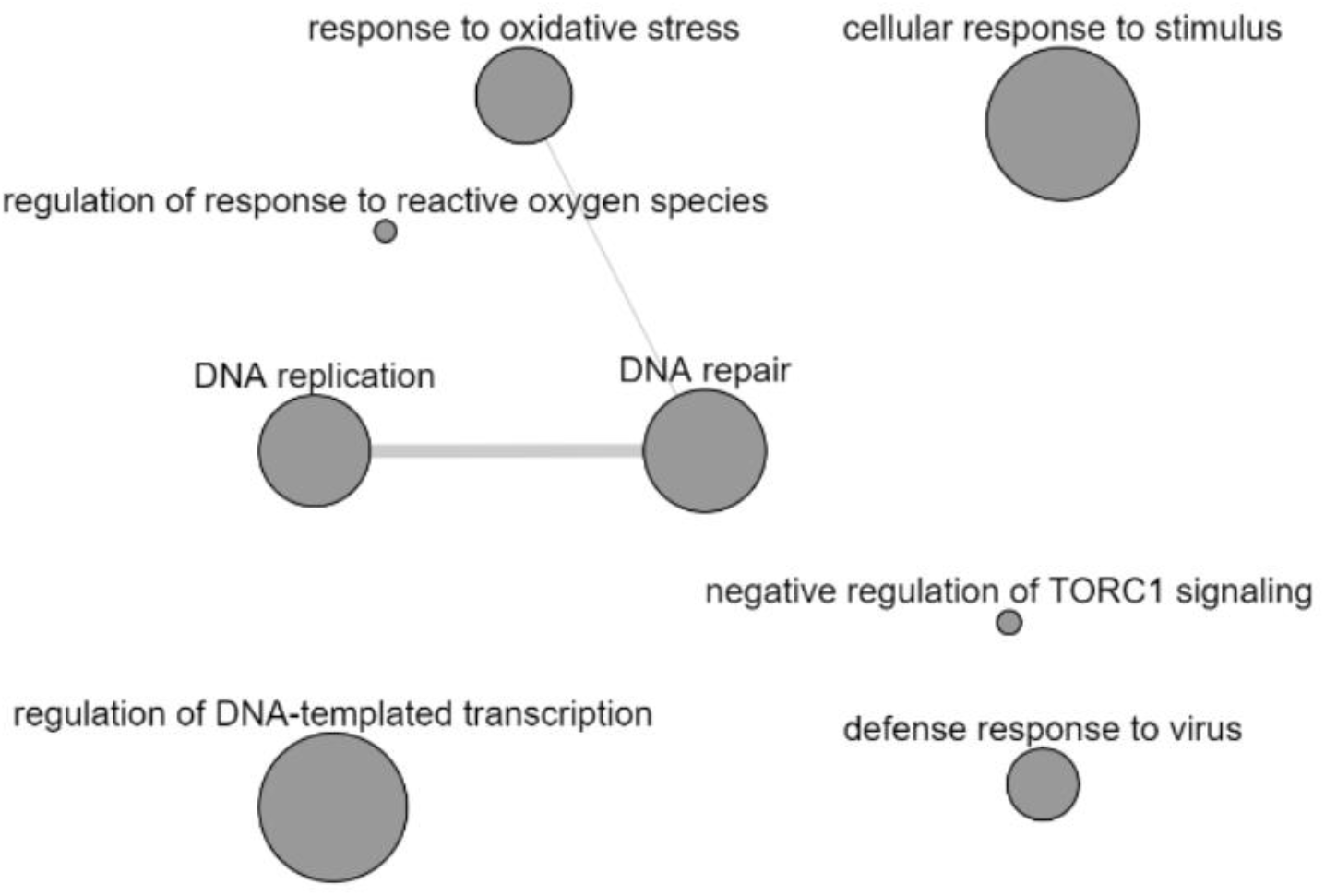
Gene ontology (GO) network of 54 functionally annotated *P. phoxinus-*specific genes in eight major GO terms: “DNA replication”, “DNA repair”, “regulation of DNA-templated transcription”, “response to oxidative stress”, “defense response to virus”, “cellular response to stimulus”, “regulation of response to reactive oxygen species” and “negative regulation of TORC1 signalling”. Bubble size indicates the frequency of the GO term in the gene ontology annotation database i.e larger bubbles correspond to more general terms. Edges on the graph connect highly similar or linked GO terms to each other.

**Figure 7:**
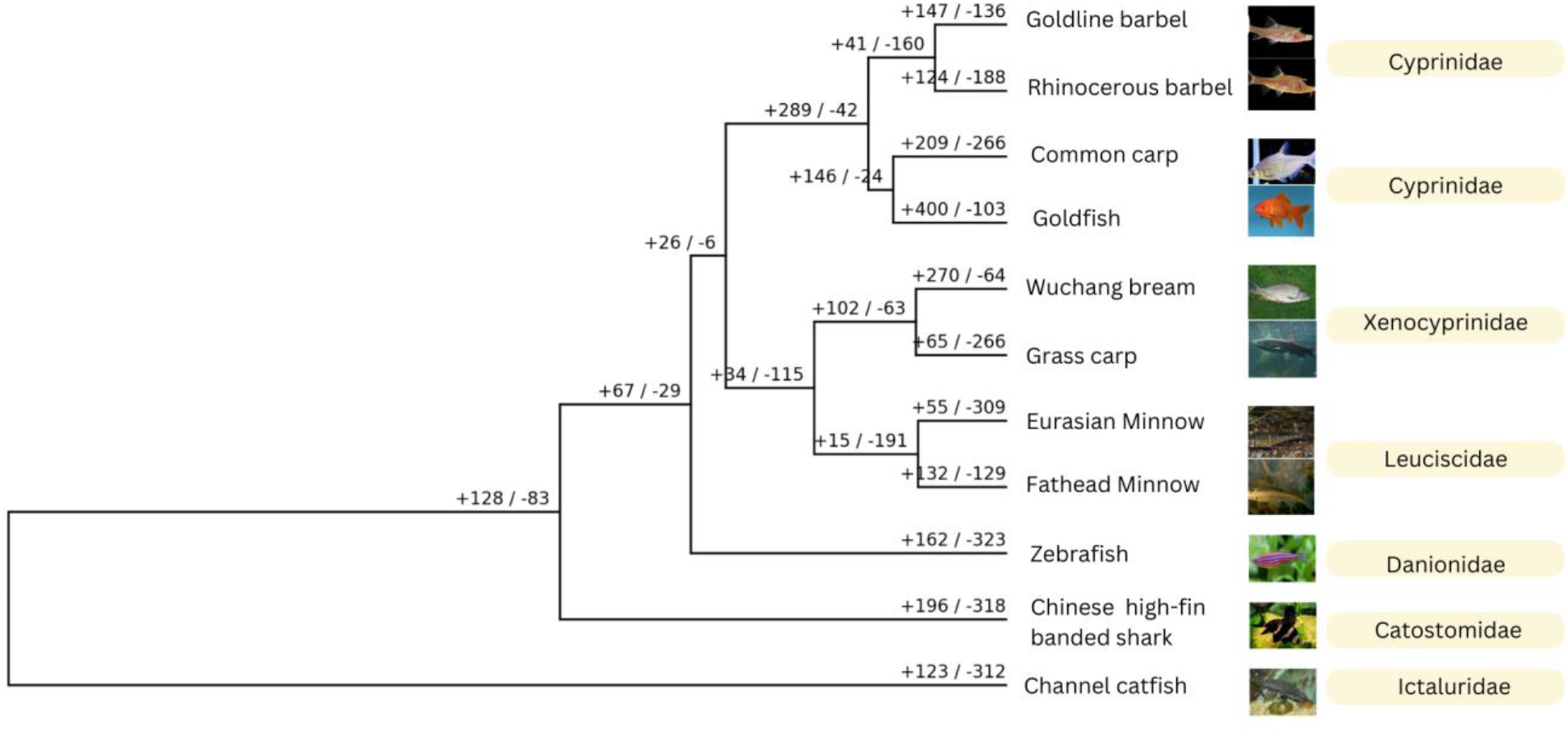
Ultrametric phylogenetic tree of selected teleost species and *P. phoxinus*. Numbers on the branches represent counts for expanded (+) and contracted (-) gene families. Positive numbers indicate significant gene families that have been expanded since the split from the last common ancestor and negative numbers indicate significant gene families that have been contracted since the split from the last common ancestor.

We investigated what type of genes were enriched in regions of copy number structural variants, specifically deletions and insertions, which can also be characterised as presence-absence variations (PAVs). GO terms overrepresented in PAVs between both haplomes included terms such as “dephosphorylation”, “peptidyl-serine phosphorylation”, and “protein processing”, which have been linked with the activation of sperm motility in numerous fish species (Dumorné et al., 2018; Zilli et al., 2008, 2017). GO terms related to development were also enriched within PAVs, such as “regulation of developmental growth” and “developmental pigmentation” (Supplementary Tables S8-S9 and Figure S6). GO terms overrepresented within inversions, were primarily associated with immunity and transposable elements in hAT, DDE, and L1 families: namely “phagocytosis”, “phagosome maturation involved in apoptotic cell clearance” and “phagolysosome assembly involved in apoptotic cell clearance”. Phagocytosis is a conserved immune defence mechanism across multicellular organisms that prevents infection and invasion of the body by pathogens (Esteban et al., 2015; Neumann et al., 2001) (Supplementary Tables S10-S11 and Figure S7).

#### Haplotype-resolved genome assembly revealed high level of within-genome variation in P. phoxinus

Analysis of mean genome-wide heterozygosity and structural variation, as outlined in the preceding sections, revealed a high level of variation present between the *P. phoxinus* haplomes. This is a possible outcome of high fecundity and large population size, expected in most freshwater fish species (Karjalainen et al., 2016; Röpke et al., 2021). The detection of genetic diversity in highly heterozygous species can be compromised by either un-phased genome-assembly approaches or haplotype-merged assemblies, which would ultimately lead to a significant underestimation of true genetic diversity (Nakandala et al., 2023; Takeuchi et al., 2022). Collapsed reference genomes restrict assembly accuracy and exploration of heterozygous regions of the genome, obscuring potentially interesting intra-individual variation (Low et al., 2020).

### Gene Family Evolution — Histone Gene Families Retracted, NLRC Genes Expanded in *P. phoxinus*

Inferring gene orthology from sequence data is a fundamental approach for understanding evolutionary relationships between genes and species through their evolutionary history (Tekaia, 2016). A total of 396,455 genes across all 11 species described in the section on annotation of protein-coding genes were included and 381,263 (96.12%) of these genes were successfully assigned to 27,672 orthogroups, i,e. gene sets in those species or in a subset of these species, that derived from their respective last common ancestor (Supplementary Table S12). The total number of orthogroups comprising protein-coding genes from all species was 14,914 (Table 5). Single-copy orthogroups, also known as 1-to-1 orthogroups, contain genes that have evolved from a common ancestral gene and are present in a single copy across all proteomes used in an ortholog analysis (Creevey et al., 2011; Simão et al., 2015; Tekaia, 2016). Our orthology inference analysis revealed the presence of 301 single-copy orthogroups which provided a solid basis for subsequent phylogenetic studies.

**Table 5.** Summary statistics of orthology inference analysis of *P. phoxinus* and 10 other teleost species.

| Overall Summary Statistic |  |
| --- | --- |
| Number of species | 11 |
| Number of genes | 396,455 |
| Number of genes in orthogroups | 381,263 |
| Number of unassigned genes | 15,192 |
| Percentage of genes in orthogroups | 96.2 |
| Percentage of unassigned genes | 3.8 |
| Number of orthogroups | 27,672 |
| Number of species-specific orthogroups | 2319 |
| Number of genes in species-specific orthogroups | 7975 |
| Percentage of genes in species-specific orthogroups | 2.0 |
| Mean orthogroup size | 13.8 |
| Median orthogroup size | 15.0 |
| Number of single-copy orthogroups | 301 |
| Number of orthogroups with all species present | 14,914 |

There were 273 genes distributed across 72 *P. phoxinus*-specific orthogroups; a sequence similarity search of these genes to the TrEMBL database returned results for only 130 of these genes, many of which were uncharacterised proteins (Supplementary Table S13). Only 54 of the successfully mapped genes annotated in the TrEMBL database returned GO annotations, which clustered into eight major GO terms (Figure 6). The presence of GO terms “response to oxidative stress” and “regulation of response to reactive oxygen”, suggests that *P. phoxinus* has adaptive potential to changes in water quality. Observational studies have found conflicting evidence, with Eurasian minnows ranging from sensitive (Bagge & Hakkari, 1992) to insensitive to intermediate pollution (King et al., 2011). Those findings may be attributed to the fact that different species of *Phoxinus* were studied. Cyprinoidea in general have been thought to be sensitive to oxidative stress, demonstrated in Amur minnow (*Phoxinus lagowskii*) (Q. Wang et al., 2019; Yao et al., 2022) and Chinese rare minnow (*Gobiocypris rarus*) (Z.-H. Li et al., 2014), while the fathead minnow (*Pimephales promelas*) has been used as a bioindicator of water quality (Elizalde-Velázquez et al., 2020).

Gene family changes are expressed as duplication and loss of genes among related species and are based on mutation, fixation, and retention rates of the genes in question (Demuth & Hahn, 2009). Gene family expansion might be an indication of positive selection (Demuth & Hahn, 2009). An example is the expansion of Myxovirus resistance (Mx) genes in the rainbow trout, providing special antiviral resistance for their anadromous lifestyle (T. Wang et al., 2019). While the immune system is relatively conserved across different jawed vertebrates (Flajnik & Kasahara, 2010; Litman et al., 2010), a wide array of species-specific modifications in the underlying immune pathway were observed in other osteichthyes, such as cods, coelacanths, but also chondrichthyes like sharks (Buonocore & Gerdol, 2016; Malmstrøm et al., 2016; Stein et al., 2007).

We analysed gene family expansions and contractions in *P. phoxinus* relative to 10 other teleost species (Supplementary Figure S8). Since the split of *P. phoxinus* and the fathead minnow from their last common ancestor, *P. phoxinus* counted more significant gene contractions (309 gene families) than significant gene expansions (55 gene families) relative to the fathead minnow, which had 132 gene family expansions and 129 gene family contractions (Figure 7). The significantly expanded gene families contained genes such as HIST1, H2B, CENP-T, GCRV, HIST2H2A, HIST2H3A, H3F3A, TRBV25-1, which mostly belong to histone PFAM domains (Supplementary Figure S9). Immune genes belonging to the NACHT PFAM domain were also more abundant in the set of contracted gene families (Supplementary Figure S10, genes like NLRP12, NLRP3, CD22, CLEC17A, GIMAP7). The expanding immune gene families mostly belonged to histone gene families, whereas the contracted gene families were mostly represented by NLRC gene families (Supplementary Tables S14 and S15).

Evolution of the immune system has greatly contributed to speciation in teleosts and a lot of diversity in their immune system pathways has been identified, like the expanded MHC1 and TLE genes in the Atlantic cod (*Gadus morhua*) (Star et al., 2011) and the absence of MHC class II-mediated immunity in the broad-nosed pipefish (*Syngnathus typhle)* (Haase et al., 2013). In the *P. phoxinus* genome, we observed an expansion in immunity-boosting histones H1, H2B, H3, and H4, particularly H2A and H2B. Histones are highly conserved proteins involved in the innate host defence against microbes (X. Li et al., 2022); they include core histones like H2A, H2B, H3, and H4. The final histone type, called H1, is generally referred to as a linker histone (X. Li et al., 2022; Westman et al., 2015). Histones are known to play a key role in antibacterial immunity across multiple species (X. Li et al., 2022; Pasupuleti et al., 2012). Histone 2A and 2B have antibacterial activity against different types of bacteria in fish. Histones play a distinctive role in mucosal immunity in fish (Ángeles Esteban & Cerezuela, 2015), acting as a strong barrier to microbial invasion. This has been confirmed in the Indian major carp (*Labeo catla*) (Nigam et al., 2017), lumpfish (*Cyclopterus lumpu*s) (Patel & Brinchmann, 2017), gilthead seabream (*Sparus aurata*) (Jurado et al., 2015), and rainbow trout (*Oncorhynchus mykiss*) (Fernandes et al., 2002; Oscoz et al., 2005). The expansion of histone genes in *P. phoxinus* suggests a strong focus on mucosal protection against bacteria and other pathogens. Assuming, that similar evolutionary trends can be found in other *Phoxinus* species, this is a particularly interesting find, given that there are populations of *Phoxinus* spp. in the Baltic Sea, some of which are anadromous and some of which are entirely resident in the brackish water (Svirgsden et al., 2018). Such migration habits demand an efficient immune system to cope with the different pathogens and parasite communities present in brackish and freshwater. Histones also play important roles in gene expression, DNA replication, and DNA damage repair (Best et al., 2018; Seal et al., 2022), in addition to their involvement in immune activity. It is important to note that histones are not restricted to immune activity alone. In fish, they have also been linked to spermatogenesis and can be an indicator of sperm quality (Herráez et al., 2017).

In contrast, NLRC genes were found to be reduced in the *P. phoxinus* genome, particularly NLRP3 and NLRP12, which are important inflammatory genes (Morimoto et al., 2021). NLRC genes are majorly involved in the immune response to exogenous pathogens like bacterial and parasitic infections (Chang, 2023; Morimoto et al., 2021). NLRC genes are involved in immune response in Nile tilapia (*Oreochromis niloticus*) to *Streptococcus agalactiae* and *Aeromonas hydrophila* infections (Q. Li et al., 2022). NLRC genes were also overexpressed in response to *V. anguillarum* infection in the Mi-iuy croaker (*Miichthys miiuy*) (G. Liu et al., 2023). In summary, the increased presence of genes involved in preventive mucosal defence and decreased number of genes involved in immune responsiveness might hint to a change in immune response strategy towards infection prevention. All the identified genetic diversity is potentially enabling more expeditious responses to environmental changes and even greater potential for adaptation across diverse environments (Lai et al., 2019; Matuszewski et al., 2015).

## Conclusion

In this paper, we present a highly contiguous and complete genome assembly for the Eurasian minnow, *Phoxinus phoxinus,* achieved by combining Pacbio Hifi reads and Hi-C data, along with a comprehensive repeats and gene annotation. Demographic history analysis and high observed heterozygosity and structural variation seem to suggest substantial population and individual diversity, respectively. While the identified sequence diversity could pose a challenge for a collapsed assembly, our phased genome enables a more accurate depiction of intra-individual variation. Consequently, we anticipate that this genome will prove an invaluable resource for forthcoming research investigating the intricate taxonomic and demographic complexities of the *Phoxinus* genus. For future studies, it will be of great importance to determine the genetic identity of the *Phoxinus* spp. studied before associating natural history traits with the species name. Similarly, a comprehensive understanding of the behaviour of Eurasian minnows requires tracing the possible genetic lineages in historical literature, in order to obtain a clearer picture.

## Acknowledgments

We thank the Long Read Team of the DRESDEN Concept Genome Center, part of the MPI-CBG and the technology platform of the CMCB at the TU Dresden, supported by DFG (INST 269/768-1).

## Author Contributions

MSt devised the project. TOO, MSt, AB, and IC led the writing and devising of the manuscript. EWM, SW, and TB performed genome sequencing and assembly. IC established the genome repeat and protein-coding gene annotation pipeline with support from SM. TOO performed genome-wide comparisons, gene family evolution, and demographic history analyses. SK performed DNA extractions and provided laboratory expertise. SM and AB provided TOO and IC with bioinformatic support. AB contributed with genome assembly expertise and input on study design. All authors contributed to revising the manuscript.

## Data, scripts, code, and supplementary information availability

Scripts used for genome assembly by EMW, SW and TB, and scripts used for assembly, genome-wide comparison, gene family evolution and demographic history by TOO are available here: https://doi.org/10.5281/zenodo.12191118.

The repeat and protein annotation were implemented using a snakemake pipeline. The code is available here: https://doi.org/10.5281/zenodo.11925110. RNA data for annotation is available under NCBI accession numbers SRR26699630 to SRR26699636. *P. phoxinus* haplomes and annotations are available under NCBI Bioproject number PRJNA1040855, which is linked to the Euro-Fish project under umbrella Bioproject number PRJNA768423.

## Appendix

Supplementary figures S1 - S8 and tables S2 to S5: https://zenodo.org/doi/10.5281/zenodo.10218893

Supplementary tables S1, S6 to S15: https://zenodo.org/doi/10.5281/zenodo.10220265

## Supplementary figures

• **Figure S1:** Heatmap of haplome 1 Hi-C assembly with darker blocks indicating higher intensity of sequence interaction
• **Figure S2:** Heatmap of haplome 2 Hi-C assembly with darker blocks indicating higher intensity of sequence interaction
• **Figure S3:** GenomeScope plots of 19, 23, 25 and 30 mer analysis
• **Figure S4:** Repeats landscape of annotated repeats in haplome 2
• **Figure S5:** Shared homologs between *P. phoxinus* and zebrafish, fathead minnow, goldfish, grass carp and common carp
• **Figure S6:** Gene ontology of genes in Insertions/Deletions
• **Figure S7:** Gene ontology of genes in Inversions
• **Figure S8:** Expansion/contraction profile including all gene families (significant and non-significant families)
• **Figure S9:** Pfam domain distribution of genes expanding in *P. phoxinus* genome
• **Figure S10:** Pfam domain distribution of genes contracting in *P. phoxinus* genome

## Supplementary tables

• **Table S1:** Mapping statistics of both haplomes
• **Table S2:** Genome assembly statistics of 25 chromosomes of both haplomes
• **Table S3:** Statistics of repeat sequences annotated in haplome 1
• **Table S4:** Statistics of repeat sequences annotated in haplome 2
• **Table S5:** Mapping statistics of RNA sequence data used for annotation of protein-coding genes in both haplomes
• **Table S6:** Summary statistics of annotated protein-coding genes in haplome 1
• **Table S7:** Summary statistics of annotated protein-coding genes in haplome 2
• **Table S8:** List of genes present in insertion/deletions
• **Table S9:** Gene ontology of genes present in insertions/deletions
• **Table S10:** List of genes present in inversions
• **Table S11:** Gene ontology of genes present in inversions
• **Table S12:** Summary statistics of orthology between *P. phoxinus* proteomes and selected teleosts
• **Table S13:** List of genes identified by orthology analysis as *P. phoxinus-*specific
• **Table S14:** List of genes significantly expanding in *P. phoxinus* genome
• **Table S15:** List of genes significantly contracting in *P. phoxinus* genome

