## Supplemental figures S1 to S10, and tables S2 to S5 for "A chromosome-level, haplotype-resolved genome assembly and annotation for the Eurasian minnow (Leuciscidae: *Phoxinus phoxinus*) provide evidence of haplotype diversity"

### shared first authorship

*Corresponding author

#### Table of Contents

**Table of Contents** 2

**Supplementary Tables** 3

Table S2: Summary statistics of the 25 largest chromosomes of both haplomes 3

Table S3: Statistics of repeat sequences annotated in Haplotype 1 4

Table S4: Statistics of repeat sequences annotated in Haplotype 2 4

Table S5: Summary mapping statistics of RNA sequence data used for protein annotation in the Phoxinus phoxinus genome 4

**Supplementary Figures** 5

Figure S1: Heatmap of Haplotype 1 Hi-C assembly with darker blocks indicating higher intensity of sequence interaction 5

Figure S2: Heatmap of Haplotype 2 Hi-C assembly with darker blocks indicating higher intensity of sequence interaction 6

Figure S3: GenomeScope plots of 19, 23, 25 and 30 mer analysis 7

Figure S4: Repeat Landscape of annotated repeats in Haplotype 2 8

Figure S5: Shared homologs between Phoxinus phoxinus and zebrafish, fathead minnow, goldfish, grass carp and common carp. 9

Figure S6: Gene ontology of genes in Insertion/Deletions 11

Figure S8: Expansion/contraction profile including all gene families (significant and non-significant families) 13

Figure S9: Pfam domain distribution of expanded genes 14

Figure S10: Pfam domain distribution of contracted genes 15

#### Supplementary Tables

---

**Table S2: Summary statistics of the 25 largest chromosomes of both haplomes**

| Chromosome | Length Hap1 | Length Hap2 | N_counts Hap1 | N_counts Hap2 | GC_counts Hap1 | GC_counts Hap2 |
|---|---|---|---|---|---|---|
| chr1 | 54,986,461 | 54,993,545 | 6080 | 5480 | 21,640,651 | 21,621,225 |
| chr2 | 48,474,465 | 52,476,322 | 4080 | 4000 | 19,024,086 | 20,630,447 |
| chr3 | 47,672,159 | 47,342,291 | 6440 | 2600 | 18,449,742 | 18,281,120 |
| chr4 | 45,191,066 | 42,481,172 | 4240 | 5800 | 17,541,055 | 16,516,799 |
| chr5 | 44,890,755 | 40,393,998 | 3320 | 3720 | 17,622,671 | 15,727,182 |
| chr6 | 44,260,849 | 44,933,711 | 6160 | 5000 | 17,588,082 | 17,886,922 |
| chr7 | 40,134,292 | 42,607,789 | 4400 | 4680 | 15,662,236 | 16,632,902 |
| chr8 | 39,332,310 | 38,661,445 | 4000 | 4200 | 15,514,313 | 15,225,733 |
| chr9 | 39,047,206 | 31,210,743 | 2600 | 1880 | 15,680,757 | 12,369,284 |
| chr10 | 37,018,565 | 37,176,178 | 4600 | 3400 | 14,417,725 | 14,477,234 |
| chr11 | 36,381,605 | 37,099,300 | 5000 | 6000 | 14,084,356 | 14,391,568 |
| chr12 | 36,320,318 | 36,599,732 | 4520 | 3080 | 14,106,636 | 14,221,779 |
| chr13 | 36,012,166 | 35,414,644 | 4480 | 3600 | 140,13,961 | 13,767,350 |
| chr14 | 35,931,519 | 33,969,163 | 4000 | 1800 | 14,052,979 | 13,294,563 |
| chr15 | 35,750,743 | 35,292,775 | 4400 | 3200 | 13,924,384 | 13,776,540 |

| | | | | | | |
|---|---|---|---|---|---|---|
| chr16 | 35,231,784 | 35,722,923 | 2000 | 3000 | 13,763,990 | 13,977,112 |
| chr17 | 35,024,062 | 34,260,779 | 3600 | 3000 | 13,723,061 | 13,378,019 |
| chr18 | 32,836,095 | 32,014,133 | 3000 | 3000 | 12,891,976 | 12,568,077 |
| chr19 | 32,517,511 | 32,398,972 | 3080 | 2800 | 12,718,626 | 12,705,184 |
| chr20 | 32,504,701 | 33,217,526 | 3600 | 3400 | 12,703,219 | 12,964,862 |
| chr21 | 32,232,413 | 32,235,896 | 3600 | 3880 | 12,521622 | 12,566,351 |
| chr22 | 30,694,596 | 29,484,087 | 4040 | 1880 | 11,995,137 | 11,529,814 |
| chr23 | 29,429,680 | 30,284,920 | 3800 | 2640 | 11,489,442 | 11,850,020 |
| chr24 | 27,735,089 | 27,536,296 | 2600 | 3400 | 10,767,244 | 10,669,027 |
| chr25 | 26,155,947 | 26,933,469 | 2720 | 2720 | 10,308,350 | 10,652,242 |

**Table S3: Statistics of repeat sequences annotated in Haplotype 1**

| Type | Number of elements | Length occupied(bp) | % of genome |
|---|---|---|---|
| SINEs | 20,701 | 9,685,118 | 1.03 |
| LINEs | 95,968 | 39,861,338 | 4.24 |
| LTR | 88,887 | 58,530,202 | 6.23 |
| DNA Transposons | 986,164 | 199,681,960 | 21.25 |
| Simple repeats | 357,271 | 25,011,696 | 2.66 |
| Low complexity | 30,066 | 1,632,335 | 0.17 |
| Small RNA | 15,964 | 6,297,372 | 0.47 |
| Unclassified | 678,241 | 132,275,811 | 14.07 |
| Rolling-circles | 28,488 | 19,457,434 | 2.07 |
| Total Repeats Masked | | 506,150,215 | 53.86 |

| Sample | Read Length | Total Sequences | Reads Mapped Hap1 | Reads Mapped Hap2 | Mapping % Hap1 | Mapping % Hap2 |
|---|---|---|---|---|---|---|
| Brain | 148 | 14,569,393,866 | 14,470,079,592 | 14,475,502,985 | 99.32 | 99.36 |
| Gill | 148 | 14,866,778,876 | 14,757,045,591 | 14,749,480,068 | 99.26 | 99.21 |
| Gonad | 148 | 15,792,931,193 | 15,650,843,930 | 15,602,416,417 | 99.10 | 98.79 |
| liver | 148 | 15,180,023,298 | 15,036,318,160 | 14,793,448,652 | 99.05 | 97.45 |
| Muscle | 148 | 14,416,254,932 | 14,279,684,294 | 13,979,419,332 | 99.05 | 96.97 |

| | | | | | | |
|---|---|---|---|---|---|---|
| Skin | 148 | 13,858,527,681 | 13,753,002,576 | 13,853,592,946 | 99.24 | 99.96 |
| Spleen | 148 | 14,035,619,158 | 13,903,999,665 | 13,832,603,441 | 99.06 | 98.55 |

**Table S4: Statistics of repeat sequences annotated in Haplotype 2**

| Type | Number of elements | Length occupied(bp) | % of genome |
|---|---|---|---|
| SINEs | 20,038 | 10,138,453 | 1.09 |
| LINEs | 93,623 | 38,556,326 | 4.15 |
| LTR | 87,308 | 57,276,336 | 6.16 |
| DNA Transposons | 977,301 | 196,769,671 | 21.17 |
| Simple repeats | 350,147 | 24,557,902 | 2.64 |
| Low complexity | 29,831 | 1,637,165 | 0.18 |
| Small RNA | 15,155 | 6,228,729 | 0.67 |
| Unclassified | 668,657 | 130,249,015 | 14.01 |
| Rolling-circles | 27700 | 20,074,520 | 2.16 |
| Total Repeats Masked | | 499,170,338 | 53.71 |

**Table S5: Summary mapping statistics of RNA sequence data used for protein annotation in the *Phoxinus phoxinus* genome**

#### Supplementary Figures

**Figure S1: Heatmap of Haplotype 1 Hi-C assembly with darker blocks indicating higher intensity of sequence interaction**

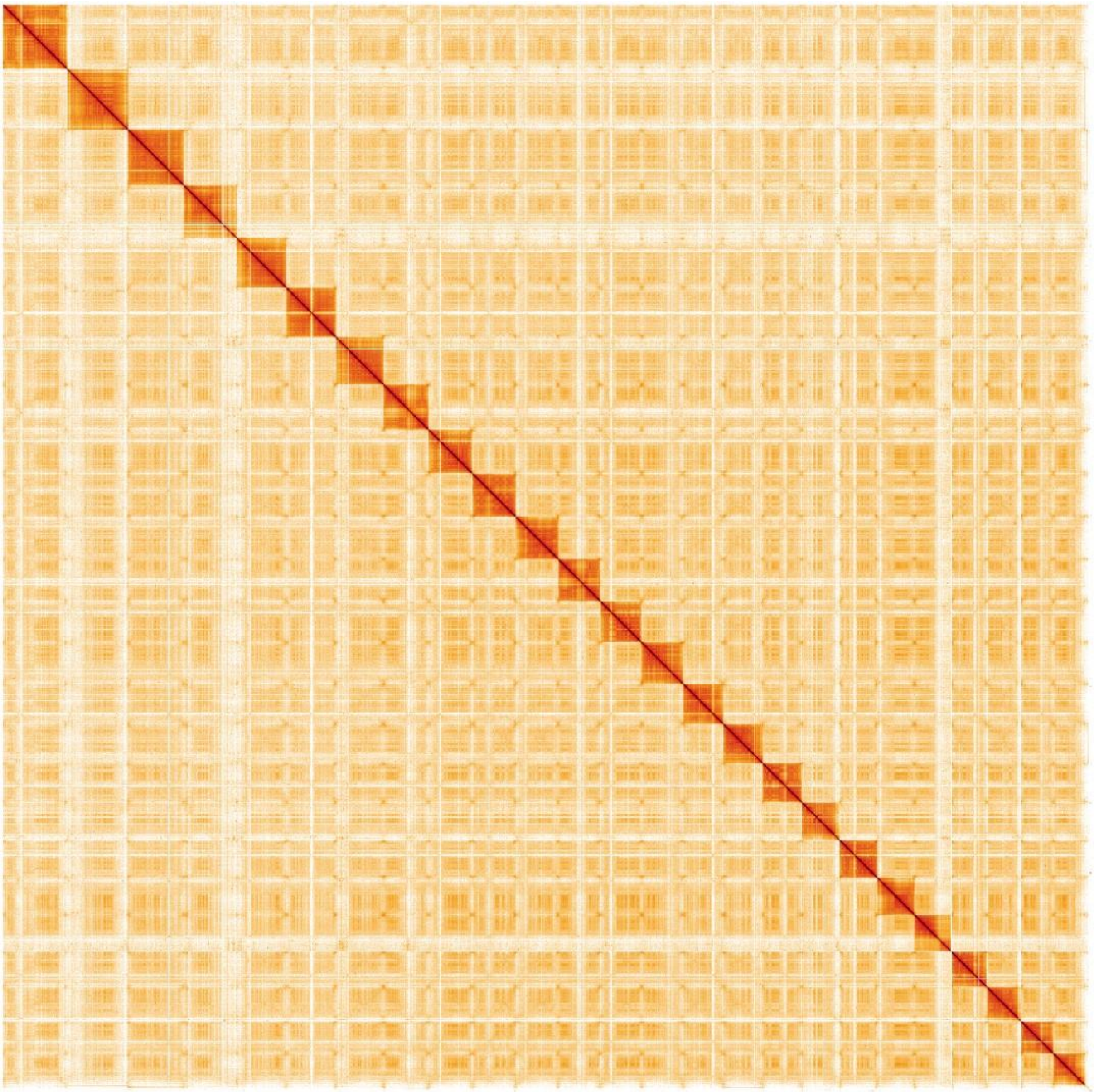

**Figure**

**S2: Heatmap of Haplotype 2 Hi-C assembly with darker blocks indicating higher intensity of sequence interaction**

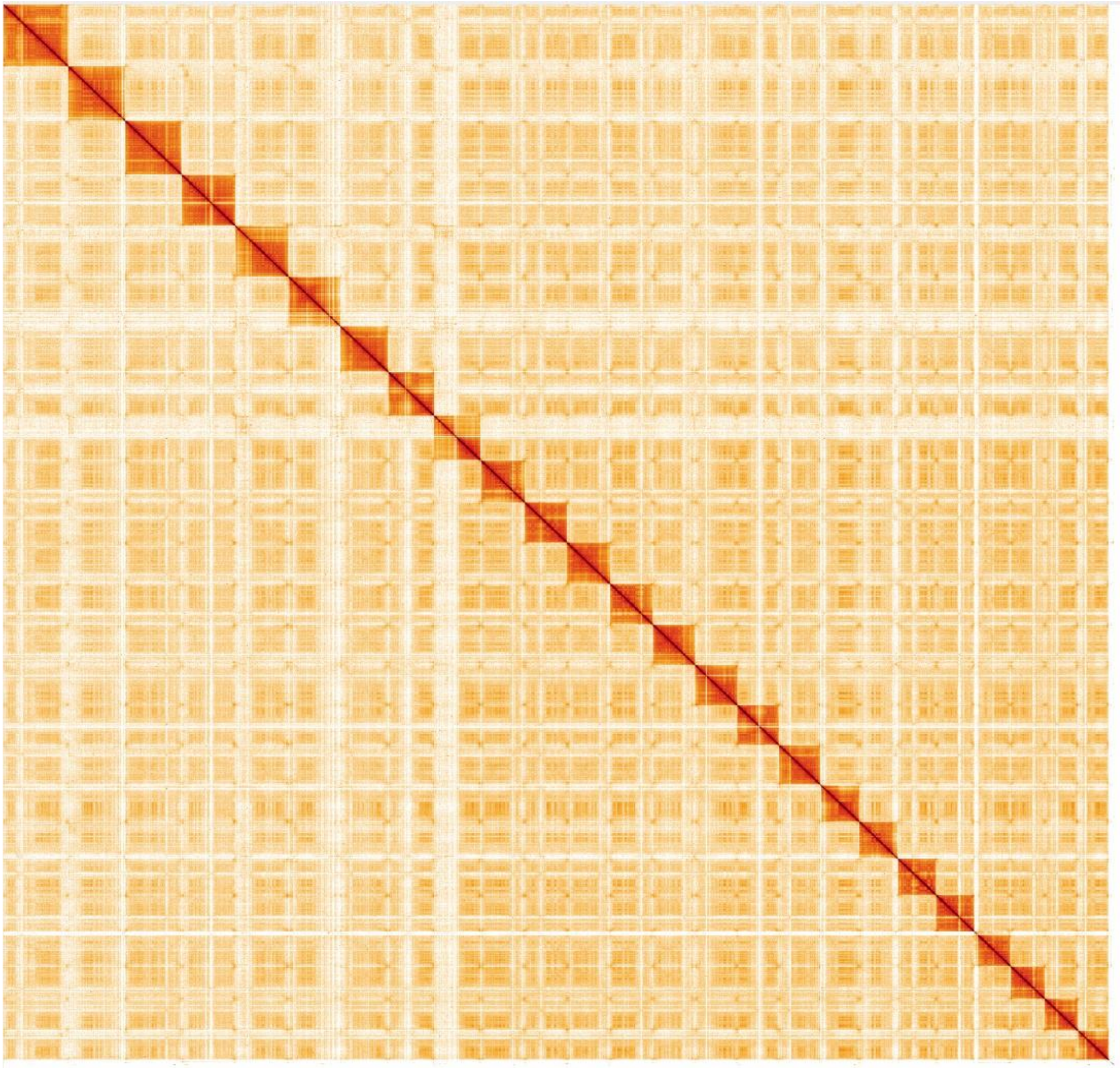

**Figure S3: GenomeScope plots of 19, 23, 25 and 30 mer analysis**

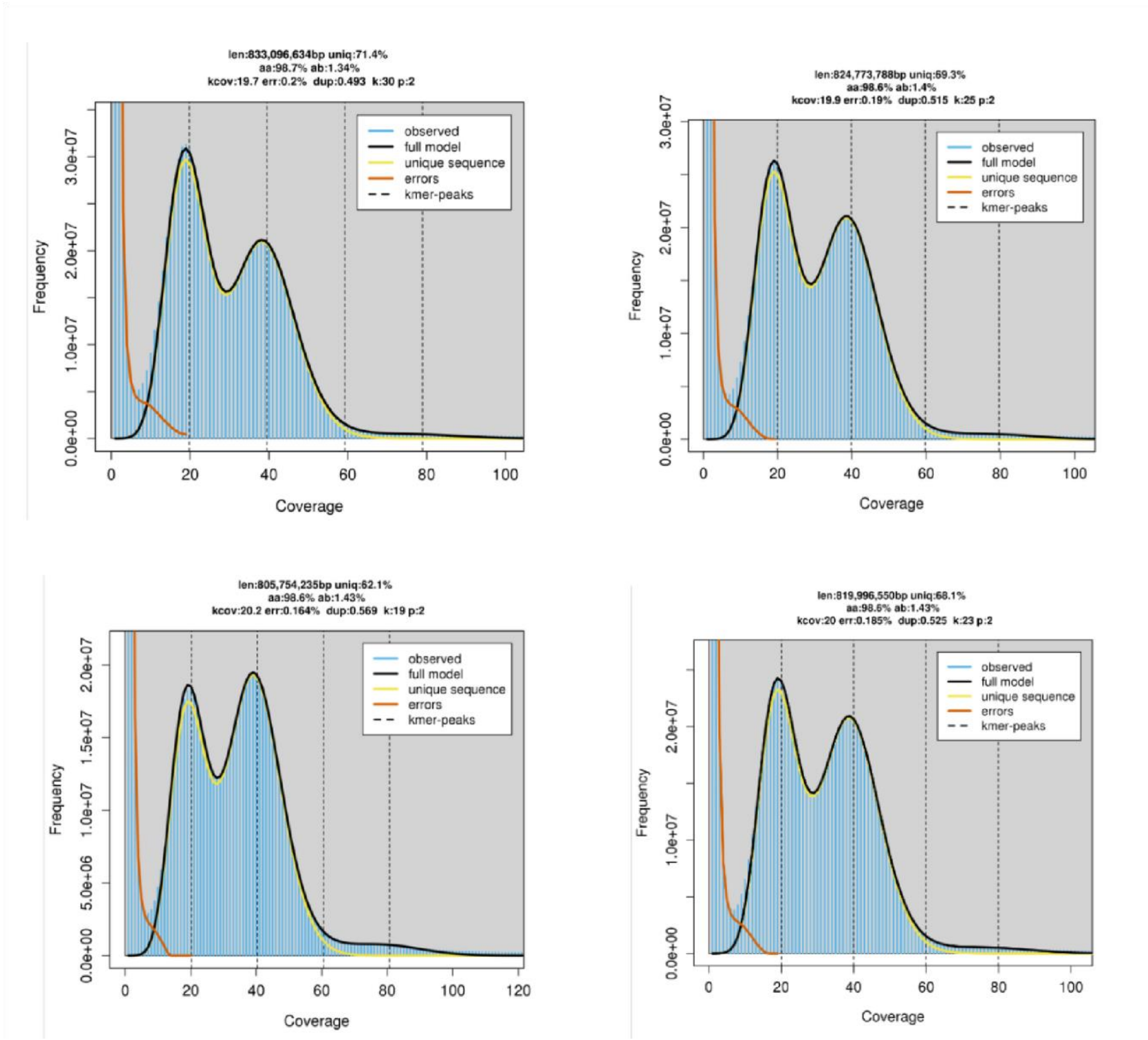


**S4: Repeat Landscape of annotated repeats in Haplotype 2**

**Figure**

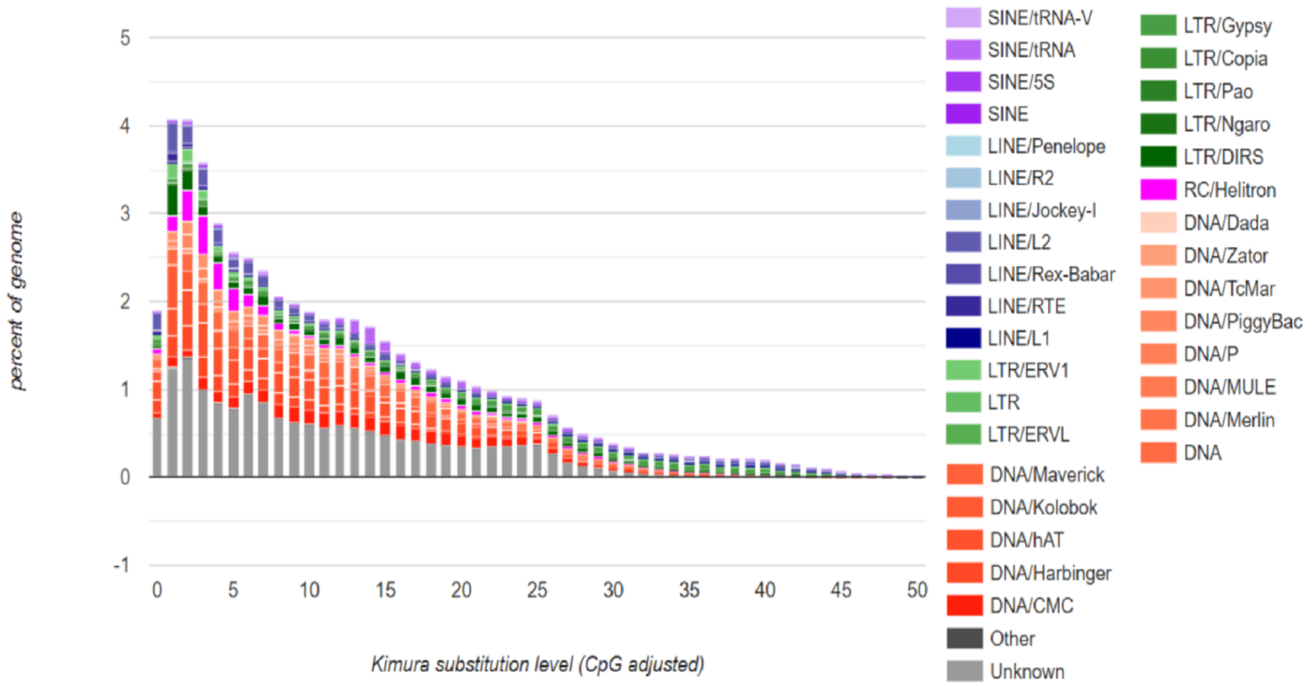
percent of genome
5
4
3
2
1
0
-1
0
5
10
15
20
25
30
35
40
45
50
Kimura substitution level (CpG adjusted)
SINE/tRNA-V
SINE/tRNA
SINE/5S
SINE
LINE/Penelope
LINE/R2
LINE/Jockey-I
LINE/L2
LINE/Rex-Babar
LINE/RTE
LINE/L1
LTR/ERV1
LTR
LTR/ERVL
DNA/Maverick
DNA/Kolobok
DNA/hAT
DNA/Harbinger
DNA/CMC
Other
Unknown
LTR/Gypsy
LTR/Copia
LTR/Pao
LTR/Ngaro
LTR/DIRS
RC/Helitron
DNA/Dada
DNA/Zator
DNA/TcMar
DNA/PiggyBac
DNA/P
DNA/MULE
DNA/Merlin
DNA

**Figure**

**S5: Shared homologs between *Phoxinus phoxinus* and zebrafish, fathead minnow, goldfish, grass carp and common carp.**

A. Venn diagram of shared homologs between multiple fish species and the *Phoxinus phoxinus*.
B. Total number of homologs shared by each species with the Phoxinus phoxinus.

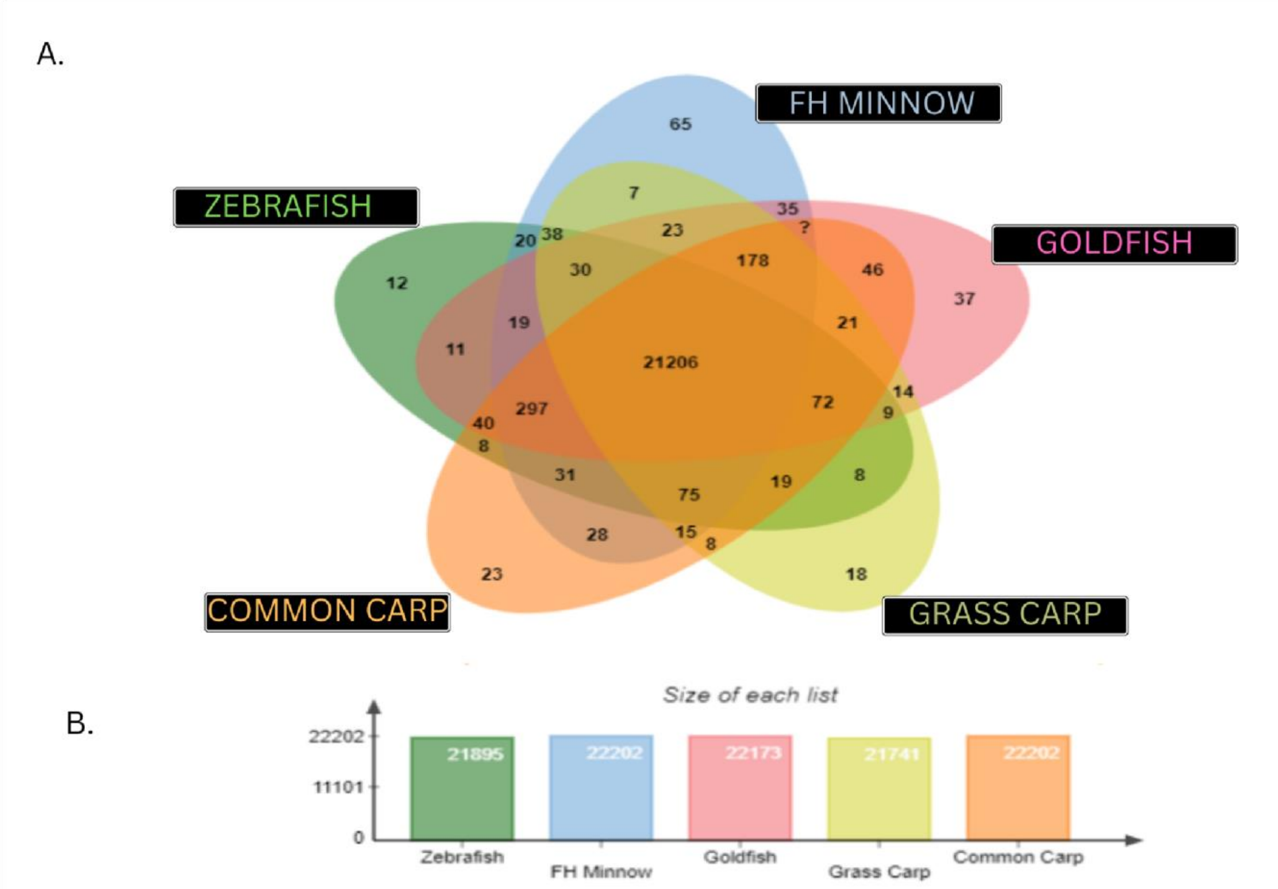


**S6: Gene ontology of genes in Insertion/Deletions**

**Figure**

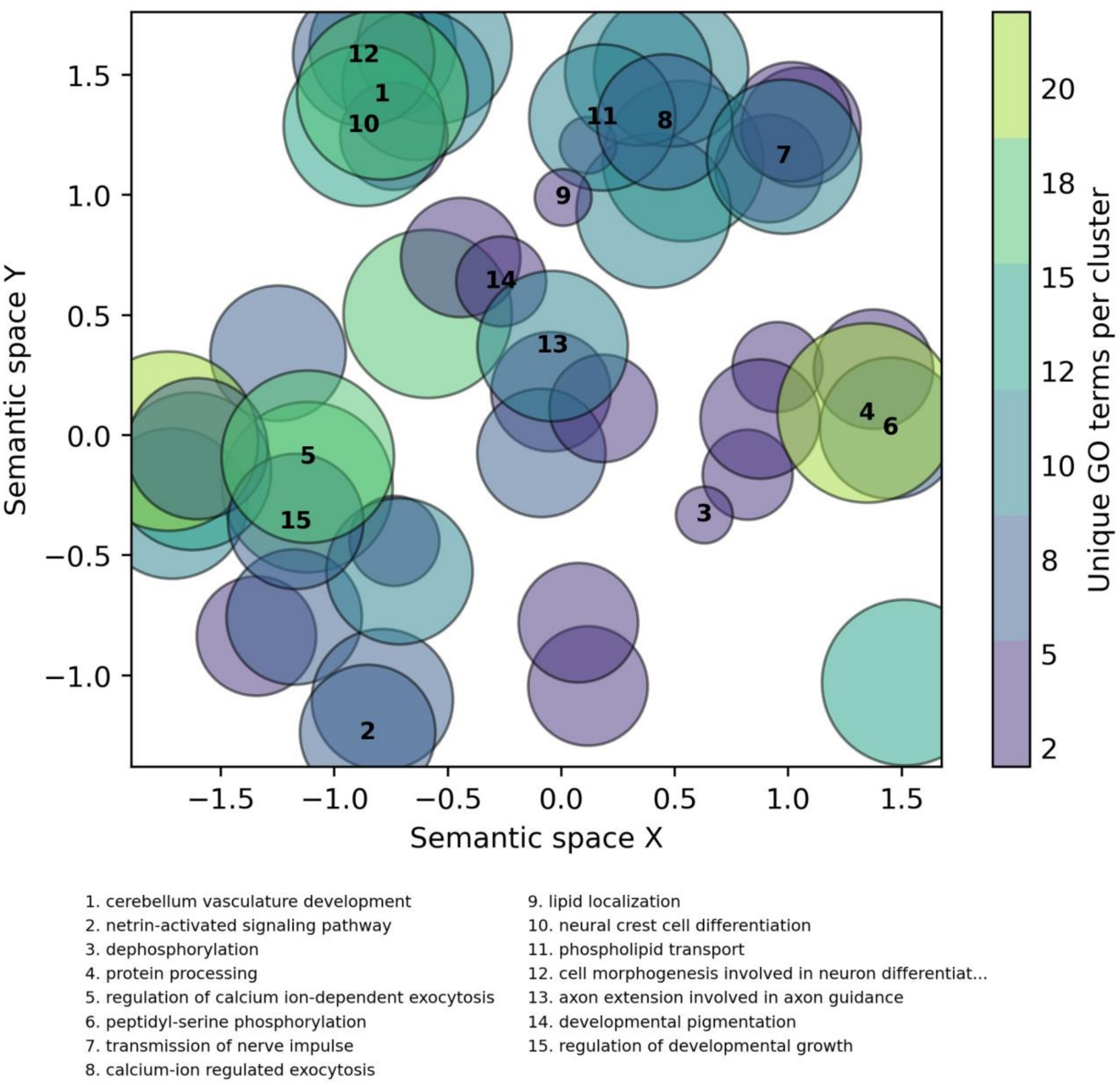


**Figure S7: Gene ontology of genes in inversions**

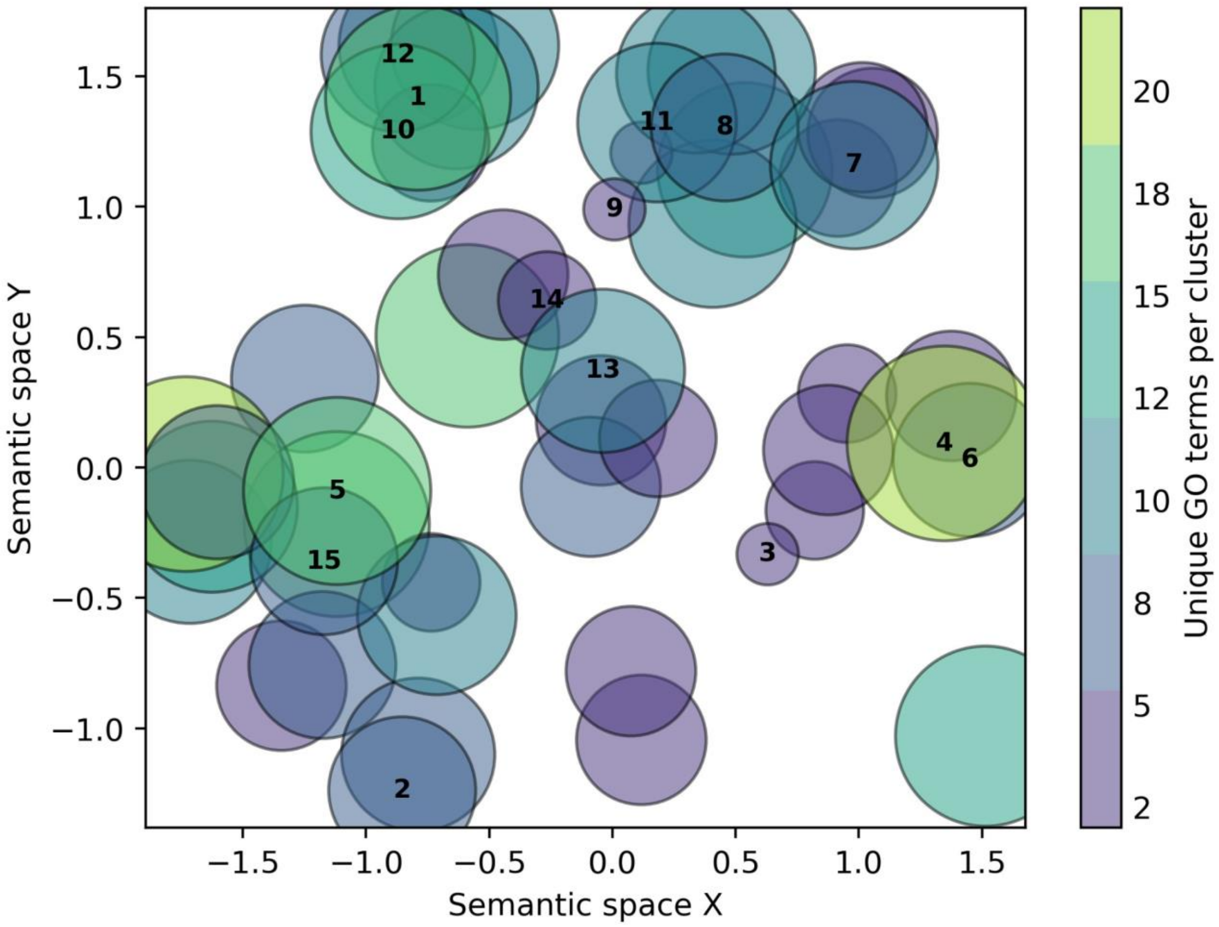


1. cerebellum vasculature development
2. netrin-activated signaling pathway
3. dephosphorylation
4. protein processing
5. regulation of calcium ion-dependent exocytosis
6. peptidyl-serine phosphorylation
7. transmission of nerve impulse
8. calcium-ion regulated exocytosis
9. lipid localization
10. neural crest cell differentiation
11. phospholipid transport
12. cell morphogenesis involved in neuron differentiat...
13. axon extension involved in axon guidance
14. developmental pigmentation
15. regulation of developmental growth

**Figure S8: Expansion/contraction profile including all gene families (significant and non-significant families)**

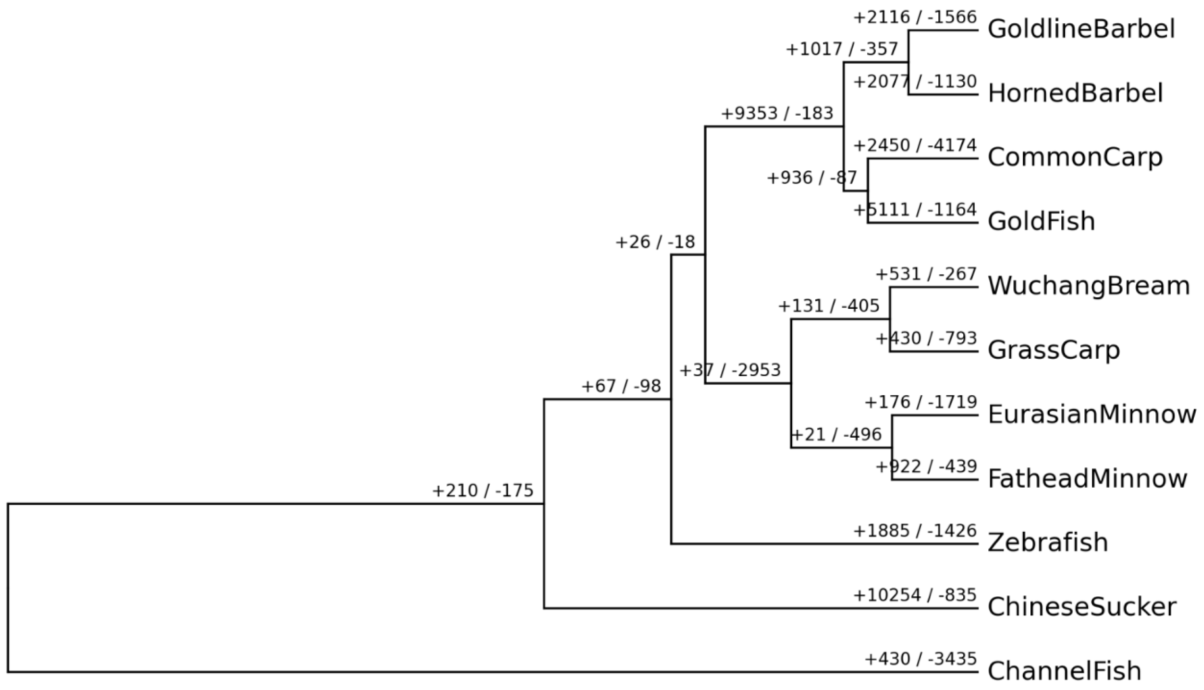


**Figure S9: Pfam domain distribution of expanded genes**

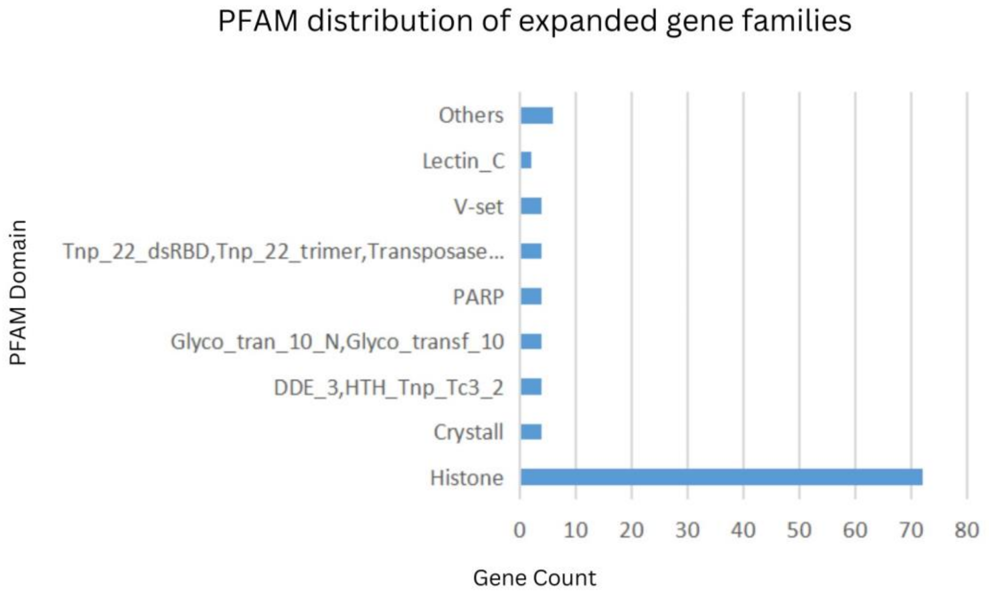

**Figure S10: Pfam domain distribution of contracted genes**

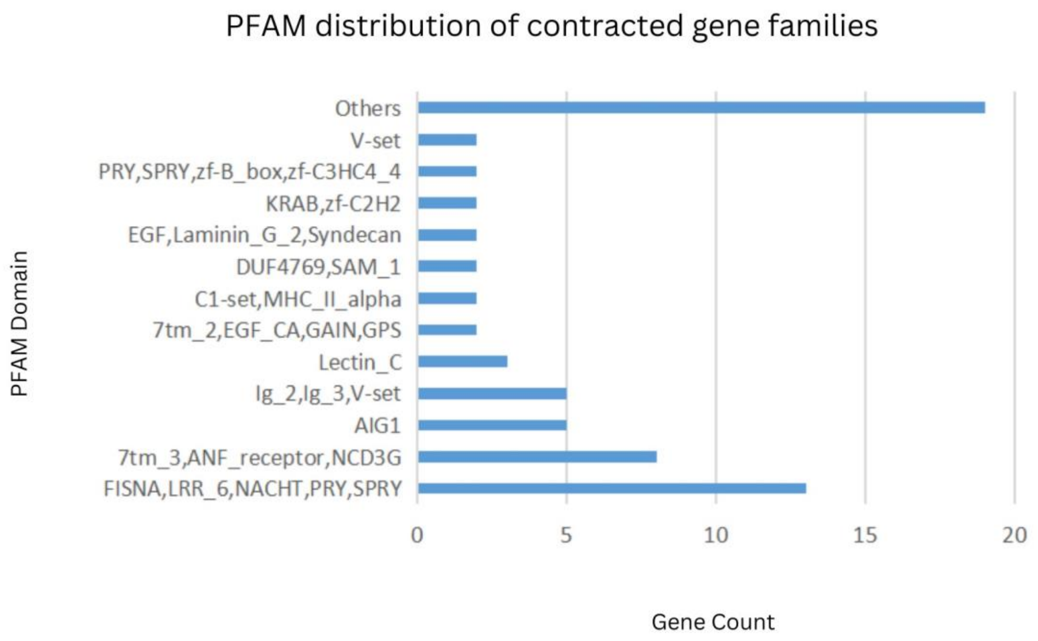
