## Supplemental table S1 for "A chromosome-level, haplotype-resolved genome assembly and annotation for the Eurasian minnow (Leuciscidae: *Phoxinus phoxinus*) provide evidence of haplotype diversity"

**Supplementary Table S1: Mapping Statistics**

**Haplome 1**

| Reads mapped to Hap1 | number of bases | number of contigs | number of windows | number of reads | number of mapped reads | number of mapped reads (%) | number of supplementary alignments | number of supplementary alignments (%) | number of secondary alignments | number of mapped bases | number of sequenced bases |
|---|---|---|---|---|---|---|---|---|---|---|---|
| hifi reads | 939831526 | 101 | 500 | 3462850 | 3449873 | 99.63 | 1535608 | 44.35 | 0 | 36681631436 | 35216974085 |
| hic reads | 939831526 | 101 | 500 | 590535188 | 586166434 | 99.26 | NA | NA | 130988058 | 78595073563 | 78351649598 |
| PacBio raw subreads | 939831526 | 101 | 500 | 66073882 | 66073882 | 100 | 37668671 | 57.01 | 0 | 5.64496E+11 | 5.38756E+11 |

**Haplome 2**

| | | | | | | | | | | | |
|---|---|---|---|---|---|---|---|---|---|---|---|
| hifi reads | 929446345 | 81 | 480 | 3462850 | 3449461 | 99.61 | 1635249 | 47.22 | 0 | 36564140852 | 35102754391 |
| hic reads | 929446345 | 81 | 480 | 590535188 | 585693378 | 99.18 | NA | NA | 131311899 | 78430259433 | 78185010131 |
| PacBio raw subreads | 929446345 | 81 | 480 | 66062482 | 66062482 | 100 | 39728898 | 60.14 | 0 | 5.62804E+11 | 5.37131E+11 |

| number of aligned bases | number of duplicated reads (estimated) | duplication rate | mean mapping quality | GC percentage | general error rate | number of mismatches | number of insertions | mapped reads with insertion percentage | number of deletions | mapped reads with deletion percentage | homopolymer indels | mean coverageData | std coverageData |
|---|---|---|---|---|---|---|---|---|---|---|---|---|---|
| 0 | 1444440 | 6.26 | 52.294 | 38.87 | 0.0725 | 1695914699 | 59080445 | 122.56 | 60163311 | 120.12 | 71.36 | 39.03 | 28.1393 |
| 0 | 288990095 | 41 | 42.4001 | 39.56 | 0.0133 | 811734298 | 75219118 | 11.79 | 48598107 | 7.28 | 34.66 | 83.6268 | 132.7673 |
| 0 | 57094113 | 26.7 | 50.7177 | 37.18 | 0.1518 | 0 | NA | NA | NA | NA | NA | 600.6354 | 378.496 |
| 0 | 1546609 | 6.43 | 52.1922 | 38.86 | 0.0725 | 1695465676 | 59364776 | 124.86 | 60419514 | 122.66 | 70.77 | 39.3397 | 25.4997 |
| 0 | 289940357 | 41.2 | 43.6282 | 39.56 | 0.0135 | 819627461 | 75619157 | 11.84 | 49165891 | 7.36 | 34.67 | 84.3838 | 82.6599 |
| 0 | 58730960 | 26.95 | 50.7296 | 37.17 | 0.1519 | 0 | | 159.72 | | 159.31 | 13.83 | 605.5266 | 353.4762 |
