## Supplemental table S6 for "A chromosome-level, haplotype-resolved genome assembly and annotation for the Eurasian minnow (Leuciscidae: *Phoxinus phoxinus*) provide evidence of haplotype diversity"

**Table S6:** Haplotype 1 protein annotation summary statistics

| | |
|---|---|
| *Statistics with Isoform* | |
| Number of gene | 23397 |
| Number of mrna | 30980 |
| Number of mrnas with utr both sides | 3 |
| Number of mrnas with at least one utr | 61 |
| Number of cds | 30980 |
| Number of exon | 335300 |
| Number of five_prime_utr | 17 |
| Number of intron | 304284 |
| Number of start_codon | 30956 |
| Number of stop_codon | 30967 |
| Number of three_prime_utr | 47 |
| Number of exon in cds | 335264 |
| Number of exon in five_prime_utr | 17 |
| Number of exon in three_prime_utr | 47 |
| Number of intron in cds | 304284 |
| Number of intron in exon | 304320 |
| Number of intron in intron | 275263 |
| Number gene overlapping | 105 |
| Number of single exon gene | 1706 |
| Number of single exon mrna | 1959 |
| mean mrnas per gene | 1.3 |
| mean cdss per mrna | 1 |
| mean exons per mrna | 10.8 |
| mean five_prime_utrs per mrna | 0 |
| mean introns per mrna | 9.8 |
| mean start_codons per mrna | 1 |
| mean stop_codons per mrna | 1 |
| mean three_prime_utrs per mrna | 0 |
| mean exons per cds | 10.8 |
| mean exons per five_prime_utr | 1 |
| mean exons per three_prime_utr | 1 |
| mean introns in cdss per mrna | 9.8 |
| mean introns in exons per mrna | 9.8 |
| mean introns in introns per mrna | 8.9 |
| Total gene length | 368122196 |
| Total mrna length | 574698649 |
| Total cds length | 54378219 |
| Total exon length | 54505112 |
| Total five_prime_utr length | 52039 |
| Total intron length | 520130354 |
| Total start_codon length | 92801 |
| Total stop_codon length | 92864 |
| Total three_prime_utr length | 74854 |
| Total intron length per cds | 520130354 |
| Total intron length per exon | 520193537 |

| | |
|---|---|
| Total intron length per intron | 39999524 |
| mean gene length | 15733 |
| mean mrna length | 18550 |
| mean cds length | 1755 |
| mean exon length | 162 |
| mean five_prime_utr length | 3061 |
| mean intron length | 1709 |
| mean start_codon length | 2 |
| mean stop_codon length | 2 |
| mean three_prime_utr length | 1592 |
| mean cds piece length | 162 |
| mean five_prime_utr piece length | 3061 |
| mean three_prime_utr piece length | 1592 |
| mean intron in cds length | 1709 |
| mean intron in exon length | 1709 |
| mean intron in intron length | 145 |
| % of genome covered by gene | 39.2 |
| % of genome covered by mrna | 61.1 |
| % of genome covered by cds | 5.8 |
| % of genome covered by exon | 5.8 |
| % of genome covered by five_prime_utr | 0 |
| % of genome covered by intron | 55.3 |
| % of genome covered by start_codon | 0 |
| % of genome covered by stop_codon | 0 |
| % of genome covered by three_prime_utr | 0 |
| % of genome covered by intron from cds | 55.3 |
| % of genome covered by intron from exon | 55.3 |
| % of genome covered by intron from intron | 4.3 |
| Longest gene | 531808 |
| Longest mrna | 531808 |
| Longest cds | 76851 |
| Longest exon | 24342 |
| Longest five_prime_utr | 11305 |
| Longest intron | 528063 |
| Longest start_codon | 3 |
| Longest stop_codon | 3 |
| Longest three_prime_utr | 24186 |
| Longest cds piece | 17172 |
| Longest five_prime_utr piece | 11305 |
| Longest three_prime_utr piece | 24186 |
| Longest intron into cds part | 528063 |
| Longest intron into exon part | 528063 |
| Longest intron into intron part | 17172 |
| Shortest gene | 31 |
| Shortest mrna | 31 |
| Shortest cds | 9 |
| Shortest exon | 1 |

| | |
|---|---|
| Shortest five_prime_utr | 60 |
| Shortest intron | 21 |
| Shortest start_codon | 1 |
| Shortest stop_codon | 1 |
| Shortest three_prime_utr | 3 |
| Shortest cds piece | 1 |
| Shortest five_prime_utr piece | 60 |
| Shortest three_prime_utr piece | 3 |
| Shortest intron into cds part | 21 |
| Shortest intron into exon part | 21 |
| Shortest intron into intron part | 7 |

*Statistics without Isoform*

| | |
|---|---|
| Number of gene | 23397 |
| Number of mrna | 23397 |
| Number of mrnas with utr both sides | 1 |
| Number of mrnas with at least one utr | 31 |
| Number of cds | 23397 |
| Number of exon | 221676 |
| Number of five_prime_utr | 14 |
| Number of intron | 198270 |
| Number of start_codon | 23377 |
| Number of stop_codon | 23386 |
| Number of three_prime_utr | 18 |
| Number of exon in cds | 221667 |
| Number of exon in five_prime_utr | 14 |
| Number of exon in three_prime_utr | 18 |
| Number of intron in cds | 198270 |
| Number of intron in exon | 198279 |
| Number of intron in intron | 176589 |
| Number gene overlapping | 102 |
| Number of single exon gene | 1716 |
| Number of single exon mrna | 1716 |
| mean mrnas per gene | 1 |
| mean cdss per mrna | 1 |
| mean exons per mrna | 9.5 |
| mean five_prime_utrs per mrna | 0 |
| mean introns per mrna | 8.5 |
| mean start_codons per mrna | 1 |
| mean stop_codons per mrna | 1 |
| mean three_prime_utrs per mrna | 0 |
| mean exons per cds | 9.5 |
| mean exons per five_prime_utr | 1 |
| mean exons per three_prime_utr | 1 |
| mean introns in cdss per mrna | 8.5 |
| mean introns in exons per mrna | 8.5 |

| | |
|---|---|
| mean introns in introns per mrna | 7.5 |
| Total gene length | 368122196 |
| Total mrna length | 363173395 |
| Total cds length | 37177004 |
| Total exon length | 37296849 |
| Total five_prime_utr length | 50121 |
| Total intron length | 325838658 |
| Total start_codon length | 70079 |
| Total stop_codon length | 70131 |
| Total three_prime_utr length | 69724 |
| Total intron length per cds | 325838658 |
| Total intron length per exon | 325876546 |
| Total intron length per intron | 25850875 |
| mean gene length | 15733 |
| mean mrna length | 15522 |
| mean cds length | 1588 |
| mean exon length | 168 |
| mean five_prime_utr length | 3580 |
| mean intron length | 1643 |
| mean start_codon length | 2 |
| mean stop_codon length | 2 |
| mean three_prime_utr length | 3873 |
| mean cds piece length | 167 |
| mean five_prime_utr piece length | 3580 |
| mean three_prime_utr piece length | 3873 |
| mean intron in cds length | 1643 |
| mean intron in exon length | 1643 |
| mean intron in intron length | 146 |
| % of genome covered by gene | 39.2 |
| % of genome covered by mrna | 38.6 |
| % of genome covered by cds | 4 |
| % of genome covered by exon | 4 |
| % of genome covered by five_prime_utr | 0 |
| % of genome covered by intron | 34.7 |
| % of genome covered by start_codon | 0 |
| % of genome covered by stop_codon | 0 |
| % of genome covered by three_prime_utr | 0 |
| % of genome covered by intron from cds | 34.7 |
| % of genome covered by intron from exon | 34.7 |
| % of genome covered by intron from intron | 2.8 |
| Longest gene | 531808 |
| Longest mrna | 528270 |
| Longest cds | 76851 |
| Longest exon | 24342 |
| Longest five_prime_utr | 11305 |
| Longest intron | 528063 |
| Longest start_codon | 3 |

| | |
|---|---|
| Longest stop_codon | 3 |
| Longest three_prime_utr | 24186 |
| Longest cds piece | 17172 |
| Longest five_prime_utr piece | 11305 |
| Longest three_prime_utr piece | 24186 |
| Longest intron into cds part | 528063 |
| Longest intron into exon part | 528063 |
| Longest intron into intron part | 17172 |
| Shortest gene | 31 |
| Shortest mrna | 31 |
| Shortest cds | 9 |
| Shortest exon | 1 |
| Shortest five_prime_utr | 60 |
| Shortest intron | 24 |
| Shortest start_codon | 1 |
| Shortest stop_codon | 1 |
| Shortest three_prime_utr | 3 |
| Shortest cds piece | 1 |
| Shortest five_prime_utr piece | 60 |
| Shortest three_prime_utr piece | 3 |
| Shortest intron into cds part | 24 |
| Shortest intron into exon part | 24 |
| Shortest intron into intron part | 7 |

--------------------------------------------------------------------------------

--------------------------------------------------------------------------------
