## Supplemental table S7 for "A chromosome-level, haplotype-resolved genome assembly and annotation for the Eurasian minnow (Leuciscidae: *Phoxinus phoxinus*) provide evidence of haplotype diversity"

**Table S7:** Haplotype 2 protein annotation summary statistics

| | |
|---|---|
| *Statistics with Isoform* | |
| Number of gene | 23191 |
| Number of mrna | 29614 |
| Number of cds | 29614 |
| Number of exon | 321995 |
| Number of intron | 292381 |
| Number of start_codon | 29608 |
| Number of stop_codon | 29612 |
| Number of exon in cds | 321995 |
| Number of intron in cds | 292381 |
| Number of intron in exon | 292381 |
| Number of intron in intron | 264699 |
| Number gene overlapping | 81 |
| Number of single exon gene | 1726 |
| Number of single exon mrna | 1932 |
| mean mrnas per gene | 1.3 |
| mean cdss per mrna | 1 |
| mean exons per mrna | 10.9 |
| mean introns per mrna | 9.9 |
| mean start_codons per mrna | 1 |
| mean stop_codons per mrna | 1 |
| mean exons per cds | 10.9 |
| mean introns in cdss per mrna | 9.9 |
| mean introns in exons per mrna | 9.9 |
| mean introns in introns per mrna | 8.9 |
| Total gene length | 353358641 |
| Total mrna length | 526393142 |
| Total cds length | 52759456 |
| Total exon length | 52759456 |
| Total intron length | 473633686 |
| Total start_codon length | 88768 |
| Total stop_codon length | 88799 |
| Total intron length per cds | 473633686 |
| Total intron length per exon | 473633686 |
| Total intron length per intron | 38795161 |
| mean gene length | 15236 |
| mean mrna length | 17775 |
| mean cds length | 1781 |
| mean exon length | 163 |
| mean intron length | 1619 |
| mean start_codon length | 2 |
| mean stop_codon length | 2 |
| mean cds piece length | 163 |
| mean intron in cds length | 1619 |
| mean intron in exon length | 1619 |

| | |
|---|---|
| mean intron in intron length | 146 |
| % of genome covered by gene | 38 |
| % of genome covered by mrna | 56.6 |
| % of genome covered by cds | 5.7 |
| % of genome covered by exon | 5.7 |
| % of genome covered by intron | 51 |
| % of genome covered by start_codon | 0 |
| % of genome covered by stop_codon | 0 |
| % of genome covered by intron from c | 51 |
| % of genome covered by intron from e | 51 |
| % of genome covered by intron from ii | 4.2 |
| Longest gene | 569087 |
| Longest mrna | 569087 |
| Longest cds | 76851 |
| Longest exon | 17172 |
| Longest intron | 315204 |
| Longest start_codon | 3 |
| Longest stop_codon | 3 |
| Longest cds piece | 17172 |
| Longest intron into cds part | 315204 |
| Longest intron into exon part | 315204 |
| Longest intron into intron part | 17172 |
| Shortest gene | 31 |
| Shortest mrna | 31 |
| Shortest cds | 9 |
| Shortest exon | 1 |
| Shortest intron | 21 |
| Shortest start_codon | 1 |
| Shortest stop_codon | 1 |
| Shortest cds piece | 1 |
| Shortest intron into cds part | 21 |
| Shortest intron into exon part | 21 |
| Shortest intron into intron part | 8 |

*Statistics without Isoform*

| | |
|---|---|
| Number of gene | 23191 |
| Number of mrna | 23191 |
| Number of cds | 23191 |
| Number of exon | 220419 |
| Number of intron | 197228 |
| Number of start_codon | 23186 |
| Number of stop_codon | 23189 |
| Number of exon in cds | 220419 |
| Number of intron in cds | 197228 |
| Number of intron in exon | 197228 |
| Number of intron in intron | 175771 |

| | |
|---|---|
| Number gene overlapping | 76 |
| Number of single exon gene | 1734 |
| Number of single exon mrna | 1734 |
| mean mrnas per gene | 1 |
| mean cdss per mrna | 1 |
| mean exons per mrna | 9.5 |
| mean introns per mrna | 8.5 |
| mean start_codons per mrna | 1 |
| mean stop_codons per mrna | 1 |
| mean exons per cds | 9.5 |
| mean introns in cdss per mrna | 8.5 |
| mean introns in exons per mrna | 8.5 |
| mean introns in introns per mrna | 7.6 |
| Total gene length | 353358641 |
| Total mrna length | 348329171 |
| Total cds length | 37081292 |
| Total exon length | 37081292 |
| Total intron length | 311247879 |
| Total start_codon length | 69506 |
| Total stop_codon length | 69541 |
| Total intron length per cds | 311247879 |
| Total intron length per exon | 311247879 |
| Total intron length per intron | 25782302 |
| mean gene length | 15236 |
| mean mrna length | 15020 |
| mean cds length | 1598 |
| mean exon length | 168 |
| mean intron length | 1578 |
| mean start_codon length | 2 |
| mean stop_codon length | 2 |
| mean cds piece length | 168 |
| mean intron in cds length | 1578 |
| mean intron in exon length | 1578 |
| mean intron in intron length | 146 |
| % of genome covered by gene | 38 |
| % of genome covered by mrna | 37.5 |
| % of genome covered by cds | 4 |
| % of genome covered by exon | 4 |
| % of genome covered by intron | 33.5 |
| % of genome covered by start_codon | 0 |
| % of genome covered by stop_codon | 0 |
| % of genome covered by intron from c | 33.5 |
| % of genome covered by intron from e | 33.5 |
| % of genome covered by intron from ir | 2.8 |
| Longest gene | 569087 |
| Longest mrna | 569087 |
| Longest cds | 76851 |

| | |
|---|---|
| Longest exon | 17172 |
| Longest intron | 300229 |
| Longest start_codon | 3 |
| Longest stop_codon | 3 |
| Longest cds piece | 17172 |
| Longest intron into cds part | 300229 |
| Longest intron into exon part | 300229 |
| Longest intron into intron part | 17172 |
| Shortest gene | 31 |
| Shortest mrna | 31 |
| Shortest cds | 9 |
| Shortest exon | 1 |
| Shortest intron | 21 |
| Shortest start_codon | 1 |
| Shortest stop_codon | 1 |
| Shortest cds piece | 1 |
| Shortest intron into cds part | 21 |
| Shortest intron into exon part | 21 |
| Shortest intron into intron part | 8 |

--------------------------------------------------------------------------------
