## Supplemental table S8 for "A chromosome-level, haplotype-resolved genome assembly and annotation for the Eurasian minnow (Leuciscidae: *Phoxinus phoxinus*) provide evidence of haplotype diversity"

***Table S8:*** *Genes present in Insertion/Deletions between both Halpomes*

| Gene Name | Gene ID |
|---|---|
| Serine arginine repetitive matrix protein 1-like | - |
| 5-hydroxytryptamine receptor 7-like | - |
| - | - |
| Fibronectin type 3 domain | - |
| Serine threonine-protein kinase NIM1-like | - |
| Si dkey-122a22.2 | - |
| Nucleophosmin 1b (nucleolar phosphoprotein B23, numatrin) | - |
| BTB POZ domain-containing protein KCTD8-like | - |
| spliceosomal complex disassembly | - |
| Parathyroid hormone 1 receptor b | - |
| Arrestin 3a, retinal (X-arrestin) | - |
| THAP | - |
| Mucin-2-like | - |
| ATP-binding cassette, sub-family C (CFTR MRP), member 3 | ABCC3 |
| Ankyrin repeat and BTB POZ | ABTB1 |
| Adenosine deaminase, RNA-specific, B2 (RED2 homolog rat) | ADARB2 |
| Belongs to the adenylyl cyclase class-4 guanylyl cyclase family | ADCY1 |
| Catalyzes the exchange of an acyl for a long-chain alkyl group and the | AGPS |
| Ankyrin repeat | ANKRD44 |
| Ankyrin repeat | ANKRD52 |
| Anthrax toxin receptor | ANTXR2 |
| Amyloid beta (A4) | APP |
| ArfGAP with RhoGAP domain, ankyrin repeat and PH domain 3 | ARAP3 |
| Rho GTPase activating protein 22 | ARHGAP22 |
| Rho GTPase activating protein 42 | ARHGAP42 |
| Arylsulfatase family, member K | ARSK |
| Belongs to the amiloride-sensitive sodium channel (TC 1.A.6) family | ASIC2 |
| Autophagy related 2A | ATG2A |
| transcriptional regulator | ATRX |
| Breast cancer anti-estrogen resistance | BCAR3 |

| | |
|---|---|
| Apoptosis facilitator | BCL2L14 |
| Forms chloride channels | BEST2 |
| complement | C3 |
| complement | C3 |
| Si dkey-8k3.2 | c4 |
| Calcium channel, voltage-dependent, beta 4b subunit | CACNB4 |
| Calcium channel, voltage-dependent, gamma subunit 8a | CACNG8 |
| Cell adhesion molecule | CADM1 |
| Calcium calmodulin-dependent protein kinase | CAMK2D |
| Belongs to the peptidase C2 family | CAPN3 |
| Formin Homology 2 Domain | CBLN4 |
| Coiled-coil domain containing 132 | CCDC132 |
| Cell division cycle 40 homolog (S. cerevisiae) | CDC40 |
| Cadherin 12, type 2 (N-cadherin 2) | CDH12 |
| Cugbp, Elav-like family member | CELF5 |
| Belongs to the CDP-alcohol phosphatidyltransferase class-I family | CEPT1 |
| ATP-binding cassette, sub-family C (CFTR MRP), member | cft-1 |
| Chloride channel 6 | CLCN6 |
| cyclic nucleotide gated channel beta 3 | CNGB3 |
| Procollagen, type V, alpha 1 | COL5A1 |
| Copine VIII | CPNE8 |
| CTD (carboxy-terminal domain, RNA polymerase II, polypeptide A) small | CTDSPL |
| Cytochrome P450, family 27, subfamily A, polypeptide | CYP27A1 |
| cytochrome P450 | CYP2B6 |
| dachshund | DACH1 |
| Dachsous 1b (Drosophila) | DCHS1 |
| large homolog | DLG1 |
| Double C2-like domain-containing protein beta | DOC2B |
| dedicator of cytokinesis | DOCK11 |
| Involved in pyrimidine base degradation. Catalyzes the reduction of uracil and | DPYD |
| Dymeclin | DYM |
| Transcription factor | E2F6 |
| Endothelin converting enzyme 2 | ECE2 |

| | |
|---|---|
| EGF-like-domain, multiple 7 | EGFL7 |
| RNA polymerase II | ELL |
| Elongation protein 4 homolog (S. cerevisiae) | ELP4 |
| Echinoderm microtubule associated protein like 4 | EML4 |
| Ecto-NOX disulfide-thiol exchanger | ENOX1 |
| V-erb-a erythroblastic leukemia viral oncogene homolog 4b (avian) | ERBB4 |
| Family with sequence similarity 19 (chemokine (C-C motif)-like), member A3 | FAM19A3 |
| Formin homology 2 domain containing 3 | FHOD3 |
| Filamin A, alpha (actin binding protein 280) | FLNA |
| Mitochondrial GTPase that catalyzes the GTP-dependent ribosomal | GFM1 |
| Glutamine-fructose-6-phosphate transaminase 1 | GFPT1 |
| Belongs to the Glu Leu Phe Val dehydrogenases family | GLUD1 |
| Glutamate receptor, ionotropic | GRID1 |
| Glutamate receptor ionotropic, kainate 3 | GRIK3 |
| glutamate receptor | GRM4 |
| Glycogen synthase kinase 3 alpha b | GSK3A |
| HECT and RLD domain containing E3 ubiquitin protein ligase 4 | HERC4 |
| ubiquitin-like domain member 1 | HERPUD1 |
| Hydroxysteroid dehydrogenase like 2 | HSDL2 |
| 5-hydroxytryptamine (serotonin) receptor 4, G protein-coupled | HTR4 |
| Immunoglobulin superfamily member 11 | IGSF11 |
| superfamily, member | IGSF21 |
| Immunoglobulin superfamily member | IGSF3 |
| Type II inositol 3,4-bisphosphate | INPP4B |
| integrin | ITGB6 |
| voltage-gated potassium channel activity | KCNH6 |
| PHD finger protein 24 | KIAA1045 |
| leucyl-tRNA synthetase 2, mitochondrial | LARS2 |
| Prolyl 3-hydroxylase 2 | LEPREL1 |
| Lethal giant larvae homolog 2 (Drosophila) | LLGL2 |
| LPS-responsive vesicle trafficking, beach and anchor containing | LRBA |
| lipoprotein receptor-related protein | LRP1 |
| lipoprotein receptor-related protein | LRP3 |

| | |
|---|---|
| Latent transforming growth factor beta binding protein | LTBP4 |
| guanylate kinase, WW and PDZ | MAGI1 |
| Mitogen-activated protein kinase kinase kinase | MAP3K10 |
| Mitogen-activated protein kinase kinase kinase | MAP3K3 |
| muscleblind-like | MBNL2 |
| Transformed 3T3 cell double minute 2 homolog (mouse) | MDM2 |
| Malic enzyme 1, NADP( )-dependent, cytosolic | ME1 |
| Alpha-1,6-mannosylglycoprotein 6-beta-N-acetylglucosaminyltransferase | MGAT5B |
| Megakaryoblastic leukemia (translocation) | MKL1 |
| protein-like | MLXIPL |
| Musashi homolog | MSI2 |
| Matrix-remodelling associated 8a | MXRA8 |
| V-myb avian myeloblastosis viral oncogene homolog-like 2b | MYBL2 |
| Belongs to the TRAFAC class myosin-kinesin ATPase superfamily. Myosin family | MYO1C |
| Belongs to the TRAFAC class myosin-kinesin ATPase superfamily. Myosin family | MYO3B |
| Si ch1073-219n12.1 | nab-1 |
| N-ethylmaleimide-sensitive factor attachment protein, gamma | NAPG |
| Naked cuticle homolog | NKD2 |
| neuroligin 2b | NLGN2 |
| NLRP3 inflammasome complex assembly | NLRP12 |
| Nephronectin | NPNT |
| Nudix (nucleoside diphosphate linked moiety X)-type motif 15 | nudt15 |
| Olfactomedin 2a | OLFM2 |
| P21 protein (Cdc42 Rac)-activated kinase 3 | PAK3 |
| Presenilin associated, rhomboid-like a | PARL |
| Proprotein convertase subtilisin kexin type 1 | PCSK1 |
| PDZ domain containing 8 | PDZD8 |
| Platelet endothelial cell adhesion | PECAM1 |
| Peptidase D | PEPD |
| Progesterone immunomodulatory binding factor 1 | PIBF1 |
| PITPNM family member 3 | PITPNM3 |
| Phospholipase C, delta 3b | PLCD3 |
| Pleckstrin homology domain containing, family G (with RhoGef domain) | PLEKHG5 |

| | |
|---|---|
| DNA-dependent RNA polymerase catalyzes the transcription of DNA into RNA | POLR1A |
| regulatory subunit B | PPP2R3C |
| Protein kinase C, alpha | PRKCA |
| Belongs to the peptidase S1 family | PRSS8 |
| Pleckstrin and Sec7 domain containing 3, like | PSD3 |
| Proteasome (prosome, macropain) 26S subunit, non-ATPase, 4b | PSMD4 |
| Catalyzes the third of the four reactions of the long- chain fatty acids | PTPLB |
| Paxillin a | PXN |
| Ras-related C3 botulinum toxin substrate 1 (rho family, small GTP binding | RAC1 |
| Rap guanine nucleotide exchange factor | RAPGEF1 |
| RNA binding motif protein 18 | RBM18 |
| RAB6A GEF complex partner 1 | RIC1 |
| E3 ubiquitin-protein ligase | RNF123 |
| Zinc finger, C3HC4 type (RING finger) | RNF169 |
| RAR-related orphan receptor C | RORC |
| Signal peptide, CUB | SCUBE1 |
| Sidekick cell adhesion molecule | SDK1 |
| Sema domain, seven thrombospondin repeats (type 1 and type 1-like), | SEMA5A |
| Solute carrier family 2 (facilitated glucose transporter), member 13 | SLC2A13 |
| Belongs to the sodium neurotransmitter symporter (SNF) (TC 2.A.22) family | SLC6A13 |
| Solute carrier family 9, subfamily A (NHE1, cation proton antiporter 1), | SLC9A1 |
| Sphingomyelin phosphodiesterase | SMPD3 |
| Sphingosine kinase | SPHK1 |
| spinster homolog 2 | SPNS2 |
| Storkhead-box protein 2 | STOX2 |
| Tyrosine aminotransferase | TAT |
| Tektin 2 (testicular) | TEKT2 |
| Testis-expressed sequence 264 | TEX264 |
| Transcription factor | TFE3 |
| THO complex | THOC2 |
| Thrombospondin, type I, domain containing | THSD7B |
| Transmembrane protein 132D-like | TMEM132C |
| Topoisomerase I binding, arginine serine-rich | TOPORS |

| | |
|---|---|
| Thymocyte selection-associated high mobility group box | TOX |
| Trio Rho guanine nucleotide exchange factor a | TRIO |
| tetratricopeptide repeat | TTC14 |
| tetratricopeptide repeat | TTC27 |
| Ttk protein kinase | TTK |
| tweety homolog 2 | TTYH2 |
| Ubiquitin protein ligase E3B | UBE3B |
| Unc-5 homolog C (C. elegans) | UNC5C |
| Unc-5 homolog Da (C. elegans) | unc5d |
| May play a role in anchoring the cytoskeleton to the plasma membrane | UTRN |
| Vitamin K epoxide reductase complex, subunit 1-like | VKORC1L1 |
| protein family member 1 | WASF1 |
| Xanthine dehydrogenase | XDH |
| Xylulokinase homolog (H. influenzae) | XYLB |
| zinc finger and BTB | ZBTB16 |
