## Supplemental table S9 for "A chromosome-level, haplotype-resolved genome assembly and annotation for the Eurasian minnow (Leuciscidae: *Phoxinus phoxinus*) provide evidence of haplotype diversity"

**Table S9**: Gene ontology of genes within Insertions/Deletions in both haplomes

| dotCount | description | legend | members | representative |
|---|---|---|---|---|
| 1 | cerebellum vasculature | 1. cerebellum vasculature development | ['GO:0061300', 'GO:0001944', | GO:0061300 |
| 2 | netrin-activated signaling pathway | 2. netrin-activated signaling pathway | ['GO:0038007', 'GO:0007166', | GO:0038007 |
| 3 | dephosphorylation | 3. dephosphorylation | ['GO:0016311'] | GO:0016311 |
| 4 | protein processing | 4. protein processing | ['GO:0051604', 'GO:0016485', | GO:0016485 |
| 5 | regulation of calcium ion- | 5. regulation of calcium ion-dependent | ['GO:0017158', 'GO:1903305', | GO:0017158 |
| 6 | peptidyl-serine phosphorylation | 6. peptidyl-serine phosphorylation | ['GO:0018105', 'GO:0018209', | GO:0018105 |
| 7 | transmission of nerve impulse | 7. transmission of nerve impulse | ['GO:0019226', 'GO:0050877', | GO:0019226 |
| 8 | calcium-ion regulated exocytosis | 8. calcium-ion regulated exocytosis | ['GO:0045055', 'GO:0017156', | GO:0017156 |
| 9 | lipid localization | 9. lipid localization | ['GO:0010876'] | GO:0010876 |
| 10 | neural crest cell differentiation | 10. neural crest cell differentiation | ['GO:0048762', 'GO:0014033', | GO:0014033 |
| 11 | phospholipid transport | 11. phospholipid transport | ['GO:0006869', 'GO:0015914', | GO:0015914 |
| 12 | cell morphogenesis involved in | 12. cell morphogenesis involved in neuron | ['GO:0048667', 'GO:0000902', | GO:0048667 |
| 13 | axon extension involved in axon | 13. axon extension involved in axon | ['GO:0048846', 'GO:1902284', | GO:0048846 |
| 14 | developmental pigmentation | 14. developmental pigmentation | ['GO:0048066', 'GO:0043473'] | GO:0048066 |
| 15 | regulation of developmental | 15. regulation of developmental growth | ['GO:0048638', 'GO:0040008', | GO:0048638 |
| 16 | negative chemotaxis | 16. negative chemotaxis | ['GO:0050919', 'GO:0006935', | GO:0050919 |
| 17 | late stripe melanocyte | 17. late stripe melanocyte differentiation | ['GO:0050934', 'GO:0030318', | GO:0050934 |
| 18 | D-xylose metabolic process | 18. D-xylose metabolic process | ['GO:0042732', 'GO:0019321', | GO:0042732 |
| 19 | engulfment of apoptotic cell | 19. engulfment of apoptotic cell | ['GO:0043652', 'GO:0006911', | GO:0043652 |
| 20 | post-translational protein | 20. post-translational protein modification | ['GO:0043687', 'GO:0006468', | GO:0043687 |
| 21 | positive regulation of filopodium | 21. positive regulation of filopodium | ['GO:0051491', 'GO:0051489', | GO:0051491 |
| 22 | neurotransmitter receptor | 22. neurotransmitter receptor | ['GO:0099590', 'GO:0031623', | GO:0099590 |
| 23 | cell-cell signaling by wnt | 23. cell-cell signaling by wnt | ['GO:0198738', 'GO:0007267', | GO:0198738 |
| 24 | extrinsic apoptotic signaling | 24. extrinsic apoptotic signaling pathway in | ['GO:0038034', 'GO:0097192', | GO:0097192 |
| 25 | cell-cell adhesion via plasma- | 25. cell-cell adhesion via plasma- | ['GO:0098609', 'GO:0098742', | GO:0098742 |
| 26 | spontaneous synaptic transmission | 26. spontaneous synaptic transmission | ['GO:0098814', 'GO:0098916'] | GO:0098814 |
| 27 | neurotransmitter receptor | 27. neurotransmitter receptor transport, | ['GO:0099637', 'GO:0098943', | GO:0098943 |
| 28 | import into cell | 28. import into cell | ['GO:0098657'] | GO:0098657 |
| 29 | calcium-dependent self proteolysis | 29. calcium-dependent self proteolysis | ['GO:0097264', 'GO:1990092'] | GO:1990092 |
| 30 | mitochondrion-endoplasmic | 30. mitochondrion-endoplasmic reticulum | ['GO:1990456', 'GO:0140056', | GO:1990456 |

| | | | |
|---|---|---|---|
| 31 quinone metabolic process | 31. quinone metabolic process | ['GO:0042180', 'GO:1901661'] | GO:1901661 |
| 32 negative regulation of endoplasmic | 32. negative regulation of endoplasmic | ['GO:1903573', 'GO:1902236', | GO:1902236 |
| 33 chloride transmembrane transport | 33. chloride transmembrane transport | ['GO:0098661', 'GO:1902476', | GO:1902476 |
| 34 cilium-dependent cell motility | 34. cilium-dependent cell motility | ['GO:0060285', 'GO:0001539', | GO:0060285 |
| 35 cilium movement involved in cell | 35. cilium movement involved in cell | ['GO:0060294', 'GO:0003341', | GO:0060294 |
| 36 heart trabecula formation | 36. heart trabecula formation | ['GO:0060343', 'GO:0060347', | GO:0060347 |
| 37 skeletal muscle organ | 37. skeletal muscle organ development | ['GO:0060538', 'GO:0007517', | GO:0060538 |
| 38 left/right pattern formation | 38. left/right pattern formation | ['GO:0060972', 'GO:0003002', | GO:0060972 |
| 39 determination of heart left/right | 39. determination of heart left/right | ['GO:0061371', 'GO:0007368', | GO:0061371 |
| 40 regulation of postsynaptic | 40. regulation of postsynaptic membrane | ['GO:0042391', 'GO:0060078', | GO:0060078 |
| 41 regulation of cell cycle | 41. regulation of cell cycle | ['GO:0051726', 'GO:0050794', | GO:0051726 |
| 42 heart trabecula morphogenesis | 42. heart trabecula morphogenesis | ['GO:0061383', 'GO:0061384', | GO:0061384 |
| 43 positive regulation of synaptic | 43. positive regulation of synaptic | ['GO:0051966', 'GO:0051968', | GO:0051968 |
| 44 activation of GTPase activity | 44. activation of GTPase activity | ['GO:0090630', 'GO:0043547', | GO:0090630 |
| 45 cellular response to BMP stimulus | 45. cellular response to BMP stimulus | ['GO:0071772', 'GO:0071773', | GO:0071773 |
| 46 negative regulation of canonical | 46. negative regulation of canonical Wnt | ['GO:0060828', 'GO:0090090', | GO:0090090 |
| 47 epithelium migration | 47. epithelium migration | ['GO:0090130', 'GO:0090132', | GO:0090132 |
| 48 nucleic acid metabolic process | 48. nucleic acid metabolic process | ['GO:0043170', 'GO:0090304', | GO:0090304 |
| 49 spontaneous neurotransmitter | 49. spontaneous neurotransmitter | ['GO:0061669', 'GO:0007269', | GO:0061669 |
| 50 calcium ion transmembrane | 50. calcium ion transmembrane transport | ['GO:0070588', 'GO:0006816', | GO:0070588 |
| 51 cellular response to calcium ion | 51. cellular response to calcium ion | ['GO:1902075', 'GO:0071277', | GO:0071277 |
| 52 complement activation | 52. complement activation | ['GO:0006956', 'GO:0006959', | GO:0006956 |
| 53 cell cycle | 53. cell cycle | ['GO:0007049'] | GO:0007049 |
| 54 establishment or maintenance of | 54. establishment or maintenance of cell | ['GO:0007163'] | GO:0007163 |
| 55 apoptotic process | 55. apoptotic process | ['GO:0006915', 'GO:0012501', | GO:0006915 |
| 56 cell population proliferation | 56. cell population proliferation | ['GO:0008283'] | GO:0008283 |
| 57 positive regulation of cell | 57. positive regulation of cell population | ['GO:0042127', 'GO:0008284', | GO:0008284 |
| 58 insulin receptor signaling pathway | 58. insulin receptor signaling pathway | ['GO:0008286', 'GO:0030509', | GO:0008286 |
| 59 glycoprotein biosynthetic process | 59. glycoprotein biosynthetic process | ['GO:0009100', 'GO:0009101'] | GO:0009101 |
| 60 vitamin biosynthetic process | 60. vitamin biosynthetic process | ['GO:0006766', 'GO:0009110', | GO:0009110 |
| 61 Rho protein signal transduction | 61. Rho protein signal transduction | ['GO:0007265', 'GO:0007266', | GO:0007266 |
| 62 neuromuscular junction | 62. neuromuscular junction development | ['GO:0007528', 'GO:0050808', | GO:0007528 |
| 63 ultradian rhythm | 63. ultradian rhythm | ['GO:0048511', 'GO:0007624'] | GO:0007624 |

| | | | |
|---|---|---|---|
| 64 motor neuron axon guidance | 64. motor neuron axon guidance | ['GO:0008045', 'GO:0060271', | GO:0008045 |
| 65 sphingomyelin metabolic process | 65. sphingomyelin metabolic process | ['GO:0006684', 'GO:0006665', | GO:0006684 |
| 66 chondrocyte development | 66. chondrocyte development | ['GO:0002063', 'GO:0030097', | GO:0002063 |
| 67 tRNA wobble uridine modification | 67. tRNA wobble uridine modification | ['GO:0002097', 'GO:0002098', | GO:0002098 |
| 68 cardiac chamber morphogenesis | 68. cardiac chamber morphogenesis | ['GO:0003206', 'GO:0002009', | GO:0003206 |
| 69 fructose 6-phosphate metabolic | 69. fructose 6-phosphate metabolic | ['GO:0006002'] | GO:0006002 |
| 70 negative regulation of | 70. negative regulation of transcription by | ['GO:0000122', 'GO:0045892', | GO:0000122 |
| 71 MAPK cascade | 71. MAPK cascade | ['GO:0000165', 'GO:0007186', | GO:0000165 |
| 72 protein polyubiquitination | 72. protein polyubiquitination | ['GO:0000209', 'GO:0016567', | GO:0000209 |
| 73 mRNA splicing, via spliceosome | 73. mRNA splicing, via spliceosome | ['GO:0000377', 'GO:0000398', | GO:0000398 |
| 74 UDP-N-acetylglucosamine | 74. UDP-N-acetylglucosamine biosynthetic | ['GO:0046349', 'GO:0006048', | GO:0006048 |
| 75 translational elongation | 75. translational elongation | ['GO:0006414', 'GO:0006412', | GO:0006414 |
| 76 regulation of translation | 76. regulation of translation | ['GO:0051246', 'GO:0006417', | GO:0006417 |
| 77 leucyl-tRNA aminoacylation | 77. leucyl-tRNA aminoacylation | ['GO:0006429', 'GO:0006418', | GO:0006429 |
| 78 protein N-linked glycosylation | 78. protein N-linked glycosylation | ['GO:0006486', 'GO:0006487', | GO:0006487 |
| 79 glutamine metabolic process | 79. glutamine metabolic process | ['GO:0009064', 'GO:0006541', | GO:0006541 |
| 80 tyrosine catabolic process | 80. tyrosine catabolic process | ['GO:0006570', 'GO:0006572', | GO:0006572 |
| 81 phosphatidylethanolamine | 81. phosphatidylethanolamine biosynthetic | ['GO:0046337', 'GO:0006646', | GO:0006646 |
| 82 mRNA export from nucleus | 82. mRNA export from nucleus | ['GO:0006406', 'GO:0006405', | GO:0006406 |
| 83 RNA localization | 83. RNA localization | ['GO:0006403', 'GO:0033036'] | GO:0006403 |
| 84 dGTP catabolic process | 84. dGTP catabolic process | ['GO:0046070', 'GO:0006203', | GO:0006203 |
| 85 'de novo' pyrimidine nucleobase | 85. 'de novo' pyrimidine nucleobase | ['GO:0006207', 'GO:0019856', | GO:0006207 |
| 86 DNA-templated DNA replication | 86. DNA-templated DNA replication | ['GO:0006260', 'GO:0006261', | GO:0006261 |
| 87 regulation of DNA-templated | 87. regulation of DNA-templated | ['GO:2001141', 'GO:0006355', | GO:0006355 |
| 88 transcription by RNA polymerase I | 88. transcription by RNA polymerase I | ['GO:0006360', 'GO:0006351', | GO:0006360 |
| 89 transcription by RNA polymerase II | 89. transcription by RNA polymerase II | ['GO:0006366'] | GO:0006366 |
| 90 transcription elongation by RNA | 90. transcription elongation by RNA | ['GO:0006354', 'GO:0006368'] | GO:0006368 |
| 91 chromatin remodeling | 91. chromatin remodeling | ['GO:0006325', 'GO:0006338', | GO:0006338 |
| 92 neutrophil chemotaxis | 92. neutrophil chemotaxis | ['GO:1990266', 'GO:0030593', | GO:0030593 |
| 93 cortical cytoskeleton organization | 93. cortical cytoskeleton organization | ['GO:0030865', 'GO:0030036', | GO:0030865 |
| 94 endoplasmic reticulum unfolded | 94. endoplasmic reticulum unfolded | ['GO:0030968', 'GO:0030433', | GO:0030968 |
| 95 replication fork processing | 95. replication fork processing | ['GO:0045005', 'GO:0031297'] | GO:0031297 |
| 96 actin filament-based process | 96. actin filament-based process | ['GO:0030029'] | GO:0030029 |

| | | | |
|---|---|---|---|
| 97 regulation of actin cytoskeleton | 97. regulation of actin cytoskeleton | ['GO:0051493', 'GO:0032956', | GO:0032956 |
| 98 embryonic heart tube | 98. embryonic heart tube development | ['GO:0035050', 'GO:0048565', | GO:0035050 |
| 99 TOR signaling | 99. TOR signaling | ['GO:0031929'] | GO:0031929 |
| 100 regulation of appetite | 100. regulation of appetite | ['GO:0032098', 'GO:0008360', | GO:0032098 |
| 101 endoplasmic reticulum calcium ion | 101. endoplasmic reticulum calcium ion | ['GO:0032469', 'GO:0051560', | GO:0032469 |
| 102 positive regulation of type I | 102. positive regulation of type I interferon | ['GO:0001819', 'GO:0032481', | GO:0032481 |
| 103 type I interferon production | 103. type I interferon production | ['GO:0032606', 'GO:0001816', | GO:0032606 |
| 104 cellular process | 104. cellular process | ['GO:0009987'] | GO:0009987 |
| 105 rRNA transcription | 105. rRNA transcription | ['GO:0098781', 'GO:0009303', | GO:0009303 |
| 106 macroautophagy | 106. macroautophagy | ['GO:0006914', 'GO:0016236', | GO:0016236 |
| 107 peptide biosynthetic process | 107. peptide biosynthetic process | ['GO:0043043', 'GO:0006518', | GO:0043043 |
| 108 positive regulation of apoptotic | 108. positive regulation of apoptotic | ['GO:0043065', 'GO:0043068', | GO:0043065 |
| 109 apoptotic cell clearance | 109. apoptotic cell clearance | ['GO:0043277', 'GO:0006909', | GO:0043277 |
| 110 regulation of dephosphorylation | 110. regulation of dephosphorylation | ['GO:0035303', 'GO:0019220', | GO:0035303 |
| 111 lymph vessel morphogenesis | 111. lymph vessel morphogenesis | ['GO:0036303', 'GO:0009653', | GO:0036303 |
| 112 calcitriol biosynthetic process from | 112. calcitriol biosynthetic process from | ['GO:0042368', 'GO:0036378', | GO:0036378 |
| 113 negative regulation of locomotion | 113. negative regulation of locomotion | ['GO:0040013', 'GO:2000145', | GO:0040013 |
| 114 regulation of RNA splicing | 114. regulation of RNA splicing | ['GO:0043484'] | GO:0043484 |
| 115 lipid catabolic process | 115. lipid catabolic process | ['GO:0016042', 'GO:1901575', | GO:0016042 |
| 116 autophagosome assembly | 116. autophagosome assembly | ['GO:0000045', 'GO:1905037', | GO:0000045 |
