## Supplemental table S10 for "A chromosome-level, haplotype-resolved genome assembly and annotation for the Eurasian minnow (Leuciscidae: *Phoxinus phoxinus*) provide evidence of haplotype diversity"

***Table S10:*** *Genes present in Inversions between both Halpomes*

| Gene Name | Gene ID |
|---|---|
| Family 104 | FAM104A |
| Family 104 | FAM104A |
| HID1 domain containing | HID1 |
| HID1 domain containing | HID1 |
| Otopetrin 2 | OTOP2 |
| Usher syndrome 1G (autosomal recessive) | USH1G |
| Amnion associated transmembrane protein | AMN |
| Golgi apparatus membrane protein TVP23 homolog | TVP23B |
| ligand-gated ion | ZACN |
| ligand-gated ion | ZACN |
| endoplasmic reticulum-plasma membrane tethering | ESYT3 |
| Domain of unknown function (DUF4371) | - |
| extracellular ligand-gated ion channel activity | - |
| Serine threonine kinase 19 | STK19 |
| Serine threonine kinase 19 | STK19 |
| Serine threonine kinase 19 | STK19 |
| Serine threonine kinase 19 | STK19 |
| Ras-related C3 botulinum toxin substrate 1 (rho family, small GTP | RAC1 |
| Ras-related C3 botulinum toxin substrate 1 (rho family, small GTP | RAC1 |
| Domain of unknown function (DUF4560) | SMIM10 |
| Protein kinase C, alpha | PRKCA |
| Protein kinase C, alpha | PRKCA |
| Centrosomal protein | CEP112 |
| Centrosomal protein | CEP112 |
| Axin 2 (conductin, axil) | AXIN2 |
| Axin 2 (conductin, axil) | AXIN2 |
| Testis expressed 2 | TEX2 |
| Testis expressed 2 | TEX2 |
| Endoplasmic reticulum to nucleus signaling 1 | ERN1 |
| Sterile alpha motif domain-containing protein 9-like | SAMD9L |

| | |
|---|---|
| Sterile alpha motif domain-containing protein 9-like | SAMD9L |
| Chromosome 9 open reading frame 69 | C9orf69 |
| Chromosome 9 open reading frame 69 | C9orf69 |
| NACC family member 2, BEN and BTB (POZ) domain containing | NACC2 |
| NACC family member 2, BEN and BTB (POZ) domain containing | NACC2 |
| Nucleoporin 214 | NUP214 |
| Family with sequence similarity 78 member | FAM78A |
| Family with sequence similarity 78 member | FAM78A |
| Phosphatidic acid phosphatase type 2 domain containing 3 | PPAPDC3 |
| Phosphatidic acid phosphatase type 2 domain containing 3 | PPAPDC3 |
| - | - |
| Proline-rich coiled-coil 2B | PRRC2B |
| Proline-rich coiled-coil 2B | PRRC2B |
| Endothelial differentiation-related factor 1 | EDF1 |
| Endothelial differentiation-related factor 1 | EDF1 |
| Endothelial differentiation-related factor 1 | EDF1 |
| Belongs to the vasopressin oxytocin family | OXT |
| Belongs to the vasopressin oxytocin family | OXT |
| fatty acid binding protein | FABP1 |
| fatty acid binding protein | FABP1 |
| Sepiapterin reductase | SPR |
| Sepiapterin reductase | SPR |
| nuclear retention of pre-mRNA with aberrant 3'-ends at the site of | EXOSC2 |
| nuclear retention of pre-mRNA with aberrant 3'-ends at the site of | EXOSC2 |
| C-abl oncogene 1, receptor tyrosine kinase | ABL1 |
| C-abl oncogene 1, receptor tyrosine kinase | ABL1 |
| C-abl oncogene 1, receptor tyrosine kinase | ABL1 |
| C-abl oncogene 1, receptor tyrosine kinase | ABL1 |
| apical protein localization | RAB14 |
| apical protein localization | RAB14 |
| Complement component C1q domain. | - |
| Complement component C1q domain. | - |
| dedicator of cytokinesis | DOCK10 |

| | |
|---|---|
| dedicator of cytokinesis | DOCK10 |
| Short stature homeobox | SHOX2 |
| Short stature homeobox | SHOX2 |
| Short stature homeobox | SHOX2 |
| Arginine serine-rich coiled-coil 1 | RSRC1 |
| Si dkey-121j17.6 | - |
| Belongs to the multicopper oxidase family | HEPHL1 |
| Pannexin 1b | PANX1 |
| Pannexin 1b | PANX1 |
| hAT family C-terminal dimerisation region | - |
| - | - |
| - | - |
| DDE superfamily endonuclease | - |
| DDE superfamily endonuclease | - |
| Zinc finger protein | - |
| RAB3A interacting protein (rabin3) | RAB3IP |
| RAB3A interacting protein (rabin3) | RAB3IP |
| RAB3A interacting protein (rabin3) | RAB3IP |
| RAB3A interacting protein (rabin3) | RAB3IP |
| Myelin regulatory factor-like | MYRFL |
| protein dimerization activity | - |
| DDE superfamily endonuclease | - |
| - | - |
| MADF | - |
| - | - |
| pyridoxal phosphate binding | ACCS |
| chromosome 14 open reading frame 93 | C14orf93 |
| polyketide synthase | FASN |
| L1 transposable element RBD-like domain | L1TD1 |
| L1 transposable element RBD-like domain | L1TD1 |
| Sterol-sensing domain of SREBP cleavage-activation | PTCHD3 |
| Sterol-sensing domain of SREBP cleavage-activation | PTCHD3 |
| Protein kinase, membrane associated tyrosine threonine 1 | PKMYT1 |

| | |
|---|---|
| SUMO1 sentrin SMT3 specific peptidase 2 | SENP2 |
