## Supplemental table S11 for "A chromosome-level, haplotype-resolved genome assembly and annotation for the Eurasian minnow (Leuciscidae: *Phoxinus phoxinus*) provide evidence of haplotype diversity"

**Table S11**: Gene ontology of genes within Inversions in both haplomes

| dotCo | description | legend | members | representative |
|---|---|---|---|---|
| 1 | phagocytosis | 1. phagocytosis | ['GO:0006909', | GO:0006909 |
| 2 | phagosome maturation involved in | 2. phagosome maturation involved | ['GO:0090386', | GO:0090386 |
| 3 | phagolysosome assembly involved in | 3. phagolysosome assembly involved | ['GO:0090387', | GO:0090387 |
| 4 | cortical cytoskeleton organization | 4. cortical cytoskeleton organization | ['GO:0030865', | GO:0030865 |
| 5 | RNA-mediated gene silencing | 5. RNA-mediated gene silencing | ['GO:0031047', | GO:0031047 |
| 6 | endocytic recycling | 6. endocytic recycling | ['GO:0032456', | GO:0032456 |
| 7 | regulation of cell migration | 7. regulation of cell migration | ['GO:0030334', | GO:0030334 |
| 8 | regulation of actin cytoskeleton | 8. regulation of actin cytoskeleton | ['GO:0032970', | GO:0032956 |
| 9 | protein desumoylation | 9. protein desumoylation | ['GO:0016926', | GO:0016926 |
| 10 | Rac protein signal transduction | 10. Rac protein signal transduction | ['GO:0016601', | GO:0016601 |
| 11 | protein autoprocessing | 11. protein autoprocessing | ['GO:0016540', | GO:0016540 |
| 12 | spinal cord development | 12. spinal cord development | ['GO:0021510', | GO:0021510 |
| 13 | post-translational protein modification | 13. post-translational protein | ['GO:0043687', | GO:0043687 |
| 14 | regulation of apoptotic process | 14. regulation of apoptotic process | ['GO:0042981', | GO:0042981 |
| 15 | defense response to bacterium | 15. defense response to bacterium | ['GO:0042742', | GO:0042742 |
| 16 | ribosome biogenesis | 16. ribosome biogenesis | ['GO:0042254', | GO:0042254 |
| 17 | cell migration involved in gastrulation | 17. cell migration involved in | ['GO:0042074', | GO:0042074 |
| 18 | engulfment of apoptotic cell | 18. engulfment of apoptotic cell | ['GO:0043652', | GO:0043652 |
| 19 | peptide biosynthetic process | 19. peptide biosynthetic process | ['GO:0043043', | GO:0043043 |
| 20 | modification-dependent | 20. modification-dependent | ['GO:0043632', | GO:0043632 |
| 21 | positive regulation of GTPase activity | 21. positive regulation of GTPase | ['GO:0043547', | GO:0043547 |
| 22 | dendrite development | 22. dendrite development | ['GO:0016358', | GO:0016358 |
| 23 | Golgi to endosome transport | 23. Golgi to endosome transport | ['GO:0016482', | GO:0006895 |
| 24 | lipid transport | 24. lipid transport | ['GO:0006869', | GO:0006869 |
| 25 | apoptotic process | 25. apoptotic process | ['GO:0006915', | GO:0006915 |
| 26 | fatty acid biosynthetic process | 26. fatty acid biosynthetic process | ['GO:0006633', | GO:0006633 |
| 27 | establishment or maintenance of cell | 27. establishment or maintenance of | ['GO:0007163'] | GO:0007163 |
| 28 | endoplasmic reticulum organization | 28. endoplasmic reticulum | ['GO:0007029', | GO:0007029 |
| 29 | actin filament organization | 29. actin filament organization | ['GO:0007015', | GO:0007015 |
| 30 | alternative mRNA splicing, via | 30. alternative mRNA splicing, via | ['GO:0000380', | GO:0000380 |

| | | | |
|---|---|---|---|
| 31 maturation of 5.8S rRNA from | 31. maturation of 5.8S rRNA from | ['GO:0000460', | GO:0000466 |
| 32 neural retina development | 32. neural retina development | ['GO:0003407', | GO:0003407 |
| 33 gene expression | 33. gene expression | ['GO:0010467'] | GO:0010467 |
| 34 endomembrane system organization | 34. endomembrane system | ['GO:0010256'] | GO:0010256 |
| 35 lipid localization | 35. lipid localization | ['GO:0010876'] | GO:0010876 |
| 36 myotube differentiation | 36. myotube differentiation | ['GO:0014902', | GO:0014902 |
| 37 regulation of cell shape | 37. regulation of cell shape | ['GO:0022604', | GO:0008360 |
| 38 cell population proliferation | 38. cell population proliferation | ['GO:0008283'] | GO:0008283 |
| 39 myoblast fusion | 39. myoblast fusion | ['GO:0007520', | GO:0007520 |
| 40 ribonucleotide metabolic process | 40. ribonucleotide metabolic process | ['GO:0019693', | GO:0009259 |
| 41 cell morphogenesis involved in neuron | 41. cell morphogenesis involved in | ['GO:0048667', | GO:0048667 |
| 42 canonical Wnt signaling pathway | 42. canonical Wnt signaling pathway | ['GO:0016055', | GO:0060070 |
| 43 retina development in camera-type | 43. retina development in camera- | ['GO:0060041', | GO:0060041 |
| 44 dendritic spine development | 44. dendritic spine development | ['GO:0060996', | GO:0060996 |
| 45 response to (R)-carnitine | 45. response to (R)-carnitine | ['GO:0061959', | GO:0061959 |
| 46 endoplasmic reticulum-plasma | 46. endoplasmic reticulum-plasma | ['GO:0051643', | GO:0061817 |
| 47 cellular response to chemical stimulus | 47. cellular response to chemical | ['GO:0070887', | GO:0070887 |
| 48 synapse organization | 48. synapse organization | ['GO:0034330', | GO:0050808 |
| 49 positive regulation of RNA metabolic | 49. positive regulation of RNA | ['GO:0045935', | GO:0051254 |
| 50 equilibrioception | 50. equilibrioception | ['GO:0050957', | GO:0050957 |
| 51 regulatory ncRNA processing | 51. regulatory ncRNA processing | ['GO:0070918'] | GO:0070918 |
| 52 cell-cell signaling by wnt | 52. cell-cell signaling by wnt | ['GO:0198738', | GO:0198738 |
| 53 visual system development | 53. visual system development | ['GO:0048880', | GO:0150063 |
| 54 proton transmembrane transport | 54. proton transmembrane transport | ['GO:1902600', | GO:1902600 |
| 55 polyadenylation-dependent snoRNA | 55. polyadenylation-dependent | ['GO:0016074', | GO:0071051 |
| 56 CUT catabolic process | 56. CUT catabolic process | ['GO:0071043', | GO:0071034 |
| 57 import into cell | 57. import into cell | ['GO:0098657'] | GO:0098657 |
| 58 negative regulation of canonical Wnt | 58. negative regulation of canonical | ['GO:0060828', | GO:0090090 |
