## Supplemental table S12 for "A chromosome-level, haplotype-resolved genome assembly and annotation for the Eurasian minnow (Leuciscidae: *Phoxinus phoxinus*) provide evidence of haplotype diversity"

**Table S12:** Summary statistics of orthology between proteomes of selected Teleost species

| | Channel Fish | Chinese Sucker | Common Carp | Eurasian Minnow |
|---|---|---|---|---|
| Number of genes | 24165 | 39515 | 49592 | 23397 |
| Number of genes in orthogroups | 23379 | 38634 | 45414 | 22651 |
| Number of unassigned genes | 786 | 881 | 4178 | 746 |
| Percentage of genes in orthogroups | 96.7 | 97.8 | 91.6 | 96.8 |
| Percentage of unassigned genes | 3.3 | 2.2 | 8.4 | 3.2 |
| Number of orthogroups containing species | 18653 | 20424 | 23387 | 19476 |
| Percentage of orthogroups containing species | 67.4 | 73.8 | 84.5 | 70.4 |
| Number of species-specific orthogroups | 198 | 102 | 958 | 72 |
| Number of genes in species-specific orthogroups | 1500 | 455 | 2248 | 273 |
| Percentage of genes in species-specific orthogroups | 6.2 | 1.2 | 4.5 | 1.2 |

| Fathead Minnow | GoldFish | Goldline Barbel | Grass Carp | Horned Barbel | Wuchang Bream |
|---|---|---|---|---|---|
| 26763 | 53614 | 46135 | 25255 | 44583 | 30620 |
| 26455 | 51621 | 43108 | 25057 | 42496 | 30277 |
| 308 | 1993 | 3027 | 198 | 2087 | 343 |
| 98.8 | 96.3 | 93.4 | 99.2 | 95.3 | 98.9 |
| 1.2 | 3.7 | 6.6 | 0.8 | 4.7 | 1.1 |
| 20792 | 21956 | 22611 | 20185 | 22055 | 21207 |
| 75.1 | 79.3 | 81.7 | 72.9 | 79.7 | 76.6 |
| 66 | 293 | 233 | 14 | 111 | 77 |
| 254 | 805 | 489 | 55 | 236 | 358 |
| 0.9 | 1.5 | 1.1 | 0.2 | 0.5 | 1.2 |

| Zebrafish |
| --- |
| 32816 |
| 32171 |
| 645 |
| 98 |
| 2 |
| 20690 |
| 74.8 |
| 195 |
| 1302 |
| 4 |
