## Supplemental table S13 for "A chromosome-level, haplotype-resolved genome assembly and annotation for the Eurasian minnow (Leuciscidae: *Phoxinus phoxinus*) provide evidence of haplotype diversity"

**Table S13:** *Phoxinus phoxinus* genome specific genes

| Uniprot ID | Protein | Gene Ontology | |
|---|---|---|---|
| A0A8C8RDR1 | HAT C-terminal dimerisation domain-containing protein | protein dimerization activity [GO:0046983] | GO:0046983 |
| A0A8C8RDR1_9SAUR | HAT C-terminal dimerisation domain-containing protein | protein dimerization activity [GO:0046983] | GO:0046983 |
| A0A8C1EUN7 | HAT C-terminal dimerisation domain-containing protein | protein dimerization activity [GO:0046983] | GO:0046983 |
| A0A8C1EUN7_CYPCA | HAT C-terminal dimerisation domain-containing protein | protein dimerization activity [GO:0046983] | GO:0046983 |
| A0A672GQY9 | SPIN-DOC-like zinc-finger domain-containing protein | protein dimerization activity [GO:0046983] | GO:0046983 |
| A0A672GQY9_SALFA | SPIN-DOC-like zinc-finger domain-containing protein | protein dimerization activity [GO:0046983] | GO:0046983 |
| A0A667Y992 | TTF-type domain-containing protein | protein dimerization activity [GO:0046983] | GO:0046983 |
| A0A667Y992_9TELE | TTF-type domain-containing protein | protein dimerization activity [GO:0046983] | GO:0046983 |
| A0A8C5LX81 | Zinc finger MYM-type protein 1 | protein dimerization activity [GO:0046983] | GO:0046983 |
| A0A8C5LX81_9ANUR | Zinc finger MYM-type protein 1 | protein dimerization activity [GO:0046983] | GO:0046983 |
| A0A8D3C299 | DDE Tnp4 domain-containing protein | metal ion binding [GO:0046872] | GO:0046872 |
| A0A8D3C299_SCOMX | DDE Tnp4 domain-containing protein | metal ion binding [GO:0046872] | GO:0046872 |
| A0A498NBI9 | Nuclease HARBI1 | metal ion binding [GO:0046872] | GO:0046872 |
| A0A498NBI9_LABRO | Nuclease HARBI1 | metal ion binding [GO:0046872] | GO:0046872 |
| A0A6P6R529 | Protein ANTAGONIST OF LIKE HETEROCHROMATIN PROTEIN 1-like | metal ion binding [GO:0046872] | GO:0046872 |
| A0A6P6R529_CARAU | Protein ANTAGONIST OF LIKE HETEROCHROMATIN PROTEIN 1-like | metal ion binding [GO:0046872] | GO:0046872 |
| A0A1A8N771 | Si:dkey-228l14.1 | metal ion binding [GO:0046872] | GO:0046872 |
| A0A1A8N771_9TELE | Si:dkey-228l14.1 | metal ion binding [GO:0046872] | GO:0046872 |
| A0A6P8V9C8 | Uncharacterized protein LOC117553609 | metal ion binding [GO:0046872] | GO:0046872 |
| A0A6P8V9C8_GYMAC | Uncharacterized protein LOC117553609 | metal ion binding [GO:0046872] | GO:0046872 |

| | | | |
|---|---|---|---|
| A0A6P8VFR7 | Uncharacterized protein LOC117554636 | metal ion binding [GO:0046872] | GO:0046872 |
| A0A6P8VFR7_GYMAC | Uncharacterized protein LOC117554636 | metal ion binding [GO:0046872] | GO:0046872 |
| A0A3B3T7F3 | Zgc:194221 | metal ion binding [GO:0046872] | GO:0046872 |
| A0A3B3T7F3_9TELE | Zgc:194221 | metal ion binding [GO:0046872] | GO:0046872 |
| A0A498M129 | E3 SUMO-protein ligase KIAA1586-like | ligase activity [GO:0016874]; protein dimerization activity [GO:0046983] | GO:0016874; GO:0046983 |
| A0A498M129_LABRO | E3 SUMO-protein ligase KIAA1586-like | ligase activity [GO:0016874]; protein dimerization activity [GO:0046983] | GO:0016874; GO:0046983 |
| A0A7J6C8Y6 | Uncharacterized protein | membrane [GO:0016020] | GO:0016020 |
| A0A7J6C8Y6_9TELE | Uncharacterized protein | membrane [GO:0016020] | GO:0016020 |
| A0A7J6C5D6 | Uncharacterized protein | membrane [GO:0016020] | GO:0016020 |
| A0A7J6C5D6_9TELE | Uncharacterized protein | membrane [GO:0016020] | GO:0016020 |
| A0A3N0XSF9 | Nuclear factor 7, ovary | zinc ion binding [GO:0008270] | GO:0008270 |
| A0A3N0XSF9_ANAGA | Nuclear factor 7, ovary | zinc ion binding [GO:0008270] | GO:0008270 |
| A0A6P7X1B1 | Uncharacterized protein LOC115461455 | metal ion binding [GO:0046872]; protein dimerization activity [GO:0046983]; regulation of DNA-templated transcription [GO:0006355] | GO:0006355; GO:0046872; GO:0046983 |
| A0A6P7X1B1_9AMPH | Uncharacterized protein LOC115461455 | metal ion binding [GO:0046872]; protein dimerization activity [GO:0046983]; regulation of DNA-templated transcription [GO:0006355] | GO:0006355; GO:0046872; GO:0046983 |
| A0A1S3HT46 | Uncharacterized protein LOC106157950 | zinc ion binding [GO:0008270]; DNA repair [GO:0006281] | GO:0006281; GO:0008270 |
| A0A1S3HT46_LINUN | Uncharacterized protein LOC106157950 | zinc ion binding [GO:0008270]; DNA repair [GO:0006281] | GO:0006281; GO:0008270 |
| A0A6P8VHT6 | Uncharacterized protein LOC117558146 | zinc ion binding [GO:0008270]; DNA repair [GO:0006281] | GO:0006281; GO:0008270 |
| A0A6P8VHT6_GYMAC | Uncharacterized protein LOC117558146 | zinc ion binding [GO:0008270]; DNA repair [GO:0006281] | GO:0006281; GO:0008270 |
| A0A669DFR9 | Uncharacterized LOC102078392 | DNA repair [GO:0006281] | GO:0006281 |
| A0A669DFR9_ORENI | Uncharacterized LOC102078392 | DNA repair [GO:0006281] | GO:0006281 |

| | | | |
|---|---|---|---|
| A0A3N0XYI6 | PAK4-inhibitor inka2 | nucleus [GO:0005634]; protein serine/threonine kinase inhibitor activity | GO:0005634; GO:0030291 |
| A0A3N0XYI6_ANAGA | PAK4-inhibitor inka2 | nucleus [GO:0005634]; protein serine/threonine kinase inhibitor activity | GO:0005634; GO:0030291 |
| A0A8C1T586 | Sestrin 1 | cytoplasm [GO:0005737]; nucleus [GO:0005634]; cellular response to stimulus [GO:0051716]; negative regulation of TORC1 signaling [GO:1904262]; regulation of response to reactive oxygen species [GO:1901031] | GO:0005634; GO:0005737; GO:0051716; GO:1901031; GO:1904262 |
| A0A8C1T586_CYPCA | Sestrin 1 | cytoplasm [GO:0005737]; nucleus [GO:0005634]; cellular response to stimulus [GO:0051716]; negative regulation of TORC1 signaling [GO:1904262]; regulation of response to reactive oxygen species [GO:1901031] | GO:0005634; GO:0005737; GO:0051716; GO:1901031; GO:1904262 |
| A0A8M1PAF3 | Glutathione peroxidase | glutathione peroxidase activity [GO:0004602]; response to oxidative stress [GO:0006979] | GO:0004602; GO:0006979 |
| A0A8M1PAF3_DANRE | Glutathione peroxidase | glutathione peroxidase activity [GO:0004602]; response to oxidative stress [GO:0006979] | GO:0004602; GO:0006979 |
| A0A7J6DEY2 | Glutathione peroxidase 4 | glutathione peroxidase activity [GO:0004602]; response to oxidative stress [GO:0006979] | GO:0004602; GO:0006979 |
| A0A7J6DEY2_9TELE | Glutathione peroxidase 4 | glutathione peroxidase activity [GO:0004602]; response to oxidative stress [GO:0006979] | GO:0004602; GO:0006979 |
| A0A6J2UU45 | Deoxynucleoside triphosphate triphosphohydrolase SAMHD1 | chromosome [GO:0005694]; catalytic activity [GO:0003824]; defense response to virus [GO:0051607]; DNA repair [GO:0006281]; DNA replication [GO:0006260] | GO:0003824; GO:0005694; GO:0006260; GO:0006281; GO:0051607 |

| | | | |
|---|---|---|---|
| A0A6J2UU45_CHACN | Deoxynucleoside triphosphate triphosphohydrolase SAMHD1 | chromosome [GO:0005694]; catalytic activity [GO:0003824]; defense response to virus [GO:0051607]; DNA repair [GO:0006281]; DNA replication [GO:0006260] | GO:0003824; GO:0005694; GO:0006260; GO:0006281; GO:0051607 |
| A0A8T2MFG3 | THAP-type domain-containing protein | DNA binding [GO:0003677]; metal ion binding [GO:0046872] | GO:0003677; GO:0046872 |
| A0A8T2MFG3_ASTMX | THAP-type domain-containing protein | DNA binding [GO:0003677]; metal ion binding [GO:0046872] | GO:0003677; GO:0046872 |
| A0A3B3RYR3 | BESS domain-containing protein | nucleus [GO:0005634]; DNA binding [GO:0003677] | GO:0003677; GO:0005634 |
| A0A3B3RYR3_9TELE | BESS domain-containing protein | nucleus [GO:0005634]; DNA binding [GO:0003677] | GO:0003677; GO:0005634 |
| A0A3N0YPG9 | C2H2-type domain-containing protein | | |
| A0A3N0YPG9_ANAGA | C2H2-type domain-containing protein | | |
| A0A7J6CNT2 | Chromo domain-containing protein | | |
| A0A7J6CNT2_9TELE | Chromo domain-containing protein | | |
| A0A3N0Z136 | Coiled-coil domain-containing protein 173 | | |
| A0A3N0Z136_ANAGA | Coiled-coil domain-containing protein 173 | | |
| A0A8C1LWF1 | Core-binding (CB) domain-containing protein | | |
| A0A8C1LWF1_CYPCA | Core-binding (CB) domain-containing protein | | |
| A0A7J6CPP1 | DUF4806 domain-containing protein | | |
| A0A7J6CPP1_9TELE | DUF4806 domain-containing protein | | |
| A0A8C9X006 | HAT C-terminal dimerisation domain-containing protein | | |
| A0A8C9X006_SANLU | HAT C-terminal dimerisation domain-containing protein | | |
| A0A7J6D8R6 | Ig-like domain-containing protein | | |
| A0A7J6D8R6_9TELE | Ig-like domain-containing protein | | |

| | |
|---|---|
| A0A8C1W0D5 | Ig-like domain-containing protein |
| A0A8C1W0D5_CYPCA | Ig-like domain-containing protein |
| A0A7J6CHV6 | Myb/SANT-like DNA-binding domain-containing protein |
| A0A7J6CHV6_9TELE | Myb/SANT-like DNA-binding domain-containing protein |
| A0A7J6C6I1 | Saposin B-type domain-containing protein |
| A0A7J6C6I1_9TELE | Saposin B-type domain-containing protein |
| A0A7J6C5D8 | Saposin B-type domain-containing protein |
| A0A7J6C5D8_9TELE | Saposin B-type domain-containing protein |
| A0A6P8VNQ5 | Sterile alpha motif domain-containing protein 3-like isoform X1 |
| A0A6P8VNQ5_GYMAC | Sterile alpha motif domain-containing protein 3-like isoform X1 |
| A0A498MDX1 | Tol2 transposase |
| A0A498MDX1_LABRO | Tol2 transposase |
| A0A6F8ZX98 | Transposase Helix-turn-helix domain-containing protein |
| A0A6F8ZX98_9TELE | Transposase Helix-turn-helix domain-containing protein |
| A0A7J6CNG2 | Trichohyalin-plectin-homology domain-containing protein |
| A0A7J6CNG2_9TELE | Trichohyalin-plectin-homology domain-containing protein |
| A0A3N0Y2X1 | Tripartite motif-containing protein 35 |
| A0A3N0Y2X1_ANAGA | Tripartite motif-containing protein 35 |
| A0A7J6CY62 | Ubiquitin-like domain-containing protein |
| A0A7J6CY62_9TELE | Ubiquitin-like domain-containing protein |
| A0A498LUV0 | Uncharacterized protein |
| A0A498LUV0_LABRO | Uncharacterized protein |
| A0A498M0N6 | Uncharacterized protein |

| | |
|---|---|
| A0A498M0N6_LABRO | Uncharacterized protein |
| A0A6A5EX62 | Uncharacterized protein |
| A0A6A5EX62_PERFL | Uncharacterized protein |
| A0A1A8R7C9 | Uncharacterized protein |
| A0A1A8R7C9_9TELE | Uncharacterized protein |
| A0A7J6BQL4 | Uncharacterized protein |
| A0A7J6BQL4_9TELE | Uncharacterized protein |
| A0A7J6D4P1 | Uncharacterized protein |
| A0A7J6D4P1_9TELE | Uncharacterized protein |
| A0A3N0Z8W9 | Uncharacterized protein |
| A0A3N0Z8W9_ANAGA | Uncharacterized protein |
| A0A8C6TDK7 | Uncharacterized protein |
| A0A8C6TDK7_9GOBI | Uncharacterized protein |
| A0A8C1N735 | Uncharacterized protein |
| A0A8C1N735_CYPCA | Uncharacterized protein |
| A0A6I9NCM8 | Uncharacterized protein LOC104947713 isoform X5 |
| A0A6I9NCM8_9TELE | Uncharacterized protein LOC104947713 isoform X5 |
| A0A2I4AKY7 | Uncharacterized protein LOC106511981 |
| A0A2I4AKY7_AUSLI | Uncharacterized protein LOC106511981 |
| A0A1S3SBY2 | Uncharacterized protein LOC106608449 |
| A0A1S3SBY2_SALSA | Uncharacterized protein LOC106608449 |
| A0A6P6K9C3 | Uncharacterized protein LOC113049977 |
| A0A6P6K9C3_CARAU | Uncharacterized protein LOC113049977 |
| A0A6P6QES6 | Uncharacterized protein LOC113061694 (Uncharacterized protein LOC113111498) |
| A0A6P6QES6_CARAU | Uncharacterized protein LOC113061694 (Uncharacterized protein LOC113111498) |
| A0A6P6P3V1 | Uncharacterized protein LOC113094166 isoform X2 |

| | |
|---|---|
| A0A6P6P3V1_CARAU | Uncharacterized protein LOC113094166 isoform X2 |
| A0A6P6RCR4 | Uncharacterized protein LOC113118533 |
| A0A6P6RCR4_CARAU | Uncharacterized protein LOC113118533 |
| A0A6P8GTP5 | Uncharacterized protein LOC116224715 |
| A0A6P8GTP5_CLUHA | Uncharacterized protein LOC116224715 |
| A0A6P8T7R6 | Uncharacterized protein LOC117538314 isoform X1 |
| A0A6P8T7R6_GYMAC | Uncharacterized protein LOC117538314 isoform X1 |
| A0A5A9PJY1 | Zinc finger BED domain-containing protein 4 |
| A0A5A9PJY1_9TELE | Zinc finger BED domain-containing protein 4 |
| A0A3N0XQY8 | Zinc finger protein 271 |
| A0A3N0XQY8_ANAGA | Zinc finger protein 271 |
| A0A8J9ZZP1 | ZNF862 protein |
| A0A8J9ZZP1_BRALA | ZNF862 protein |
