## Supplemental table S14 for "A chromosome-level, haplotype-resolved genome assembly and annotation for the Eurasian minnow (Leuciscidae: *Phoxinus phoxinus*) provide evidence of haplotype diversity"

**Table S14**: Significantly expanded genes in the *Phoxinus phoxinus* genome

| **query** | **Description** | **Preferred_name** | **PFAM Domains** |
|---|---|---|---|
| g3429.t1 | Immunoglobulin C-Typ | TRBV25-1 | C1-set,V-set |
| g17672.t1 | L1 transposable eleme | L1TD1 | Tnp_22_dsRBD,Tnp_22_trimer,Transposase_ |
| g22617.t1 | L1 transposable eleme | L1TD1 | Tnp_22_dsRBD,Tnp_22_trimer,Transposase_ |
| g6017.t1 | LINE1 type transposas | L1TD1 | Tnp_22_dsRBD,Tnp_22_trimer,Transposase_ |
| g10326.t1 | L1 transposable eleme | L1TD1 | Tnp_22_dsRBD,Tnp_22_trimer,Transposase_ |
| g20807.t1 | Interferon-induced pro | IFI44L | MMR_HSR1,TLD |
| g9.t1 | Core histone H2A/H2B | HIST2H3A | Histone |
| g18.t1 | Core histone H2A/H2B | HIST2H3A | Histone |
| g250.t1 | Core histone H2A/H2B | HIST2H3A | Histone |
| g1160.t1 | Core histone H2A/H2B | HIST2H3A | Histone |
| g1169.t1 | Core histone H2A/H2B | HIST2H3A | Histone |
| g1186.t1 | Core histone H2A/H2B | HIST2H3A | Histone |
| g1197.t1 | Core histone H2A/H2B | HIST2H3A | Histone |
| g20871.t1 | Core histone H2A/H2B | HIST2H3A | Histone |
| g20882.t1 | Core histone H2A/H2B | HIST2H3A | Histone |
| g248.t1 | Histone H2A | HIST2H2AB | Histone,Histone_H2A_C |
| g251.t1 | Histone H2A | HIST2H2AB | Histone,Histone_H2A_C |
| g1153.t1 | Histone H2A-like | HIST2H2AB | Histone,Histone_H2A_C |
| g1164.t1 | Histone H2A-like | HIST2H2AB | Histone,Histone_H2A_C |
| g1182.t1 | Histone H2A-like | HIST2H2AB | Histone,Histone_H2A_C |
| g1189.t1 | Histone H2A-like | HIST2H2AB | Histone,Histone_H2A_C |
| g1198.t1 | Histone H2A-like | HIST2H2AB | Histone,Histone_H2A_C |
| g20870.t1 | Histone H2A | HIST2H2AB | Histone,Histone_H2A_C |
| g13670.t1 | Histone H2A-like | HIST2H2AB | Histone,Histone_H2A_C |
| g7.t1 | Histone H2B | HIST1H2BA | Histone |
| g16.t1 | Histone H2B | HIST1H2BA | Histone |
| g25.t1 | Histone H2B | HIST1H2BA | Histone |
| g1157.t1 | Histone H2B | HIST1H2BA | Histone |
| g1167.t1 | Histone H2B | HIST1H2BA | Histone |
| g1171.t1 | Histone H2B | HIST1H2BA | Histone |
| g1181.t1 | Histone H2B | HIST1H2BA | Histone |
| g1190.t1 | Histone H2B | HIST1H2BA | Histone |
| g1200.t1 | Histone H2B | HIST1H2BA | Histone |
| g20873.t1 | Histone H2B | HIST1H2BA | Histone |
| g20883.t1 | Histone H2B | HIST1H2BA | Histone |
| g15620.t1 | Histone H2B | HIST1H2BA | Histone |
| g15627.t1 | Histone H2B | HIST1H2BA | Histone |
| g15630.t1 | Histone H2B | HIST1H2BA | Histone |
| g15635.t1 | Histone H2B | HIST1H2BA | Histone |
| g15644.t1 | Histone H2B | HIST1H2BA | Histone |
| g15648.t1 | Histone H2B | HIST1H2BA | Histone |
| g15656.t1 | Histone H2B | HIST1H2BA | Histone |
| g15660.t1 | Histone H2B | HIST1H2BA | Histone |

| | | | |
|---|---|---|---|
| g15704.t1 | Histone H2B | HIST1H2BA | Histone |
| g15708.t1 | Histone H2B | HIST1H2BA | Histone |
| g15713.t1 | Histone H2B | HIST1H2BA | Histone |
| g15722.t1 | Histone H2B | HIST1H2BA | Histone |
| g17066.t1 | Histone H3 | H3F3A | Histone |
| g15623.t1 | histone H3 | H3 | Histone |
| g15646.t1 | histone H3 | H3 | Histone |
| g15715.t1 | histone H3 | H3 | Histone |
| g15724.t1 | histone H3 | H3 | Histone |
| g5.t1 | Histone H2A-like | H2A | Histone,Histone_H2A_C |
| g13.t1 | Histone H2A-like | H2A | Histone,Histone_H2A_C |
| g22.t1 | Histone H2A-like | H2A | Histone,Histone_H2A_C |
| g20881.t1 | Histone H2A-like | H2A | Histone,Histone_H2A_C |
| g15641.t1 | Histone H2A-like | H2A | Histone,Histone_H2A_C |
| g15652.t1 | Histone H2A-like | H2A | Histone,Histone_H2A_C |
| g15716.t1 | Histone H2A-like | H2A | Histone,Histone_H2A_C |
| g15725.t1 | Histone H2A-like | H2A | Histone,Histone_H2A_C |
| g6723.t1 | Fucosyltransferase 9 (a | fut9b | Glyco_tran_10_N,Glyco_transf_10 |
| g6726.t1 | Fucosyltransferase 9 (a | fut9b | Glyco_tran_10_N,Glyco_transf_10 |
| g6728.t1 | Fucosyltransferase 9 (a | fut9b | Glyco_tran_10_N,Glyco_transf_10 |
| g6730.t1 | Fucosyltransferase 9 (a | fut9b | Glyco_tran_10_N,Glyco_transf_10 |
| g16455.t1 | cysteine-type endopep | CSTA | Cystatin |
| g10.t1 | Centromere kinetocho | - | CENP-T_C,Histone |
| g17.t1 | Centromere kinetocho | - | CENP-T_C,Histone |
| g169.t1 | Centromere kinetocho | - | CENP-T_C,Histone |
| g1161.t1 | Centromere kinetocho | - | CENP-T_C,Histone |
| g1183.t1 | Centromere kinetocho | - | CENP-T_C,Histone |
| g1191.t1 | Centromere kinetocho | - | CENP-T_C,Histone |
| g1201.t1 | Centromere kinetocho | - | CENP-T_C,Histone |
| g15624.t1 | Centromere kinetocho | - | CENP-T_C,Histone |
| g15628.t1 | Centromere kinetocho | - | CENP-T_C,Histone |
| g15632.t1 | Centromere kinetocho | - | CENP-T_C,Histone |
| g15636.t1 | Centromere kinetocho | - | CENP-T_C,Histone |
| g15643.t1 | Centromere kinetocho | - | CENP-T_C,Histone |
| g15651.t1 | Centromere kinetocho | - | CENP-T_C,Histone |
| g15657.t1 | Centromere kinetocho | - | CENP-T_C,Histone |
| g15661.t1 | Centromere kinetocho | - | CENP-T_C,Histone |
| g15705.t1 | Centromere kinetocho | - | CENP-T_C,Histone |
| g15710.t1 | Centromere kinetocho | - | CENP-T_C,Histone |
| g15720.t1 | Centromere kinetocho | - | CENP-T_C,Histone |
| g22571.t1 | Beta/gamma crystallin | - | Crystall |
| g22573.t1 | Beta/gamma crystallin | - | Crystall |
| g22595.t1 | Beta/gamma crystallin | - | Crystall |
| g22598.t1 | Beta/gamma crystallin | - | Crystall |
| g8166.t1 | Grass carp reovirus (G | - | PARP |
| g8177.t1 | Grass carp reovirus (G | - | PARP |

| | | | |
|---|---|---|---|
| g8179.t1 | Grass carp reovirus (G( | - | PARP |
| g392.t1 | DDE superfamily endo | - | DDE_3,HTH_Tnp_Tc3_2 |
| g20808.t1 | DDE superfamily endo | - | DDE_3,HTH_Tnp_Tc3_2 |
| g21983.t1 | DDE superfamily endo | - | DDE_3,HTH_Tnp_Tc3_2 |
| g15062.t1 | Transposase | - | DDE_3,HTH_23,HTH_Tnp_Tc3_2 |
| g2078.t1 | Si dkey-46m10.3 | - | - |
| g683.t1 | Secretory phospholipa | - | Lectin_C |
| g18190.t1 | Secretory phospholipa | - | Lectin_C |
| g653.t1 | Ribonuclease H protei | - | Exo_endo_phos,Exo_endo_phos_2,RVT_1 |
| g8884.t1 | Reverse transcriptase | - | RVT_1 |
| g3424.t1 | Immunoglobulin V-set | - | V-set |
| g3426.t1 | Immunoglobulin V-set | - | V-set |
| g3427.t1 | Immunoglobulin V-set | - | V-set |
| g3432.t1 | Immunoglobulin V-Tyr | - | V-set |
| g19644.t1 | Grass carp reovirus (GCRV)-induced gene | | PARP |

_22
_22
_22
_22
