## Supplemental table S15 for "A chromosome-level, haplotype-resolved genome assembly and annotation for the Eurasian minnow (Leuciscidae: *Phoxinus phoxinus*) provide evidence of haplotype diversity"

**Table S15:** Significantly contracted genes in the *Phoxinus phoxinus*

| query | Description | Preferred_name | Pfam Domains |
|---|---|---|---|
| g17702.t1 | NACHT, LRR and PYD domains-containing protein | NLRP12 | FISNA,LRR_6,NACHT,PRY, |
| g17724.t1 | NACHT, LRR and PYD domains-containing protein | NLRP12 | FISNA,LRR_6,NACHT,PRY, |
| g20897.t1 | NACHT, LRR and PYD domains-containing protein | NLRP12 | FISNA,LRR_6,NACHT,PRY, |
| g17832.t1 | NACHT, LRR and PYD domains-containing protein | NLRP12 | FISNA,LRR_6,NACHT,PRY, |
| g19990.t2 | NLRP3 inflammasome complex assembly | NLRP3 | FISNA,LRR_6,NACHT,PRY, |
| g20006.t1 | NACHT, LRR and PYD domains-containing protein | NLRP12 | FISNA,LRR_6,NACHT,PRY, |
| g20083.t1 | NLRP3 inflammasome complex assembly | NLRP3 | FISNA,LRR_6,NACHT,PRY, |
| g20109.t1 | NACHT, LRR and PYD domains-containing protein | NLRP12 | FISNA,LRR_6,NACHT,PRY, |
| g20560.t1 | NACHT, LRR and PYD domains-containing protein | NLRP12 | FISNA,LRR_6,NACHT,PRY, |
| g11602.t1 | NLRP3 inflammasome complex assembly | NLRP3 | FISNA,LRR_6,NACHT,PRY, |
| g17800.t1 | EGF-like module-containing mucin-like hormone | CD97 | 7tm_2,EGF_CA,GAIN,GPS |
| g22584.t1 | EGF-like module-containing mucin-like hormone | CD97 | 7tm_2,EGF_CA,GAIN,GPS |
| g18880.t1 | protein dimerization activity | - | DUF659,Dimer_Tnp_hAT, |
| g10418.t1 | GTPase IMAP family member 8-like | GIMAP7 | AIG1 |
| g17746.t1 | Immunoglobulin C-2 Type | - | Ig_2,Ig_3,V-set |
| g17749.t1 | B-cell receptor CD22-like | - | Ig_2,Ig_3,V-set |
| g17758.t1 | B-cell receptor CD22-like isoform X1 | - | Ig_2,Ig_3,V-set |
| g17762.t1 | B-cell receptor CD22-like isoform X1 | - | Ig_2,Ig_3,V-set |
| g21727.t1 | B-cell receptor CD22-like isoform X1 | - | Ig_2,Ig_3,V-set |
| g7563.t1 | Belongs to the G-protein coupled receptor 1 family | TAAR2 | 7tm_1 |
| g20836.t1 | - | - | HTH_psq,SET |
| g4150.t1 | SET (Su(var)3-9, Enhancer-of-zeste, Trithorax) domain | - | SET |
| g18764.t2 | ribonuclease inhibitor activity | RNH1 | LRR_6 |
| g22476.t1 | PRY | NLRP12 | FISNA,LRR_6,NACHT,PRY, |
| g2323.t1 | NACHT, LRR and PYD domains-containing protein | NLRP12 | FISNA,LRR_6,NACHT,PRY, |
| g16352.t1 | NLRP3 inflammasome complex assembly | NLRP3 | FISNA,LRR_6,NACHT,PRY, |
| g10422.t1 | GTPase IMAP family member 8-like | - | AIG1 |
| g9676.t1 | GTPase IMAP family member 8-like | - | AIG1 |
| g9681.t1 | GTPase IMAP family member 8-like | - | AIG1 |
| g15663.t1 | piggyBac transposable | - | DDE_Tnp_1_7 |
| g18818.t1 | AIG1 family | - | AIG1 |

| | | | |
|---|---|---|---|
| g22014.t1 | Extracellular calcium-sensing receptor-like | - | 7tm_3,ANF_receptor,NC |
| g22016.t1 | Extracellular calcium-sensing receptor-like | - | 7tm_3,ANF_receptor,NC |
| g22018.t1 | Extracellular calcium-sensing receptor-like | - | 7tm_3,ANF_receptor,NC |
| g22020.t1 | Extracellular calcium-sensing receptor-like | - | 7tm_3,ANF_receptor,NC |
| g22023.t1 | Extracellular calcium-sensing receptor-like | - | 7tm_3,ANF_receptor,NC |
| g4885.t1 | - | - | DUF4769,SAM_1 |
| g11663.t1 | - | - | DUF4769,SAM_1 |
| g1115.t1 | Natterin-like protein | - | Aerolysin,ETX_MTX2,Jaca |
| g13529.t2 | Cadherin cytoplasmic C-terminal | - | Cadherin,Cadherin_2,Cad |
| g15515.t1 | Belongs to the TRAFAC class TrmE-Era-EngA-EngB-Septin- | - | AIG1,MMR_HSR1,PRY,SP |
| g11967.t1 | Belongs to the MHC class I family | - | C1-set,MHC_I |
| g6096.t2 | class II histocompatibility antigen | HLA-DPB1 | C1-set,MHC_II_beta |
| g20931.t1 | mannose binding | CLEC17A | Lectin_C |
| g20933.t2 | mannose binding | CD209 | Lectin_C |
| g4349.t1 | mannose binding | CLEC17A | Lectin_C |
| g19541.t1 | Immunoglobulin V-Type | - | V-set |
| g19553.t1 | Immunoglobulin V-Type | - | V-set |
| g21992.t1 | Olfactory receptor C family | - | 7tm_3,ANF_receptor,NC |
| g22000.t1 | Olfactory receptor C family | - | 7tm_3,ANF_receptor,NC |
| g22002.t1 | Olfactory receptor C family | - | 7tm_3,ANF_receptor,NC |
| g618.t1 | Intelectin-1a-like | ITLN1 | Fibrinogen_C |
| g8838.t1 | Four-disulfide core domains | - | WAP |
| g4998.t1 | Neurexin | NRXN1 | EGF,Laminin_G_2,Syndec |
| g12792.t1 | Neurexin | NRXN1 | EGF,Laminin_G_2,Syndec |
| g18750.t1 | class II histocompatibility antigen | - | C1-set,MHC_II_alpha |
| g6100.t1 | class II histocompatibility antigen | - | C1-set,MHC_II_alpha |
| g10643.t1 | Low affinity immunoglobulin gamma Fc region receptor | - | Ig_2,Ig_3,ig |
| g17705.t1 | Zinc finger protein 544 | ZNF544 | KRAB,zf-C2H2 |
| g17713.t1 | Zinc finger protein 544 | ZNF544 | KRAB,zf-C2H2 |
| g621.t1 | Potassium voltage-gated channel, Shaw-related | KCNC1 | BTB_2,Ion_trans |
| g6345.t1 | Potassium voltage-gated channel, Shaw-related | KCNC4 | BTB_2,Ion_trans,Potassiu |
| g20102.t2 | Tripartite motif containing 47 | - | PRY,SPRY,zf-B_box,zf- |
| g20105.t1 | Tripartite motif containing 47 | - | PRY,SPRY,zf-B_box,zf- |

| | | | |
|---|---|---|---|
| g17180.t1 | c-X-C motif | CXCL11 | IL8 |
| g15911.t1 | Mucin-2-like | MUC2 | C8,Cys_knot,F5_F8_type |
| g6883.t2 | ryanodine receptor | RYR3 | RR_TM4- |
